# Genomic codes governing enhancer RNA fate

**DOI:** 10.1101/2024.10.31.620483

**Authors:** Chi Wai Yip, Callum Parr, Hazuki Takahashi, Kayoko Yasuzawa, Matthew Valentine, Hiromi Nishiyori-Sueki, Camilla Ugolini, Valeria Ranzani, Mitsuyoshi Murata, Masaki Kato, Wenjing Kang, Giulia Fois, Wing Hin Yip, Youtaro Shibayama, Andre Darah Sim, Ying Chen, Xufeng Shu, Jonathan Moody, Ramzan Umarov, Manli Yang, Alice Lambolez, Jen-Chien Chang, Luca Pandolfini, Tsugumi Kawashima, Michihira Tagami, Tomoe Nobusada, Tsukasa Kouno, Carlos Alfonso-Gonzalez, Rodrigo Pracana, Raquel Sofia Silva, Roberto Albanese, Francesco Dossena, Nejc Haberman, Kokoro Ozaki, Takeya Kasukawa, Boris Lenhard, Martin Frith, Beatrice Bodega, Francesco Nicassio, Lorenzo Calviello, Magda Bienko, Ivano Legnini, Valérie Hilgers, Stefano Gustincich, Jonathan Göke, Charles-Henri Lecellier, Jay W. Shin, Chung-Chau Hon, Piero Carninci

**Author notes:** Correspondence: Chi Wai Yip, Chung-Chau Hon, Piero Carninci.

## Abstract

Long-read sequencing has transformed transcriptome profiling, yet capturing full-length, non-polyadenylated transcripts like enhancer RNAs (eRNAs) remains challenging. Here, we introduce CFC-seq, combining cap-trapping and *in vitro* poly(A)-tailing to sequence poly(A) and non-poly(A) RNAs with precise transcription start site. Paired with our assembler, SALA, we identified 39,425 novel transcriptional units, including ∼24,000 eRNAs. Our data reveal a distinct genomic code governing eRNA fate dictated by core promoter architecture. CpG-island enhancers show high chromatin connectivity but yield short, exosome-sensitive RNAs. Conversely, TATA-box enhancers systematically co-opt LTR retrotransposons to inherit structural motifs that produce long, stable, and spliced RNAs. Mechanistically, the pioneer factor NF-Y activates these viral elements to license transcription, balanced by TEAD4 activity across a dual-gear regulatory axis. Finally, non-poly(A) eRNAs terminate via exosome-associated processing at structural-depleted cleavage zones. This comprehensive annotation links enhancer sequence architecture to RNA fate, providing a new transformative framework for decoding the functional human genome.

**Highlights:**

- Expanded genomic architecture: CFC-seq unmasks a hidden layer of human transcriptome, identifying 39,425 novel transcriptional units with high-confidence TSS support, including ∼24,000 eRNAs.
- TSS-first assembler: We introduce SALA, a specialized long-read assembler that prioritizes authentic 5’ Cap-trapped ends to accurately reconstruct the TSS-resolved transcript models.
- Genomic code of eRNA fate: CGI enhancers drive short and exosome-sensitive transcripts associated with repressive H3K27me3 mark and high chromatin connectivity. TATA-box enhancers produce cell-type-specific, long, stable, and frequently spliced eRNAs.
- Evolutionary co-option of retrotransposons: A major fraction of TATA-box eRNAs originate from LTR retrotransposons, providing a direct mechanism for integration of viral elements into the human regulatory landscape.
- A dual-gear pioneering axis: The pioneer factor NF-Y activates unprimed LTR-TATA enhancers to license transcription independent of histone acetylation cascades, operating in parallel with TEAD4-mediated activation.
- Structural determinants of eRNA termination: Non-poly(A) eRNA TES features a secondary structure depletion zone that coordinates pol II termination and calibrates exosome-mediated turnover.

## Introduction

Pervasive transcription of the human genome yields primary transcripts covering 75% of the genome, yet only 2% contributes to protein coding.^1^ While long non-coding RNAs (lncRNAs) represent the major product of such transcription output,^2^ existing annotations predominantly rely on short-read RNA-seq assemblies,^3–6^ which frequently struggles to resolve full-length architectures. Long-read transcriptomics has emerged as a powerful alternative,^7,8^ but its application remains largely restricted to characterizing alternative isoforms at previously annotated loci rather than discovering unannotated loci.^9–13^ Because lncRNAs are typically expressed at low levels and frequently overlap with other transcriptional units, capturing their precise transcription start sites (TSSs) via Cap-trapping is essential to verify autonomous, independent initiation. Furthermore, standard long-read protocols dependent on poly(A) selection inherently discard non-polyadenylated transcripts. A tailored full-length sequencing framework is therefore required to comprehensively map the true boundaries of the non-coding genome.

Among the candidate *cis* regulatory elements (cCREs) in the human genome, over 95% exhibit enhancer-like features,^14^ suggesting that enhancer-derived ncRNAs (eRNAs) represent the majority of lncRNAs. While enhancer activity is intensively investigated,^15,16^ eRNAs themselves exert critical functional roles.^17–21^ However, defining accurate enhancers remains challenging due to overlapping definitions with canonical promoters^22^ and the multifaceted nature of enhancer identification.^23^ When identifying enhancers via their acts of transcription, distinguishing genuine initiation sites from technical artifacts is imperative.^24,25^ Consequently, a long-read sequencing strategy capable of simultaneously resolving high-confidence 5’ ends and eRNA structures is essential to map these elusive regulatory transcripts.

Classical eRNAs are described as short, non-polyadenylated, unspliced, bi-directionally transcribed molecules.^26–28^ Conversely, longer, polyadenylated, spliced and unidirectional eRNAs have also been documented.^29–31^ Owing to nuclear exosome-mediated degradation,^32^ many eRNAs exhibit half-lives of only a few minutes.^28^ Mechanistically, how underlying core DNA sequence architectures dictate these highly divergent processing pathways, lifetimes and structural fates remains completely unknown.

To overcome current limitations in annotating the non-coding transcriptome, we developed Cap-trap full-length cDNA sequencing (CFC-seq), a method combining Cap-trapping with *in vitro* poly(A)-tailing on the Oxford Nanopore platform. Applying CFC-seq to human cellular differentiation models: iPSC-NSC-Neuron and monocyte-to-macrophage (**Fig. 1a**), we simultaneously mapped precise TSSs, transcribed *cis*-regulatory elements (tCREs) and RNA architectures. Using our specialized transcript assembler, SALA, we identified 39,425 unannotated transcriptional units including ∼24,000 eRNAs. This high resolution transcriptome uncovers a fundamental genomic code where core promoter sequence motifs dictate transcriptional fate: CpG island (CGI) enhancers generate short, non-poly(A), GC-rich and exosome-sensitive transcripts, whereas TATA-box enhancers leverage the evolutionary co-option of long terminal repeats (LTR) retrotransposons to produce long, stable, spliced and poly(A) RNA (**Fig. 1b-d**). Crucially, we demonstrate that LTR-TATA enhancers are largely regulated by pioneer factors, capable of inducing transcription without H3K27ac deposition (**Fig. 1e**). Together, our study provides a definitive framework for understanding how the heterogeneity of enhancer structures governs the lifecycle of the non-coding genome.

**Figure 1.**
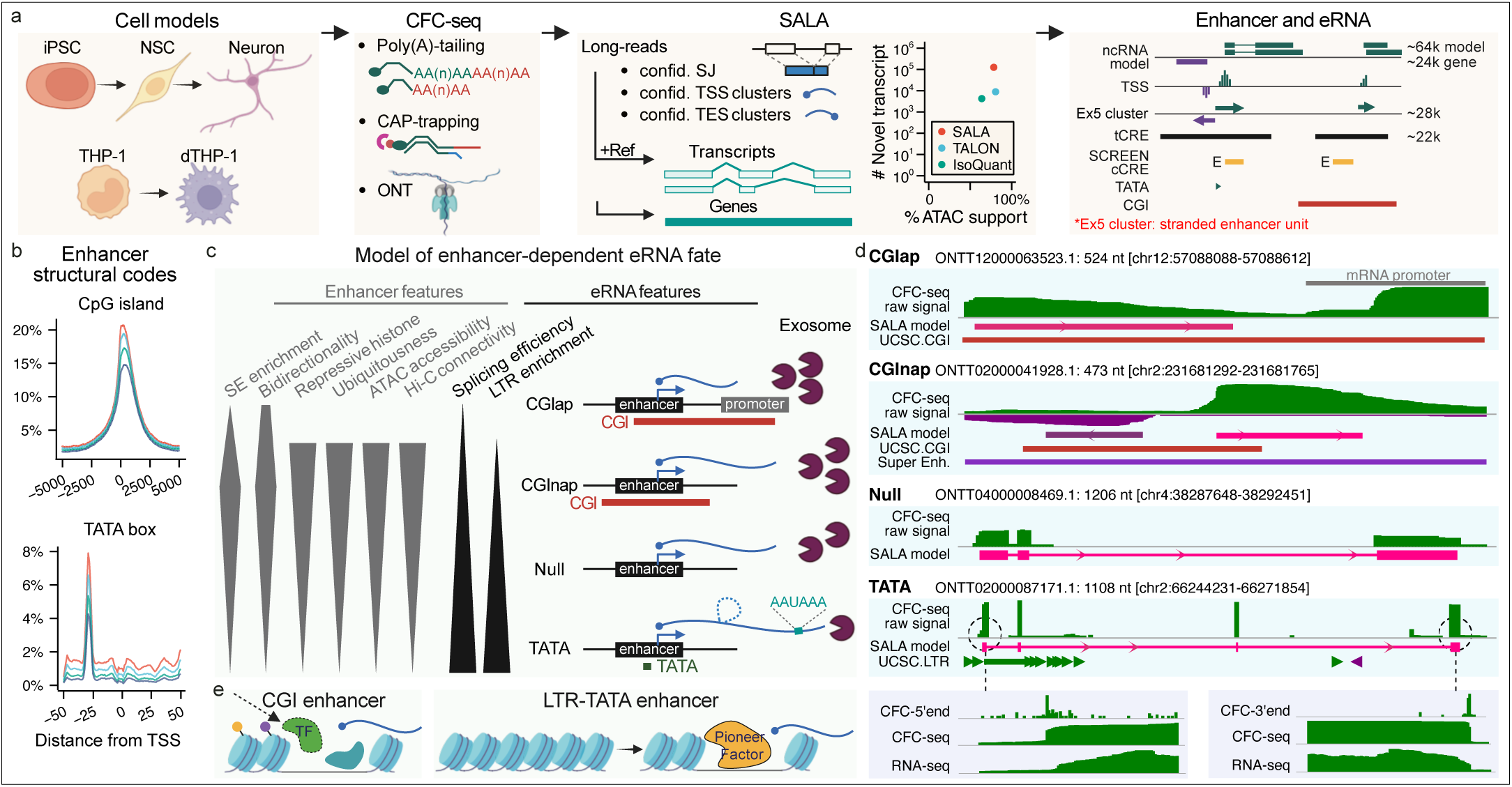
CFC-seq and SALA assembly reveal a structural code for eRNA fate. **a**, Overview of the CFC-seq, SALA assembly and enhancer annotation. Cap-trap full-length cDNA sequencing (CFC-seq) utilizes *in vitro* poly(A)-tailing and cap-trapping to capture diverse transcriptomes, including non-polyadenylated RNAs, across human differentiation models (Neurons and THP-1). The SALA assembler integrates CFC-seq long-reads with high-confidence splice junctions (SJ), transcription start site (TSS), and transcription end site (TES) clusters to generate precise transcript models. Comparison of SALA against standard assemblers (TALON, IsoQuant) reveals SALA’s superior sensitivity in capturing biologically supported novel transcripts. Annotation of enhancers and eRNAs reveals ∼24,000 eRNAs. Extended 5’ end clusters (Ex5 clusters) represents stranded enhancer units in defining CpG island enhancer proximal to promoter (CGIap), CGI not proximal to promoter (CGInap), TATA-box enhancer (TATA), and Null. **b**, Density plots showing the enrichment of CGIs and TATA-box motifs relative to the TSS of eRNAs. **c**, Schematic illustrating how enhancer structural features dictate RNA fate, including transcript length, splicing efficiency, polyadenylation and stability. **d**, Representative and simplified genomic tracks showing raw CFC-seq signal and SALA models across four eRNA classes. Transcript IDs refer to the eRNAs transcribed from plus strand. Bottom panels highlight the precise 5′ and 3′ end enrichment of CFC-seq compared to standard RNA-seq. **e**, Transcription activation from CGI enhancer and LTR-TATA enhancer. CGI enhancer is generally primed and ready to receive histone acetylation signal, while LTR-TATA enhancer is remodeled by NY-F and produce RNA without requiring histone acetylation.

## Results

### CFC-seq with comprehensive TSS coverage and non-poly(A) RNA capture

To simultaneously capture precise TSSs and RNA architectures of both polyadenylated and non-polyadenylated transcripts, we developed CFC-seq (**Ext_Fig. 1a**). Applying CFC-seq to two differentiation models yielded 236 million mappable long reads (**Ext_Fig. 1b-c**). Benchmarking against existing long-read protocols confirmed that CFC-seq achieves superior 5′ end completeness at validated promoters and chromatin-open regions (**Ext_Fig. 1d**). Furthermore, while integration of *in vitro* poly(A)-tailing (PAT) increased the detection of known non-poly(A) genes (**Ext_Fig. 1e**), capturing of non-coding non-poly(A) RNA species largely expanded (**Ext_Fig. 1f-g**). To evaluate potential artifacts introduced by PAT, we identified PAT-specific 3’ ends from the mRNA fraction while ∼2% of them revealed as nascent RNA intermediates (**Ext_Fig. 1h-j**). Finally, PAT-specific fraction uniquely recovered 5,964 ncRNA gene models compared to only 935 in non-PAT controls (**Ext_Fig. 1k**).

To quantitatively assess polyadenylation events, we trained a sequence-based Random Forest classifier on orthogonal FLAM-seq datasets,^33,34^ where transcript ends were defined by the presence of non-templated poly(A). This poly(A) score revealed high performance with AUC 0.93 from the THP-1 non-PAT and PAT-specific 3’ ends (**Ext_Fig. 1l**). Thresholds for defining poly(A)-positive and -negative were extracted from the ROC curve with 95% confidence. (**Table S1**).

### CFC-seq identifies genuine TSS to indicate independent transcription

A hallmark of genuine TSSs is the presence of an unencoded, non-template G residue immediately upstream of the mapped read boundary,^35^ a feature observed in approximately 80% of qualified CFC-seq reads (**Ext_Fig. 2a**). Combining the CFC-seq libraries across our cellular differentiation time courses, we identified 159,116 high-confidence TSS clusters, which were extended and merged into 73,328 tCREs (**Fig. 2a**, **Table S2**). Cross-referencing their genomic coordinates against SCREEN cCREs,^14^ 32,416 tCREs were classified as enhancer-like, which displayed cell-type specific expression profiles (**Fig. 2b**).

**Figure 2.**
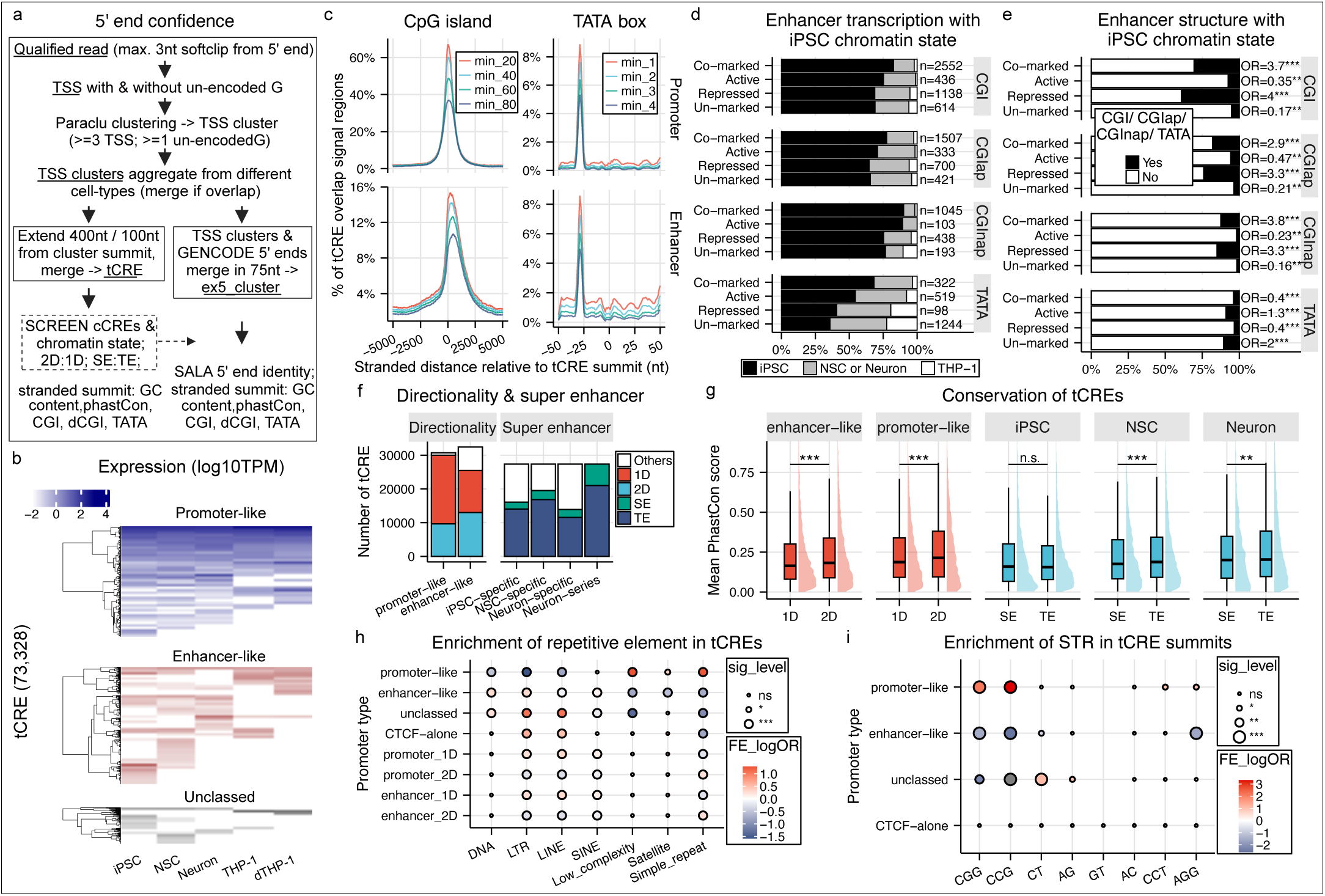
Identification of genuine TSSs and tCREs from CFC-seq. **a**, Flowchart of the SCAFE pipeline for identifying genuine TSS clusters, tCREs, and ex5_clusters from CFC-seq reads. **b**, A total of 73,328 tCREs were identified across five cell types and classified as promoter-like, enhancer-like, or unclassed based on annotation from SCREEN cCRE. Normalized tag count of 10% of these tCRE were visualized in the heatmap**c**, Distribution of CpG island (CGI) density and TATA box motif score across major-strand tCREs, stratified by promoter-typing. Positions are relative to tCRE summits. **d**, Genomic coordinates of enhancer-like tCREs (major strand only) were intersected with iPSC H3K27Ac and H3K27Me3 CUT&Tag data and grouped into Co-marked (Both signals), Active (27Ac alone), Repressed (27Me3 alone), and Un-marked (Neither). TSS counts ≥1 were considered positive in expression. Top panel: percentage of expressed tCREs per enhancer state in CGI-positive (TATA-negative) class; second to bottom panels: CGIap-positive (CGI-positive that are adjacent to promoter), dCGInap-positive (dCGI-positive that are not adjacent to promoter) and TATA-positive (CGI-negative). **e**, Enrichment between enhancer structures (CGI, CGInap, dCGInap & TATA) and chromatin states. Fisher’s exact test was used for enrichment (OR: odds ratio). **f**, Number of tCREs classified as unidirectional (1D), bidirectional (2D), super enhancer (SE), and typical enhancer (TE), based on **Extended Data Figure 3a-c**. **g**, Sequence conservation (17-way PhastCons) for 1D vs. 2D and SE vs. TE tCREs, using the averaged conservation score ±500 nt from tCRE summits. **h**, Enrichment of repetitive elements across promoter types and directionality tested by two-tailed Fisher’s exact test. Repetitive element overlap is as in **Extended Data Figure 3m**, where the the ones with the longest overlap were selected from “multiple” as representatives. **i**, Enrichment of colocalization of TSS and short tandem repeats (STR) across different promoter types tested by two-tailed Fisher’s exact test. Unless specified, statistical tests were two-sided Wilcoxon; p < 0.05 (*), p < 0.01 (**), p < 0.001 (***), n.s. = not significant.

Integrating the Neuron-series tCREs with matching single-cell ATAC-seq profiles confirmed that the vast majority correspond to regions of accessible chromatin (**Ext_Fig. 2b**). Notably, 36% of the ATAC-validated unclassified tCREs were unique to NSC, a cell type underrepresented in the SCREEN database (**Ext_Fig. 2c**). Chromatin-state mapping via ChromHMM independently verified that the remaining unclassified tCREs overlap promoter or enhancer regions (52%, **Ext_Fig. 2d**), with a minor fraction derived from 3’UTRs, which was previously reported as capped 3’UTR-derived RNAs.^36^ Collectively, this multi-omic cross-validation provides a robust framework to anchor long-read transcripts to TSS clusters, tCREs and promoter-type.

### Epigenetic divergence distinguishes CGI and TATA-box enhancers

To dissect the features driving non-coding initiation, we mapped the strand-specific summits of the major transcriptional strands of our tCREs relative to core genomic landmarks. Notably, both promoter-like and enhancer-like tCREs showed enrichment of TATA-boxes, initiator motifs, CGIs, high GC content and sequence conservation (**Ext_Fig. 2e**), demonstrating their capacity to initiate transcription. While promoters align symmetrically with CGIs, enhancer-like tCREs exhibited a spatial asymmetry, with CGIs enriched downstream of their TSSs (**Fig. 2c**). Interestingly, this spatial bias was restricted to promoter-adjacent enhancers (Enhancer-ap), whereas distal enhancer (Enhancer-nap) maintained a symmetrical promoter-like alignment centered over the CGI peak (**Ext_Fig. 2f**). Conversely, the antisense minor strand of CGIap enhancer exhibited a mirrored, opposite orientation (**Ext_Fig. 2f&g**). This demonstrates that a directional TSS-upstream-of-CGI configuration is a hallmark of eRNA initiation at promoter-proximal regulatory hubs.

To determine whether these genomic architectures dictate different chromatin environments, we stratified enhancer-like tCREs into CGI (CGIap/CGInap), TATA-box, or unclassified (Null) (**Ext_Fig. 2h&i**), and profiled them against active (H3K27ac) and repressive (H3K27me3) histone modifications. Expression profiling revealed that CGI enhancers are transcribed more ubiquitously across our cell types than TATA-box enhancers (**Fig. 2d**). Crucially, repressed or co-marked (both H3K27ac and H3K27me3) chromatin states were significantly enriched at CGI-associated enhancers (**Fig. 2e**), coincident with a previous study.^37^ In contrast, TATA-box enhancers were typically depleted of H3K27me3, associating instead with either active chromatin or regions devoid of both marks (**Fig. 2e**). This epigenetic divergence was consistently maintained across iPSC, NSC and Neuron (**Ext_Fig. 2i**), revealing that core sequence elements hardwire distinct chromatin states of enhancers.

### Bidirectionality and CGIs drive enhancer sequence conservation

Bidirectional transcription is a hallmark of enhancers.^26^ Utilizing strand-specific 5’ cap signals, we systematically segregated our tCREs into bidirectional (2D) and unidirectional (1D) cohorts (**Fig. 2f, Ext_Fig. 3a**). We further integrated chromatin-state profiles to stratify these elements into super-enhancers (SEs) and typical enhancers (TE, **Fig. 2f, Ext_Fig. 3b-c**). While prior research indicates that SEs are generally more conserved than TEs,^38^ our analysis of transcriptionally active enhancers revealed comparable baseline conservation levels between the two groups (**Fig. 2g**). Instead, we found that sequence conservation is correlated with transcriptional directionality, with 2D enhancers exhibiting significantly higher conservation (**Fig. 2g & Ext_Fig. 3d**).

We hypothesized that this conservation signature is rooted in the presence of CGIs, which are enriched in 2D tCREs (**Ext_Fig. 3e**). Further analysis revealed that 2D enhancers containing CGIs are significantly more conserved than their non-CGI counterparts (**Ext_Fig. 3f**), suggesting that the underlying CGI sequence contributes to enhancer conservation. To integrate strand-specific enhancer features of transcript models, we repeated these analyses on ex5_clusters that linked to the finalized transcript models and found consistent results as from the tCREs (**Supplementary Fig. S1 & Table S3**).

### CFC-seq identifies TSSs within repetitive genomic elements

To benchmark the fidelity of CFC-seq, we compared its performance against CAGE, which utilizes the same CAP-trapping molecular protocol. Both platforms demonstrated highly comparable accuracy in capturing genuine transcription initiation events, exhibiting strong concordance in unencoded G frequencies, TSS positioning at single-nucleotide resolution, overall tCRE identification and quantification (**Ext_Fig. 3g-j**). However, a limitation of short-read CAGE protocols is their inability to uniquely map reads with highly repetitive genomic intervals. Benefiting from extended read-length, CFC-seq demonstrated superior performance over single-end CAGE in resolving TSSs within several classes of repetitive elements (**Ext_Fig. 3k**). These repetitive-element-rich TSSs uniquely identified by CFC-seq were evolutionarily younger (**Ext_Fig. 3l**), aligning with recent findings.^39^

Stratifying the major-strand tCRE landscape revealed that 66.1% of these elements overlap transposable or repetitive sequences (**Ext_Fig. 3m**). Enhancer-like tCREs were enriched for DNA transposons, SINEs, and LTRs, which preferentially enriched in unidirectional elements (**Fig. 2h**). Conversely, promoter-like tCREs were enriched for low-complexity and simple repeats, with HipSTR intersection identified CGG/CCG expansions as the most prominent motif (**Fig. 2i**), consistent with a previous study.^40^ Together, these benchmarking analyses demonstrate that CFC-seq achieves CAGE-level single-nucleotide accuracy while successfully uncovering repeat-dense transcriptional landscape.

### SALA resolves thousands of novel transcriptional units from CFC-seq

To construct a high-confidence transcriptome, we processed 236 million mapped alignments through our specialized long-read assembler, SALA (**Fig. 3a**; Methods). To enforce stringent initiation-site accuracy, we restricted all the transcript models with SCAFE TSS-cluster support (**Ext_Fig. 4a-c**, Methods). This rigorous filtering strategy yielded a finalized assembly of 194,138 transcript models including 146,154 putative novel transcripts (**Fig. 3b, Table S4**).

**Figure 3.**
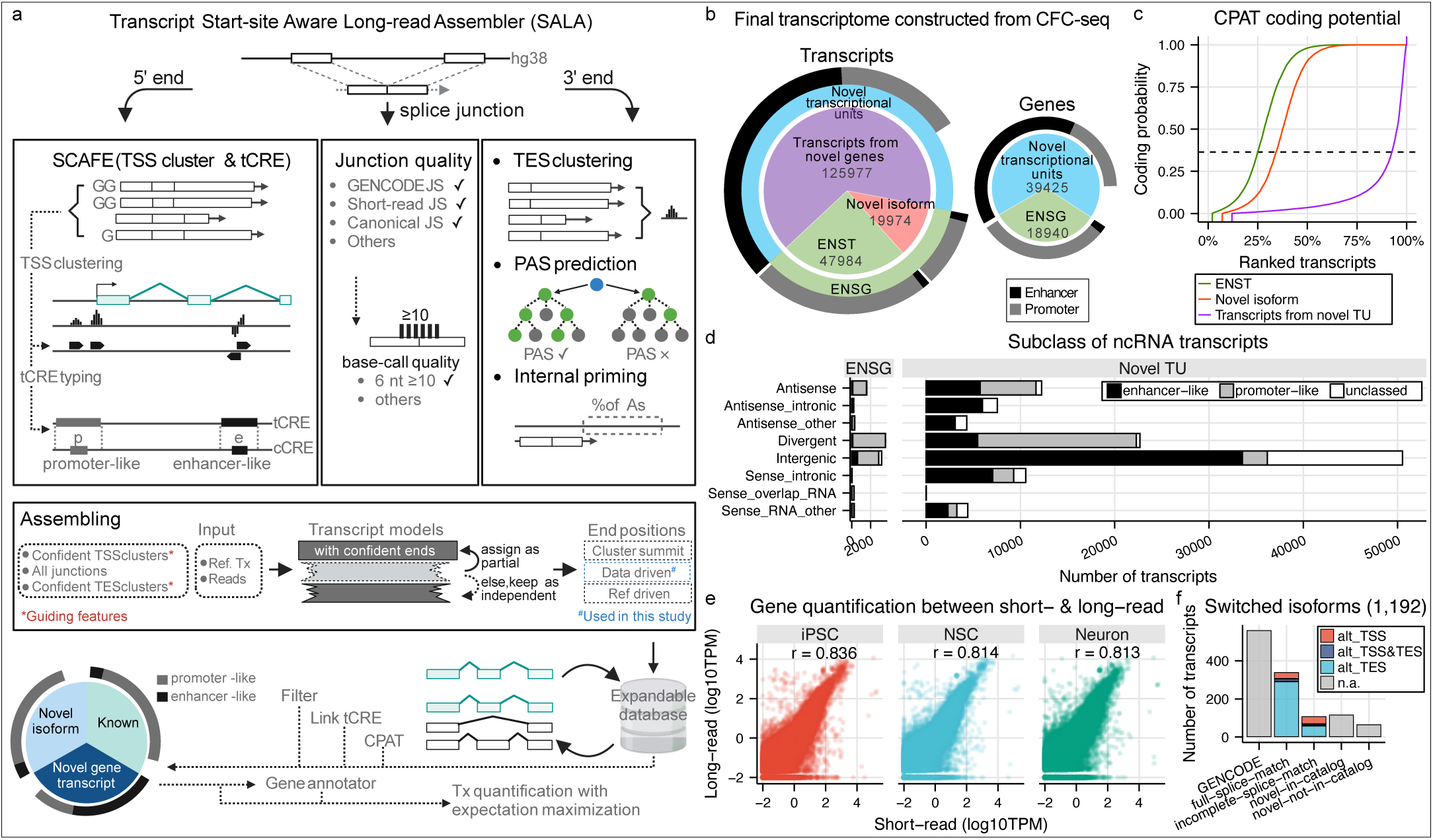
Transcript models identified by SALA. **a**, Overview of the analytical pipeline. Three transcript features—5′ end, splice junctions, and 3′ end—were extracted from all reads. TSS clusters were defined and filtered by SCAFE; splice junctions were evaluated against GENCODE (v39), matched short-read RNA-seq, and base-calling quality summarization; TES clusters were formed and filtered by read count, with additional 3′ end labels from PAS and internal priming detection. These features, along with all reads and the reference transcriptome, were used for guided transcript assembly. The resulting transcript models were annotated with tCRE-based promoter types and filtered for downstream analysis. Final quantification was performed using expectation maximization in Bambu. **b**, Number of transcript and gene models identified by SALA. Transcript models were classified into GENCODE-annotated (ENST), novel isoforms (within known genes), and transcripts from novel transcriptional units. **c**, Coding potential of transcript groups assessed using CPAT. **d**, All detected ncRNAs, including annotated lncRNAs and novel ncRNAs, were subclassified based on their location relative to protein-coding genes and pseudogenes. **e**, Comparison of gene-level expression from long- and short-read quantification using the finalized transcriptome and detectable GENCODE models, ribosomal RNAs were excluded. **f**, Classes of transcript models that exhibit significant isoform switching across iPSC, NSC, and Neuron, only isoforms from GENCODE genes were shown. Alternative (alt) TSS and TES represent novel transcript models with unannotated TSS and TES respectively.

In addition to SALA’s final transcript models, we presented the pre-filtered raw SALA assembly (**Table S5**) and an independent run of SALA default (**Table S6**) to benchmark SALA’s performance against TALON^41^ and IsoQuant^42^ utilizing identical internal priming filters across the same CFC-seq dataset (**Ext_Fig. 4a,d**). These tools show broadly comparable performance in capturing annotated GENCODE transcripts (**Ext_Fig. 4e**), while their structural handling of complex loci diverged. Unlike IsoQuant, SALA preserves nearby SJs, leading to exclusion of 3,235 ENST models while 73% of them were considered as ISM isoforms. Furthermore, by requiring precise alignment of transcript ends within TSS and transcription end site (TES) clusters, SALA effectively filtered out false-positive full-splice matches that actually utilized alternative promoters or PASs (**Ext_Fig. 4e**).

Crucially, SALA’s final assemblies identified a higher proportion of novel intergenic, antisense, and intronic transcripts than reference-centric assemblers (**Ext_Fig. 4f-g**). Despite this expanded discovery rate, the 5′ ends of SALA’s novel models exhibited precision levels (supported by ATAC-seq and SCREEN cCRE datasets) equal to or exceeding those of existing tools (**Ext_Fig. 4h**). By utilizing the superior TSS precision of Cap-trapping, SALA effectively bridges the gap between raw long-read sequencing signals and functional annotation.

### Systematic classification reveals a vast landscape of unannotated ncRNAs

Coding potential analysis revealed that over 80% of the transcripts from novel transcriptional units (TUs) are non-coding, compared to around 25% of known GENCODE transcripts (**Fig. 3c**). At the gene level, 93% of the 39,425 novel TUs were classified as non-coding, comprising 35,848 lncRNAs and 1,062 short_ncRNAs respectively (**Ext_Fig. 4i**). Only non-coding transcripts from the non-coding genes were considered as ncRNAs in the downstream analyses. We have identified 115,188 novel ncRNA transcripts and detected 6559 known lncRNA transcripts (**Ext_Fig. 4j**). These 121,747 ncRNA transcripts were then further subclassed according to the genomic location of GENCODE coding-genes and pseudogenes (**Fig. 3d**, Methods) for the downstream analyses.

### CFC-seq identified a vast repository of strictly unannotated non-coding genes

To assess whether our newly discovered loci intersect with existing non-coding databases, we cross-referenced our novel transcript models against Refseq, GENCODE (v47), FANTOM CAT^5^ and LncBook.^2^ At the transcript level, about 9% of the novel transcript models were annotated by at least one of these databases, while 55% of our novel TUs are unannotated from any of the four reference repositories (**Ext_Fig. 5a-c, Table S7**). As expected, these novel TUs uniquely observed from CFC-seq are relatively more cell-type specific and lowly expressed (**Ext_Fig. 5d**).

Next, we quantified independent short-read RNA-seq datasets of the Neuron series by kallisto.^43^ Remarkably, over 80% of our novel transcript models and more than 97% of the newly discovered gene models were detected from the short-read RNA-seq data (**Ext_Fig. 5e**), suggesting most of the novel models can be detected from an independent sequencing technique. The minor fraction of undetected models displayed a similar proportion of ncRNA subclass as their detected counterparts (**Ext_Fig. 5f**).

Expression quantification derived from the CFC-seq data (**Table S8**) showed a modest overall concordance with the short-read RNA-seq estimates (**Fig. 3e** & **Ext_Fig. 5g**). While transcript length contributed substantially to the discrepancy (**Ext_Fig. 5h**), PCR bias from the CFC-seq may influence quantification. Other causes include differences in sequencing platforms, sequencing depth, and the predictive short-read quantification algorithm.

### 16,059 novel isoforms were identified from protein-coding genes

Beyond unannotated loci, our assembly adopted a stringent requirement to identify novel isoforms mapping to annotated protein-coding genes (Methods). Over 20% of these unannotated isoforms were classified as non-coding (**Ext_Fig. 5i**). Mechanistically, while alternative TES selection drove the vast majority of these isoforms, 58% featured novel splicing configurations. Among the alternative TES isoforms, full-splice matches were predominantly polyadenylated, whereas incomplete-splice matches frequently lacked polyadenylation (**Ext_Fig. 5j**). These non-polyadenylated isoforms likely represent stable processing intermediates, with over 50% of them ended at the donor sites. Notably, canonical reference isoforms co-existed at more than 60% of these loci, validating the fidelity of our models over existing annotations.

To evaluate the functional dynamics of these variants during human neurogenesis, we performed differential isoform switching analysis across iPSC, NSC and Neuron. We identified 1,192 transcripts exhibiting significant fractional abundance shifts, while 632 of them are unannotated isoforms (**Fig. 3f** & **Table S9**). We further evaluated 266 genes with both significantly up- and down-regulated transcripts in any of the three comparisons (**Ext_Fig. 5k-i**). Crucially, these dynamic isoform-switching events target key loci linked to human neurological disorders, including *AP1S2*, *PORCN*, *PRKCZ* and *CA14* (**Figs. S2&3**).

### Enhancer transcribed ncRNAs are longer and more spliced than previously reported

To profile transcript biology across different initiation loci, we segregated our ex5_clusters (stranded genuine TSS cluster with extension) into five functional groups according to the promoter types and the coding potential of the RNA output (Methods). These resulted in ex5_clusters with mRNA outputs, promoter-derived ncRNA (p_ncRNA), enhancer-derived ncRNA (e_ncRNA), CTCF-alone ex5_clusters with ncRNA (CTCF_ncRNA) and unclassed ex5_clusters with ncRNA (other_ncRNA) (**Table S10**).

Investigating the transcript features of these classes revealed that e_ncRNAs possess a median length of 657 nt (**Fig. 4a & Ext_Fig. 6a**). This is notably longer than the historically reported range of 346-449 nt.^26,44^ To confirm this length discrepancy was not an artifact of our custom enhancer classification, we re-evaluated our data using multiple established orthogonal definitions of enhancer boundaries, including distal chromatin accessibility peaks,^45^ distal 5’ end signal derived tCREs,^46^ classical bi-directional RNA footprints,^26^ and cell-type-specific chromatin states.^38^ Across all alternative definitions, the median length of captured eRNA products remained consistently expanded, ranging from 654 to 705 nt (**Ext_Fig. 6b-c**).

**Figure 4.**
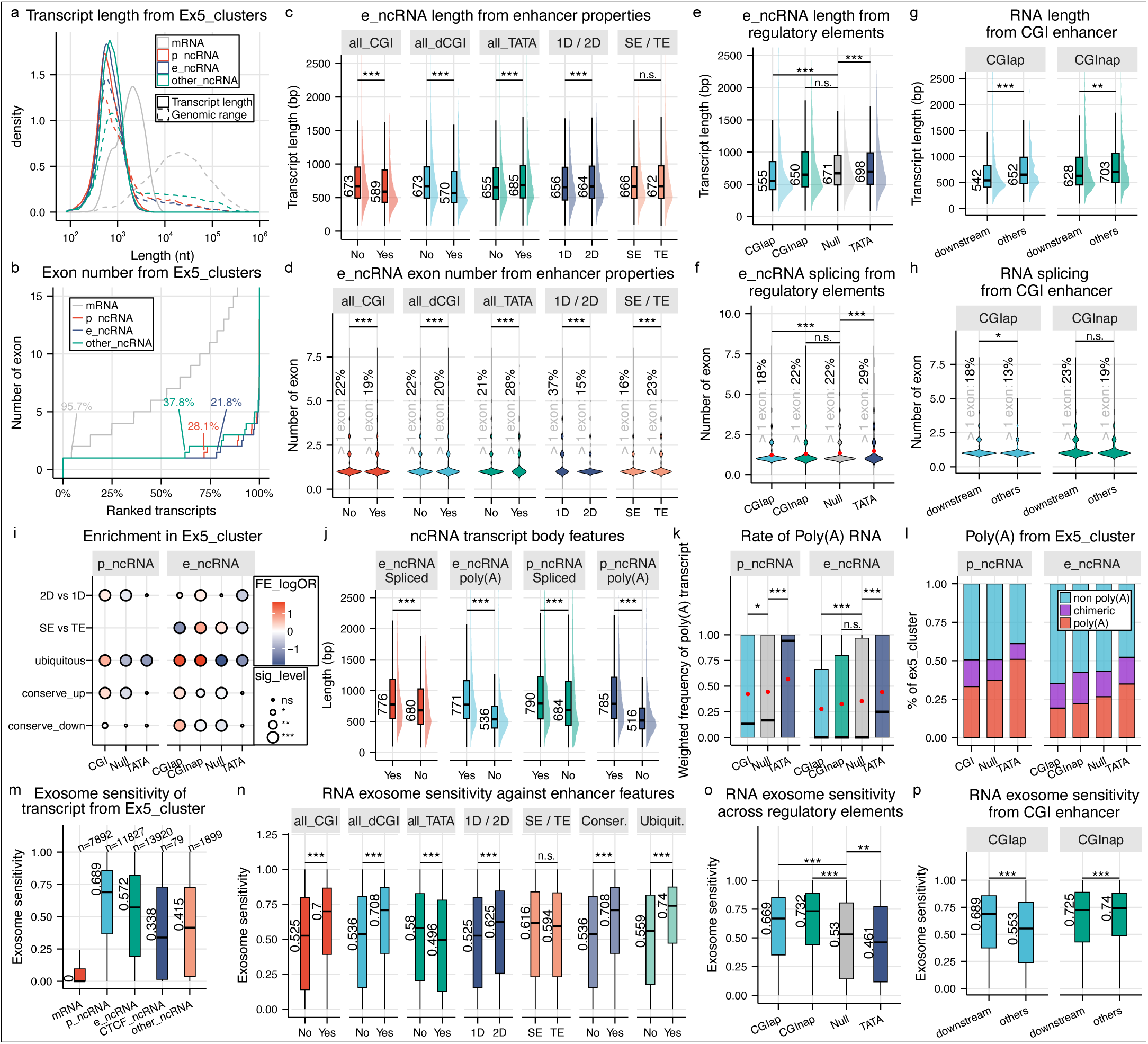
Factors that affect the transcript model of eRNAs. **a**, Transcript length and genomic range of RNA classes grouped by extended 5′ end clusters (ex5_clusters). **b**, Median number of exons for each RNA class grouped by ex5_clusters. Percentages indicate groups with more than one exon. **c**, Comparison of e_ ncRNA transcript lengths across genomic features: all_CGI (CpG island within ±500 nt of summit), downstream CpG island (dCGI, within 500 nt downstream), TATA box (motif located −35 to −11 nt upstream), directionality (unidirectional, 1D; bidirectional, 2D), and enhancer type (super enhancer, SE; typical enhancer, TE) defined from iPSC, NSC or Neuron (excluding THP-1-derived enhancers). **d**, Median number of exons of e_ncRNAs grouped by genomic features. Percentages denote the fraction with more than one exon. **e**, Transcript length of e_ncRNAs grouped by mutually exclusive features; “Null” indicates ex5_clusters lacking any of the tested features. **f**, Median number of exons of the mutually exclusive e_ncRNA groups. **g**, Transcript length of e_ncRNAs comparing with and without the presence of dCGI, among CGIap and GCInap enhancers. **h**, Median number of exons comparing with and without the presence of dCGI. **i**, Enrichment of CGI, CGIap, CGInap, Null and TATA box across genomic features in e_ ncRNA and p_ncRNA using Fisher’s exact test (two-tailed). FE_logOR denotes the natural log of odds ratio. **j**, At the transcript level, presence of poly(A) and presence of splicing were evaluated for each model, and length distributions were compared across these features. **k**, Transcript models grouped by ex5_cluster with poly(A) frequency weighted by read count. Red dots indicate the mean values. **l**, Percentage of ex5_clusters produce RNA with poly(A) (weighted poly(A) frequency > 0.9), non-poly(A) (frequency < 0.1) and chimeric. **m**, Exosome sensitivity per transcript, calculated using RNA-seq after EXOSC3 knockdown in iPSC, was grouped by ex5_cluster (weighted by read count). RNA categories include mRNA, p_ncRNA, e_ncRNA, CTCF_ncRNA, and other_ncRNA. **n** Exosome sensitivity of e_ncRNAs grouped by ex5_clusters and genomic features. Conservation was defined using 17-way phastCons scores from 500 nt upstream of the summit; clusters above the median score were classified as “Yes.” Other feature definitions as in (**c**). **o**, Exosome sensitivity of enhancer-like ex5_clusters across mutually exclusive regulatory element groups. **p**, Exosome sensitivity of RNA derived from CGIap and CGInap enhancers, with and without dCGI. Unless specified, statistical tests were two-sided Wilcoxon; p < 0.05 (*), p < 0.01 (**), p < 0.001 (***), n.s. = not significant.

Furthermore, we discovered that 21.8% of e_ncRNA loci generated transcript containing at least one splice junction (**Fig. 4b**), compared to 5% in a previous study.^26^ This splicing activity remained consistent (19–27%) across all examined enhancer definitions (**Ext_Fig. 6d**). Collectively, these data indicate that eRNAs are longer and undergo more pervasive splicing than previously recognized, likely due to our improved capture of TSS-resolved transcript models.

### Core genomic architectures correlate with eRNA length and splicing complexity

Promoter sequence architecture is known to functionally couple transcription initiation and downstream RNA splicing mechanics.^47,48^ We therefore investigated whether analogous genomic features features within enhancers, including bidirectionality, SE association and presence of TATA-box, CGI or downstream CGI (dCGI), correlate with eRNA length and processing dynamics. Interestingly, presence of CGI and dCGI strongly correlated with shorter eRNA length and reduced splicing frequency (**Fig. 4c-d**). Conversely, TATA-box-containing enhancers were associated with extended transcript length and elevated splicing activity. While bidirectional transcription and SE localization exerted only subtle effects on absolute eRNA length, both features correlated with significantly fewer splicing events (**Fig. 4d**).

To isolate the precise contributions of these regulatory motifs, we stratified enhancers into four mutually exclusive groups: promoter-adjacent CGI (CGIap) and distal orphan CGI (CGInap), unclassified (Null), and TATA-box (**Ext_Fig. 6e**). Cross-cohort comparisons confirmed that CGIap as a primary driver of short and unspliced transcripts, while TATA-boxes promote enhanced transcript length and splicing complexity (**Fig. 4e-f**), a trend also observed in p_ncRNAs (**Ext_Fig. 6f-i**).

Intriguingly, the presence of a dCGI significantly restricted the RNA length of both CGIap and CGInap transcripts without impacting splicing (**Fig. 4g-h**). This suggests that the physical proximity of downstream transcription machinery imposes a strict topographical constraint that limits eRNA elongation. Beyond transcript processing, we further observed that TATA-box enhancers are enriched for unidirectionality and mapping to typical enhancers (**Fig. 4i**). In contrast, while CGIap was depleted from SE, CGInap showed significant SE enrichment, suggesting these orphan intergenic CGIs may serve as focal points for SE assembly. Further, ubiquitous enhancers (expressed across all five cell types) are enriched for CGIs, and depleted of TATA-boxes, consistent with our previous study.^49^ Overall, these data demonstrate that enhancer sequence architectures dictate eRNA processing, while downstream structural constraints of the dCGI physically restrict their genomic reach.

### Splicing and polyadenylation drive length expansion and processing heterogeneity of ncRNA

To investigate the features governing transcript processing, four RNA classes (mRNA, p_ncRNA, e_ncRNA, and other_ncRNA) were stratified at the transcript level (**Table11**, Methods). Across both e_ncRNA and p_ncRNA, the acquisition of either a splice junction (SJ) or a PAS motif was associated with increased transcript length (**Fig. 4j**). Next, we grouped transcripts into ex5_clusters based on the weighted frequency of transcript models with polyadenylation. Notably, ncRNAs derived from CGIap enhancers showed significantly lower polyadenylation than those from TATA-box enhancers where 34.9% TATA-box enhancers versus 19.2% CGIap enhancers produce poly(A) RNAs, a trend also observed for p_ncRNAs (**Fig. 4k-l**). Beyond absolute length, we examined intra-locus length variation as a proxy for molecular processing precision. Normalized length variation was consistently higher for ncRNAs than mRNAs, peaking at loci characterized by a balanced mixture of poly(A)+ and poly(A)− species (**Ext_Fig. 6j**). This heterogeneity suggests that non-coding transcript termination and processing operate with higher stochasticity than protein-coding mRNA, with alternative TESs and variable splice junctions introducing localized plasticity.

### Chromatin-bound eRNAs and exosome sensitivity

Because eRNAs are typically unstable and transient, we performed CFC-seq on the chromatin-bound RNA fraction of iPSCs to enrich nascent transcripts. Thousands of novel ncRNA loci exclusive to the chromatin-bound fraction were identified (**Table S12**), while 6,476 of them originated from enhancer regions (**Ext_Fig. 6k**). Comparison between the total and chromatin-bound datasets revealed greater length diversity among ncRNAs than mRNAs, reflecting a prevalence of non-polyadenylated, nascent transcripts with variable TESs (**Ext_Fig. 6l**).

To evaluate the turnover dynamics of these transcripts, we integrated independent exosome knockdown RNA-seq data.^50^ Chromatin-bound ncRNAs are more sensitive to exosome-mediated degradation (**Ext_Fig. 6m**). As expected, mRNAs exhibited lower exosome sensitivity than ncRNAs (**Fig. 4m**). This susceptibility was independently supported by RBP-binding data, which showed a significant enrichment of the degradation factors EXOSC5 and XRN2 on enhancer-derived transcripts (**Ext_Fig. 6n**). Critically, exosome sensitivity varied significantly across enhancer subclasses. The RNA derived from CGI, dCGI, 2D, conserved and ubiquitously active enhancers were significantly more exosome-sensitive, where CGInap enhancers yielded the most exosome-sensitive eRNAs (**Fig. 4n-p**). Conversely, TATA-box enhancer generates significantly more stable eRNAs. A similar trend was found in p_ncRNAs (**Ext_Fig. 6o**). Collectively, these findings demonstrate that enhancer architecture influences RNA stability, by programming their susceptibility to the nuclear RNA surveillance machinery.

### Non-coding RNA exhibit reduced splicing efficiency compared to mRNAs

To evaluate the splicing activity of non-coding transcripts, we systematically profiled the confidence, recurrence and efficiency of the SJs. Cross-referencing the SJs from our finalized transcriptome against GENCODE v39, the Intropolis database,^51^ and matching short-read RNA-seq data, 88.2% of all SJs were validated with most of them supported by GENCODE (**Fig. 5a**, **Ext_Fig. 7a**). With a threshold of a summarized base-calling score ≥ 10 and a read count ≥ 3, 8.9% of these novel SJs were classified as technical confident (**Ext_Fig. 7b**). Collectively, 89.3% of all SJs observed across the final transcriptome were either verified by external databases or passed the internal quality controls. Stratifying these models at the transcript level into mRNA, p_ncRNA, e_ncRNA and other_ncRNA, revealed that approximately 90% of all spliced ncRNAs possessed entirely confident SJs (**Ext_Fig. 7c**) while the majority of ncRNA SJs utilized canonical sites (**Ext_Fig. 7d**), echoing the findings of a previous study.^31^

**Figure 5.**
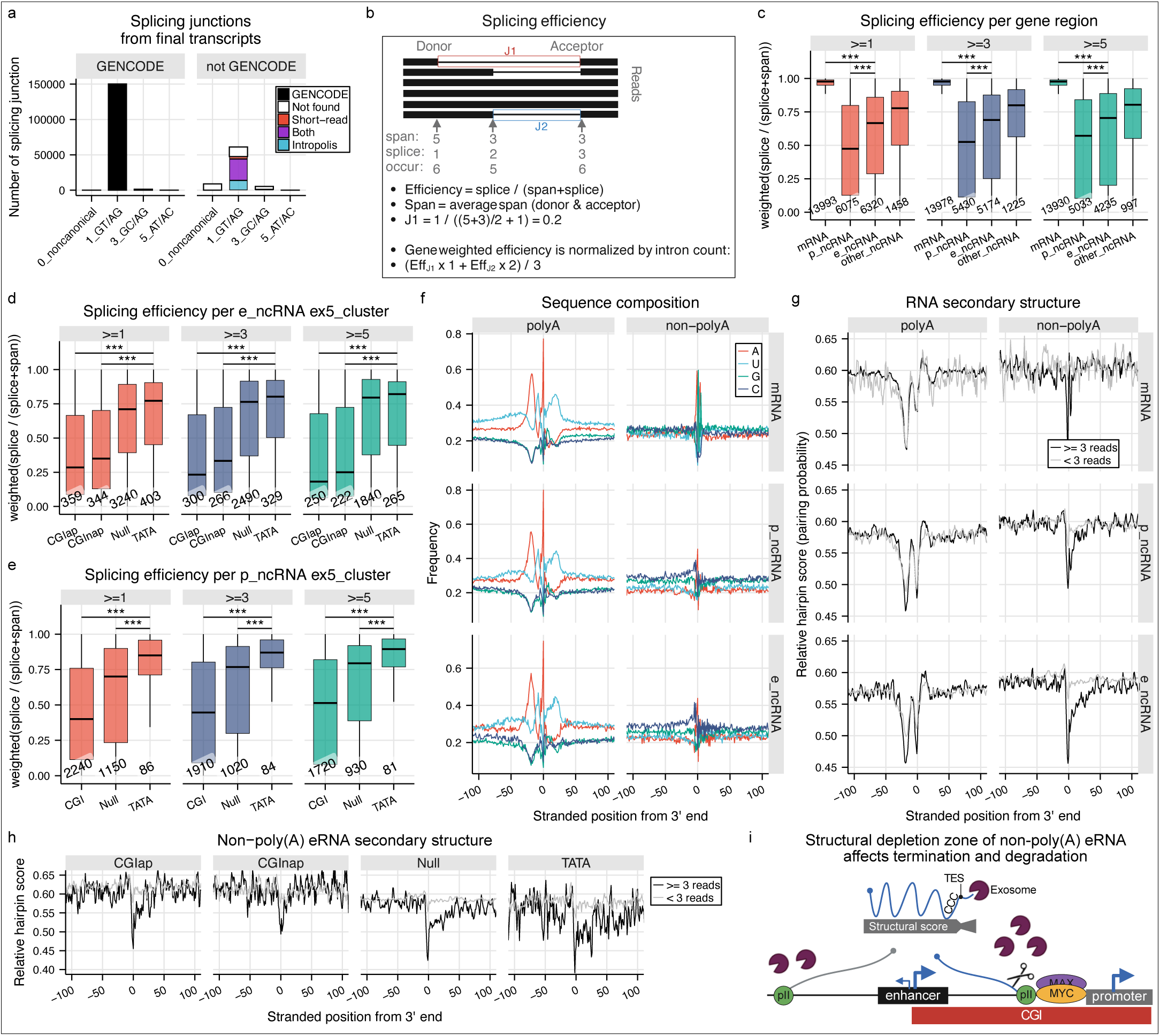
Splicing efficiency and transcription end sites of non-poly(A) ncRNAs. **a**, All splice junctions (SJs) identified in the final transcriptome. Most were within GENCODE (v39) annotation, while the majority of novel (non-GENCODE) SJs were canonical. **b**, Schematic illustrating the calculation of splicing efficiency per junction and the weighted splicing efficiency per gene. **c**, Splicing efficiency summarized at the gene level and normalized by intron count (see Methods and panel **b**). Gene models lacking splice junctions were excluded. SJs were filtered using three thresholds for junction support (≥1, ≥3, or ≥5 total reads spanning or splicing both donor and acceptor sites). **d**, Splicing efficiency of e_ncRNAs grouped by ex5_clusters and compared across regulatory elements (CGIap, CGInap, Null, TATA box). Only ex5_clusters assigned with one gene model were kept. **e**, Same analysis as in **d** for p_ncRNAs. **f**, Nucleotide composition surrounding recursive transcription end sites (TESs; ≥3 read support) of different RNA classes, stratified by poly(A) status. **g**, RNA secondary structure predictions from RNAfold using ±200 nt sequences centered on the 3′ ends of poly(A) and non-poly(A) eRNAs. Relative hairpin scores were summarized for ±100 nt around the TES. Sequences were input to RNAfold without accounting for splicing. Analyses were conducted separately for recursive (≥3 reads) and non-recursive TESs. **h**, Same analysis as panel **g**, subgrouped by regulatory elements from non-poly(A) eRNAs. **i**, Schematic illustrating structural depletion-associated non-PAS termination of RNA derived from CGIap enhancer. Unless specified, statistical tests were two-sided Wilcoxon; p < 0.05 (*), p < 0.01 (**), p < 0.001 (***), n.s. = not significant.

Although the core SJ motif of ncRNA mirrors those of protein-coding mRNA, individual transcript classes process these junctions with different efficiency.^52^ To quantify this, we calculated empirical splicing efficiencies for individual SJs and normalized them at the gene level (**Fig. 5b**). Regardless of read-depth inclusion criteria, the global splicing efficiency of ncRNAs is significantly lower than that of mRNAs (**Fig. 5c**), consistent with a previous study suggesting ncRNA enriched with alternative splicing for an enormous repertoire of potential regulatory RNAs.^53^ Because calculations of splicing efficiency limited by read depth, spliceAI was utilized to estimate sequence-base splicing probability. While splicing efficiencies at donor sites, acceptor sites and full SJ correlated with the spliceAI scores, the gene-level normalized spliceAI scores recapitulated our observation (**Ext_Fig. 7e-f**).

To determine whether core promoter and enhancer sequence architectures influence splicing activity, we mapped gene-based splicing efficiencies across the ex5_clusters (Methods). Intriguingly, eRNAs derived from CGI enhancers exhibited significantly lower splicing efficiencies than those originating from TATA-box enhancers (**Fig. 5d**). This architectural dichotomy remained consistent across the e_ncRNA and p_ncRNA groups (**Fig. 5e**). Splicing efficiencies for individual gene models are listed in **Table S13**.

### Non-poly(A) eRNAs terminate at structural depletion zone

The molecular mechanisms governing eRNA 3’ end processing remain poorly understood. Despite the recognized role of Integrator-mediated cleavage,^54,55^ its precise site-selection rules remain unresolved. Through analysis of our CFC-seq data, we identified 4,399 poly(A) and 1,971 non-poly(A) recursive TESs within e_ncRNAs. Notably, recursive non-poly(A) TESs exhibit distinct sequence features compared to their poly(A) counterparts, lacking the canonical PAS motif and displaying an enriched cytosine content upstream of the cleavage site (**Fig. 5f**). Given that RNA secondary structure can sterically hinder poly(A)-dependent processing machinery,^56^ we profiled the local thermodynamic secondary structure probability flanking these transcript boundaries. Similar to canonical poly(A) sites, non-poly(A) TESs displayed a localized drop in secondary structure scores (**Fig. 5g, Ext_Fig. 7g, Table S14**). This depletion pattern was consistently observed across all the non-poly(A) ncRNAs and subgroups of regulatory elements (**Fig. 5h**), but not in non-poly(A) mRNA. Notably, Integrator-mediated termination of snRNA preferentially occurs downstream of RNA hairpins.^57^

As non-poly(A) termination often targets transcripts for exosome and XRN degradation, the local structural differences at the TESs may modulate exonucleolytic degradation, thereby generating transiently stable non-poly(A) ncRNAs. Indeed, we propose the upstream cytosine enrichment, which was preserved across CGIap, CGInap and Null e_ncRNAs (**Ext_Fig. 7h)**, acts as a molecular brake to temporarily calibrate transcript lifespan (**Fig. 5i**). Intriguingly, CGI derived eRNAs, which form the most stable predicted upstream hairpin structure, exhibited the highest sensitivity to exosome degradation (**Fig. 4o-p**), suggesting the transient stability of these eRNAs is a result of multi-layers regulation.

Furthermore, this processing model is constrained by local genomic topography. We observed that CGIap enhancers frequently position their CGIs downstream of the TSS (**Ext_Fig. 2f**), directing short eRNAs synthesis towards adjacent gene promoters (**Fig. 4g**). At these loci, over 63% of the non-poly(A) TESs terminated immediately before or within the boundaries of the downstream promoters (**Ext_Fig. 7i**), suggesting a spatial mechanism evolved to prevent downstream transcription interference.^58^ Conversely, this spatial constraint was absent at poly(A) TESs, which exhibited read-through kinetics beyond the promoter boundary in 68% of cases.

Besides providing high CG content to structurally safeguard eRNA from degradation, CGI enhancers recruit MYC binding. Accordingly, we observed a high alignment of CGIs and MYC binding directly at non-poly(A) TESs (**Ext_Fig. 7j-k**). This pattern parallels premature PAS termination events near CGIs that are co-regulated by MYC and nuclear exosome,^59^ and aligns with evidence that MYC/MAX complex binds near TESs of non-coding loci.^60^ Crucially, transcript models linked to MYC-associated TESs displayed significantly higher exosome-sensitivity (**Ext_Fig. 7l**). Taken together, these findings suggest that non-poly(A) TES selection, subsequent PolII termination and exosome degradation may be modulated by a coordinated CGI-MYC axis (**Fig. 5i**).

### Novel transcripts capture unannotated genetic variants

Many disease-associated GWAS and expression quantitative trait loci (eQTL) SNPs map to non-coding intervals, where they potentially disrupt hidden CREs or ncRNAs. Intersecting these genetic variants with our catalog revealed that our novel transcript models increase the exonic coverage of eQTL and GWAS SNPs by 21% and 31% respectively, compared to the GENCODE v39 annotations (**Fig. 6a**). Majority of these newly captured variants reside within the exons of novel loci (**Fig. 6b**). To explore the putative functional effect of these variants, splicing potential from the donor and acceptor sites were tested against these SNPs. Our novel transcript models revealed a 5.8-19% increase in predicted splicing interference mediated by these SNPs (**Fig. 6a-b, Table S15**). Overall, these findings demonstrate that our novel transcript models unmasked functional, disease-relevant variants that escape standard genomic annotation.

**Figure 6.**
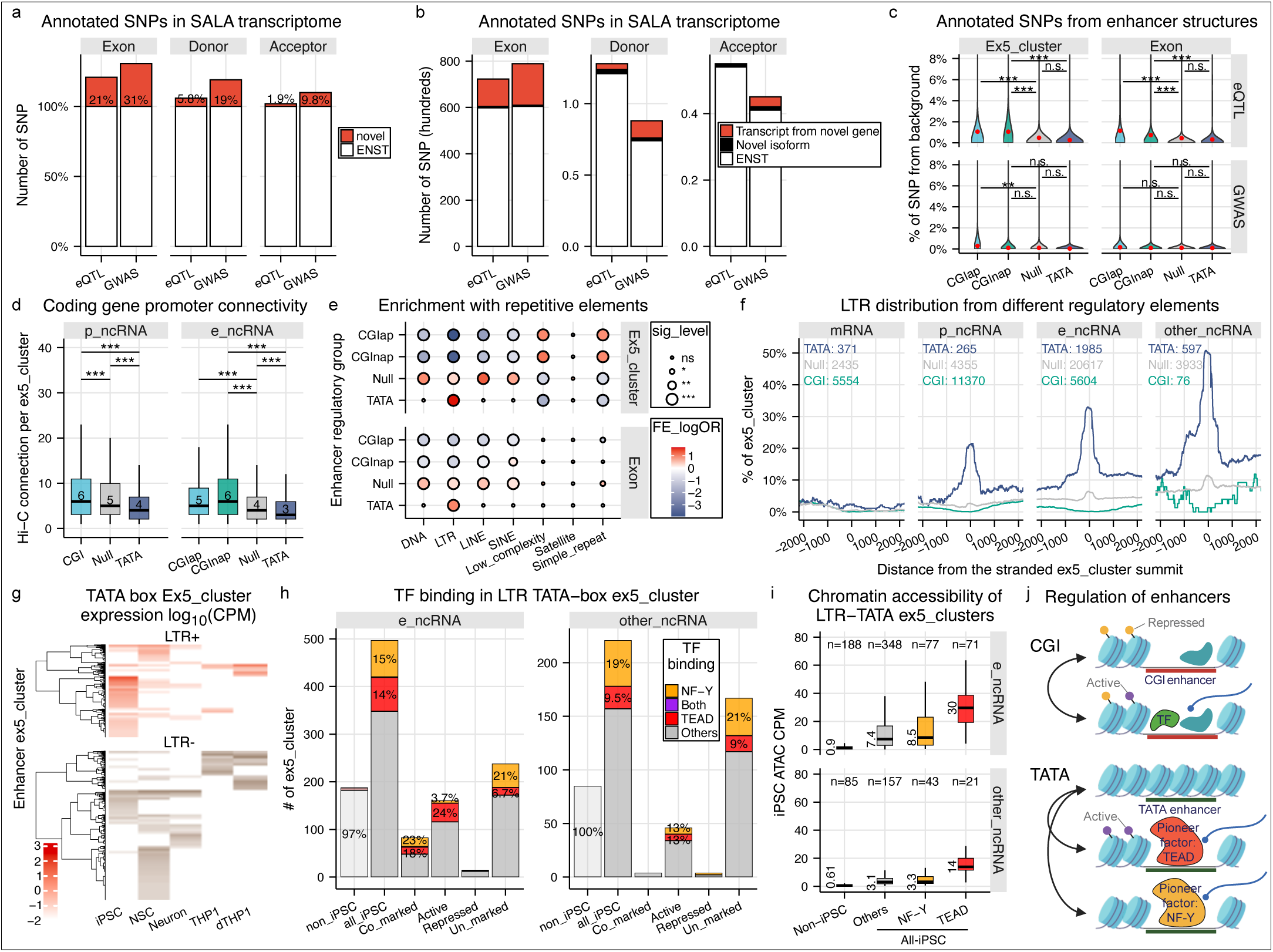
Genetic architecture, retrotransposon exaptation, and pioneering axes of the non-coding genome. **a**, SNPs from eQTL Catalogue and CAUSALdb were intersected with exons of all GENCODE (v39) transcript models and our novel models. SNPs that affect splicing potential from donor sites and acceptor sites were predicted by spliceAI (|delta| ≥ 0.5). A posterior inclusion probability (PIP) ≥ 0.3 was used for eQTLs. The analysis is based on the number of non-redundant SNPs. Percentage increases due to novel annotations are indicated. **b**, SNP intersections as in **a**, grouped by RNA class. **c**, eQTL and GWAS SNPs intersecting e_ncRNAs were grouped by mutually exclusive regulatory elements associated with ex5_clusters. SNP density was normalized to 1 kb of ex5_cluster or transcript sequence. Only SNPs from the “common_all_20180418” panel (NIH) were considered. **d**, Hi-C interactions (5 kb resolution, FDR < 0.01, ≥5 reads) between ex5_clusters linked to ncRNAs and mRNAs. Only clusters with at least one significant interaction are shown. Bins containing multiple regulatory elements (CGI and TATA) were excluded. An interaction observed in any of three cell types was considered positive. **e**, Enrichment of repetitive elements in e_ ncRNA ex5_clusters was tested using Fisher’s exact test. A minimum of 6 nt overlap was required for ex5_cluster regions, and 200 nt for merged exon regions. If multiple repetitive elements overlapped, only the longest overlap was retained. **f**, LTR retrotransposon coverage across ex5_cluster summits, grouped into mRNA, e_ncRNA, p_ncRNA, and other_ncRNA and sub-grouped by CGI, Null, and TATA. The total number of ex5_clusters in each group is shown. **g**, Expression profiles of TATA box-associated enhancer ex5_ clusters across five cell types. Heatmap is divided by presence or absence of LTR elements. **h**, iPSC-derived binding of NF-Y and TEAD4 on non-redundant LTR-TATA ex5_clusters 501 nt regions. Non-iPSC represents LTR-TATA ex5_clusters that are transcribed only in the other 4 cell types. All-iPSC represents LTR-TATA ex5_clusters that are transcribed in iPSC, which were further divided into 4 groups with distinct histone modification: Co-marked (H3K27ac^+^ & H3K27me3^+^), Active (H3K27ac^+^), Repressed (H3K27me3^+^), Un-marked (H3K27ac^-^ & H3K27me3^-^). **i**, Chromatin accessibility of iPSC re-counting the single cell-ATAC-seq fragments into the non-redundant LTR-TATA ex5_clusters 501 nt regions. **j**, Schematic illustrating regulatory mechanisms of CGI-derived versus TATA box-derived enhancers. “Active” and “Repressed” refer to the presence of histone modifications H3K27ac and H3K27me3, respectively. Unless specified, statistical tests were two-sided Wilcoxon; p < 0.05 (*), p < 0.01 (**), p < 0.001 (***), n.s. = not significant.

### CGI enhancers show greater chromatin connectivity

To determine if the structural divergence of enhancers and eRNAs reflects dintinct regulatory potencies, we examined the distribution of genetic variants across enhancer subtypes (**Ext_Fig. 8a**). CGI-associated enhancers exhibited a two-fold higher enrichment of eQTL SNPs compared to TATA-box enhancers (**Fig. 6c**). A similar trend was observed within transcript exons, implying functional potential of the eRNAs.

To further explore the functional impact of different enhancer structures, we analyzed enhancer-promoter interactions using Hi-C data from the Neuron series. Across all three cell types, protein-coding gene promoters exhibited significantly greater interactions with CGI-associated enhancers (**Fig. 6d**), with a consistent trend when analysing each cell type separately (**Ext_Fig. 8b**). These findings align with eQTL SNP enrichment and suggest that CGI enhancers exert a broader regulatory influence than TATA-box enhancers. Subsequent analysis of chromatin accessibility and GC content at Hi-C contact regions revealed that CGI enhancers were associated with increased chromatin accessibility and higher GC content, both of which correlated with contact frequency across cell types (**Ext_Fig. 8c-d**). Notably, CGInap enhancers, which are enriched within SEs (**Fig. 4g**), displayed the highest degree of connectivity. This was further supported by an enrichment test confirming that SEs maintain significantly higher chromatin connectivity than typical enhancers (**Ext_Fig. 8e**), consistent with the elevated Hi-C connectivity reported for orphan CGI enhancers.^61^ Additionally, CGI enhancers preferentially interacted with CGI-containing promoters (**Ext_Fig. 8f**), reinforcing previous findings.^37^

Finally, we incorporate Activity-By-Contact (ABC) model to predict functional targets of these enhancers (**Table S16**). While the gene connectivity predicted by the ABC model (**Ext_Fig. 8g**) closely mirrored the empirical Hi-C connectivity patterns (**Ext_Fig. 8b**), the overall computed enhancer activity of CGI enhancers was significantly greater than that of TATA enhancers (**Ext_Fig. 8h**).

### Exaptation of LTR retrotransposons shapes TATA-box derived landscapes

Given that transposable elements heavily populate the human genome and enhancer-like tCREs were found enriched with SINEs, LINEs, and LTRs (**Ext_Fig. 3l**), we hypothesized that TATA-box derived eRNAs preferentially originate from specific transposable element lineages. Intersecting repeat elements with our ex5_clusters and transcript models revealed a specific enrichment of LTRs with TATA-box enhancers, while CGI enhancers were largely depleted of LTR sequences (**Fig. 6e**, **Ext_Fig.9a**, **Table S17**). Within the LTR lineage, ERV1 family emerged as the predominant donor of TATA-box regulatory architecture (**Ext_Fig. 9b**). This TATA-LTR coupling was also observed from the p_ncRNA and other_ncRNA groups (**Fig. 6g**, **Ext_Fig. 9c**). Remarkably, 57% of TATA-box ex5_clusters of the other_ncRNA associated with LTR, suggesting that LTR exaptation drives the transcription of these unclassified elements. TATA-box eRNAs showed a high coverage of LTR in the transcript body with a median of 56.4% and 28.8% of them spanning more than one exon (**Ext_Fig. 9d**). While TATA-box enhancers displayed high cell-type specificity across differentiation (**Ext_Fig. 9e**), their LTR-derived subgroups were particularly enriched in iPSCs and NSCs (**Fig. 6g**), highlighting their potential role during early developmental stages.

### NF-Y activate transcription from unprimed LTR-TATA enhancers

In contrast to CGI enhancers, TATA-box enhancers were noticeably depleted from Polycomb-repressive chromatin signatures, tracking instead either active or unprimed loci devoid of H3K27ac/me3 (**Fig. 2e**). Concurrently, these elements resided within regions of lower chromatin accessibility (**Ext_Fig. 8c**), suggesting a highly localized, motif-dependent activation mechanism. Motif enrichment analysis within TATA-box ex5_clusters of e_ncRNA and other_ncRNA identified a significant overrepresentation of the NF-Y binding motif, which became more pronounced within LTR-containing fractions (**Ext_Fig. 9f**). Given that NF-Y is a well-established pioneer factor capable of nucleosome remodelling,^62^ we hypothesized that it plays a specialized role in organizing these unprimed loci.

To explore this hypothesis, we integrated independent NF-Y and TEAD ChIP-seq datasets from iPSC (WTC11). Although NF-Y also binds to CGI elements (**Ext_Fig. 9g**), enrichment between NF-Y binding and LTR was only observed from TATA and Null enhancers (**Ext_Fig. 9h**), consistent with prior reports linking NF-Y to LTR architectures.^63,64^ Mapping iPSC-derived NF-Y occupancy across the full differentiation series revealed its binding was almost absent from non-iPSC LTR-TATA enhancers (**Fig. 6h**) that exhibit low accessibility in iPSC (**Fig. 6i**). Instead, NF-Y occupancy was selectively restricted to LTR-TATA enhancer actively transcribed in iPSC, further supporting its active pioneering role.

Remarkably, NF-Y binding was heavily enriched at loci lacking both H3K27ac/me3 modification (**Fig. 6h**, **Ext_Fig. 9i**). This correlation suggests that NF-Y binding may license transcription independently of the classical, flanking histone acetylation cascades traditionally required for locus activation. Because this unprimed activation pattern was absent from Null enhancers (**Ext_Fig. 9i**), and given that TATA-boxes possess high intrinsic affinity for the preinitiation complex,^65^ we propose a model wherein the LTR-TATA combination initiates transcription immediately following NF-Y-mediated nucleosome displacement, bypassing the requirement for widespread histone acetylation (**Fig. 6j**). Operating in parallel, we detected a subtle but distinct enrichment of TEAD4 binding at LTR-TATA enhancers (**Ext_Fig. 9h**). Interestingly, the genomic footprints of NF-Y and TEAD4 rarely overlapped; instead, TEAD4 was specifically enriched at active, H3K27ac-marked LTR-TATA enhancers (**Fig. 6h**, **Ext_Fig. 9i**). This spatial separation points to a complementary, dual-gear pioneering axis: where TEAD4 potentially coordinates with the YAP/TAZ co-activators to drive standard chromatin remodeling and H3K27ac deposition,^66,67^ NF-Y executes rapid, histone-independent transcriptional licensing (**Fig. 6j**). We finally integrate annotated functions of our eRNAs from an external database (**Table S18**).

## Discussion

The integration of CFC-seq with the SALA assembler provides a high-resolution platform that simultaneously resolves TSSs and transcript models from single RNA molecules. By capturing non-poly(A) RNAs through in vitro poly(A)-tailing, this platform unmasks a vast layer of the non-coding transcriptome that was previously invisible to standard long-read sequencing frameworks. By leveraging this empirical precision, we have decoded how core regulatory sequence architectures dictate the life cycle and biophysical fate of eRNAs.

Our findings expose a fundamental structural dichotomy governing enhancer-derived transcription, which we propose a predictive genomic code. CGI enhancers generate short, rarely spliced, and highly exosome-sensitive transcripts. At the chromatin level, these elements display elevated spatial connectivity and are noticeably enriched with repressive H3K27me3 marks. This multi-layered restriction of RNA accumulation likely promotes localized *cis*-regulatory functions, potentially facilitating phase-separated condensate formation at active enhancer-promoter interacting loci.^68^ In contrast, TATA-box-associated enhancers produced long, stable and multi-exonic eRNAs well-suited to act in *trans* across the nuclear space.^69^ We demonstrate that this structural complexity was systematically acquired through the evolutionary co-option of LTR retrotransposons, which have been seamlessly hardwired into the host’s core developmental signaling networks.^70,71^

Crucially, transcription at these elements is driven by a distinct dual-gear pioneering axis. The capacity of NF-Y to license robust transcription at H3K27ac/me3 deficient loci independently of classical flanking histone acetylation cascades challenges conventional models of transcription activation. We propose that the TATA-box provides an ultra-rapid, streamlined shortcut for initiation. Operating in parallel with the classical, histone-depositing pioneering activity of TEAD4, this dual-gear mechanism calibrates transcriptional plasticity during early differentiation.

Finally, our platform reveals the structural determinants that dictate transcript termination. At non-poly(A) eRNA TESs, we observed a constrained, ∼25-nucleotide zone of local secondary structure depletion. Rather than representing random degradation stalling, we propose this depletion zone serves as a primary cleavage site for polymerase II termination and subsequent entry of the RNA exosome. Within this framework, the upstream cytosine-rich terminal features act as a molecular brake, temporarily halting exonucleolytic degradation to precisely calibrate eRNA turnover and establish transient stability.

## Limitations

Several methodological constraints warrant considerations. First, the integration of size-selection protocols to optimize read-length distribution (**Fig. S4**) may introduce representation biases across different RNA classes. Second, while length-dependent PCR amplification optimizes CFC-seq for transcript structural characterization and isoform resolution, it can skew absolute quantification across transcripts with different lengths. Finally, while CGI and TATA-box frameworks define highly active, mechanistically distinct functional subsets, the majority of transcribed enhancers lack these specific core motifs. Interestingly, approximately 40% of these enhancers contain transposable elements within their RNA bodies. Deciphering the precise activation and processing dynamics of these unclassified elements remains a vital frontier.

## Data Availability

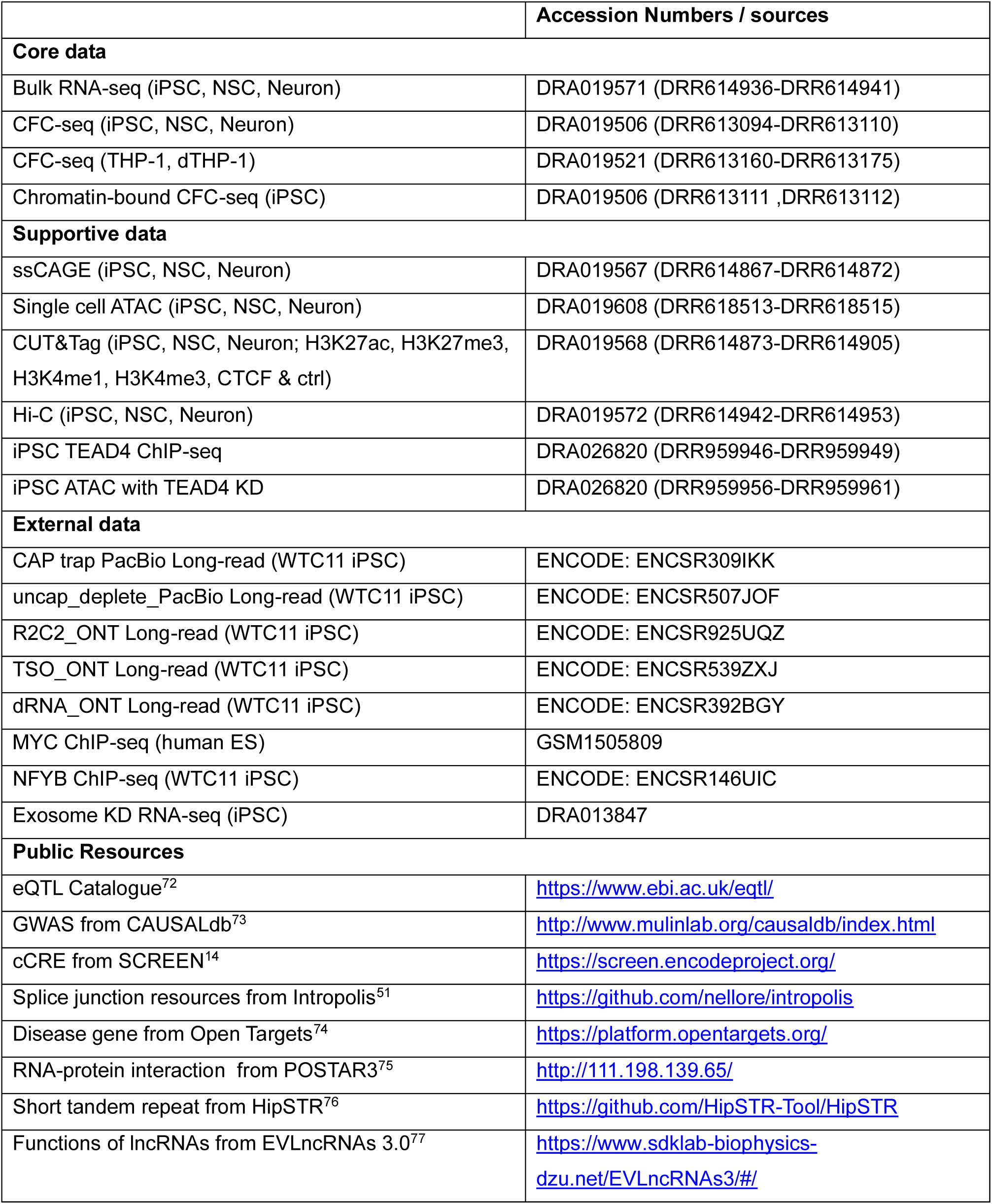

### Code Availability

Code and limited data is available in https://github.com/fantom-prj/CFC_eRNA_analysis SALA software package is available in https://github.com/fantom-prj/SALA

## Supplementary Materials

Online Methods, extended data figures and supplementary figures are available to download from supplementary materials.

## Supplementary information

Supplementary tables are available in https://figshare.com/articles/dataset/Supplementary_Table/27332796/2

**Table S1:** Poly(A) prediction for the 3’ end of all the reads

**Table S2:** Annotation of tCREs

**Table S3:** Annotation of ex5_clusters

**Table S4:** Transcript information from the SALA finalized transcriptome (for downstream analysis)

**Table S5:** Transcript information from the SALA raw transcriptome

**Table S6**: Transcript information from the SALA robust transcriptome

**Table S7**: Transcript ID and gene ID links to external databases

**Table S8:** Bambu quantification of the SALA finalized transcriptome

**Table S9:** Isoform switcher results

**Table S10:** RNA features from ex5_clusters

**Table S11:** RNA features from transcript models

**Table S12:** Transcript information from the finalized transcriptome of iPSC chromatin-bound RNA

**Table S13:** Splicing efficiency of transcript and gene models

**Table S14:** Non-poly(A) TES with structural depletion

**Table S15:** Results from spliceAI showing all donor sites and acceptor sites affected by SNPs

**Table S16:** Enhancer-based result of ABC-model

**Table S17:** Repeat elements intersect with ex5_cluster and transcript models

**Table S18**: eRNA-based characterization with function from EVLncRNAs3

**Table S19**: Molecular details and oligo sequences of CFC-seq

**Table S20:** External TSS regions

## Contributions

Investigation: C.P., H.T., C.L., I.L., C.W.Y., C.H.

Conceptualization: C.W.Y., C.H., P.C.

Manuscript writing: C.P., H.T., C.W.Y.

Computational analysis: J.M., R.U., J.C., M.V., X.S., M.T., N.H., M.F., V.R., A.D.S., Y.C., C.U.,

C.A.G., R.A., F.D., L.P., W.K., R.P., G.F., R.S.S., A.L., C.L., I.L., C.P., C.W.Y., C.H.

Experimental analysis: K.Y., Y.S., T.K., M.M., H.N., M.K., T.K., W.H.Y., C.P., H.T.

Project administration: T.K., M.K., P.C., H.T., C.W.Y.

Project supervision: J.W.S., K.O., B.L., B.B., J.G., F.N., V.H., L.C., S.G., M.B., M.K., P.C., H.T., C.L., I.L., C.W.Y., C.H.

Funding acquisition: J.W.S., P.C.

Database management: T.N., T.K.

## Acknowledgements

As part of the FANTOM6 project, this study is supported by many other studies of the project. We thank all the members in the FANTOM consortium for excellent support, discussion and inspiration. Sequencing was performed by the Laboratory for Genotyping Development in RIKEN IMS, the National facilities (formerly the sequencing facility) and the Centre of Genomics in Human Technopole. We acknowledge the Access and services provided by the National Facility for GENOMICS, Fondazione Human Technopole, Milan, Italy. We thank Nobuyuki Takeda and Teruaki Kitakura (RIKEN) for their support of the IT infrastructure for the FANTOM6 collaboration, and Emi Ito (RIKEN) for her administrative support.

---

**Extended Data Figure 1 | Technical validation and read statistics of the CFC-seq platform.**

**a**, CFC-seq experimental workflow. **b**, Sequencing statistics and library metrics for all generated datasets. **c**, Quality control breakdown of long reads (base-calling score ≥ 10) into incomplete (missing adaptors), non-mappable, and mappable fractions. **d**, Intersection of mRNA 5′ ends with SCREEN cCREs, ATAC-seq peaks, and FANTOM5 CAGE clusters. Long-read datasets from iPSC WTC11 were obtained from ENCODE LRGASP.^11^ **e**, Enrichment of non-polyadenylated (non-poly(A)) transcripts in libraries following *in vitro* poly(A)-tailing (PAT). **f**, Evaluation of internal priming based on genomic sequence composition downstream of read 3′ ends. **g**, Polyadenylation signal (PAS) motif frequency (motif score >3, −35 to −5 bp upstream of the 3′ end) in reads across mRNA, promoter-ncRNA, and eRNA classes, comparing ±PAT libraries. **h**, Genomic distribution of non-PAS-associated mRNA 3′ ends to explore fraction of potential nascent RNA. **i**, Characterization of emergent mRNA 3′ ends lacking PAS motifs in THP-1 ±PAT libraries. Left: genomic localization relative to coding exons and introns. Right: nucleotide composition frequency surrounding 3′ end sites (±30 nt). **j**, Frequency of PAT-specific mRNA 3′ ends across genomic features, showing accumulation at splice donor sites. **k**, Overlap of genes detected by CFC-seq ±PAT and the distribution of their RNA classes. **l**, ROC curve for the sequence-based poly(A) classifier, validated against FLAM-seq 3′ ends (as positives) and PAT-specific 3′ ends (as negatives).

**Extended Data Figure 2 | Identification of genuine TSSs, clusters, and tCREs from 5′ ends of CFC-seq reads.**

**a**, Summary of reads processed with SCAFE for identification of genuine TSS clusters and tCREs. **b**, Neuron-series tCREs overlap with ATAC-seq-defined open chromatin from the same samples. **c**, Unclassed tCREs were stratified by ACTA-seq support, GENOCODE annotation status, and the originating cell type. **d**, Overlap of tCREs with chromatin states defined by ChromHMM using CUT&Tag data (H3K27ac, H3K27me3, H3K4me1, H3K4me3, CTCF) from three Neuron-series cell types. Colors indicate summit positions relative to GENCODE models. **e**, Genomic features of major-strand tCREs (n = 57,597) categorized as promoter-like, enhancer-like, or unclassed. Analyses include PhastCons conservation (4way–100way), overlap with GC content and CGI (UCSC), and motif enrichment (TATA box: MA0108.3, INR: POL002.1, from JASPAR). Summits were extended ±2 kb for conservation and GC content, and ±50 bp for motif analysis; backgrounds were exon- and tCRE-masked. **f**, Distribution of CGI and TATA box around summits of major-strand and minor-strand tCREs of all the enhancers and enhancers divided into adjacent to promoter (enhancer-AP) and not adjacent (enhancer-NAP). **g**, Proportion of enhancer-like major-strand tCREs with antisense minor-strand tCREs and their regulatory features. **h**, Classification and naming schema for regulatory elements. **i**, Distance of different classes of enhancer with their closest promoter cCRE. **j**, Enrichment of chromatin states with CGI, CGIap, CGInap, and TATA elements in enhancer-like tCREs across the 3 cell types. Enrichment test is performed by using Fisher’s exact; p < 0.05 (*), p < 0.01 (**), p < 0.001 (***), n.s. = not significant.

**Extended Data Figure 3 | Features of tCREs related to directionality, super enhancers and repetitive elements.**

**a**, Schematic illustrating the classification of tCREs as bidirectional (2D) or unidirectional (1D) based on transcription activity from both strands. **b**, Read count distribution of stitched H3K27ac peaks generated by ROSE. Regions exceeding the dotted line threshold were classified as super enhancers. **c**, Size distribution of super enhancers identified in the three analyzed cell types. **d**, Sequence conservation around tCRE summits for 2D and 1D promoters and enhancers, based on phastCons scores as described in **Extended Data Figure 2e**. **e**, Proportion of 2D and 1D tCREs (major strand only) containing CpG islands (CGI), TATA boxes, or both (Mix). **f**, Sequence conservation (phastCons 17-way scores) around the summit (±500 nt) of 2D and 1D, promoter-like and enhancer-like tCREs, stratified by presence of CGI or TATA box. Only major strand tCREs were included. **g**, Classification of mapped reads analyzed by SCAFE from CFC-seq and CAGE datasets into: not qualified (5′ soft-clipping >3 nt), qualified without unencoded G, and qualified with unencoded G. **h**, Distribution of single-nucleotide TSSs found uniquely in CFC-seq, in both CFC-seq and CAGE, or uniquely in CAGE. **i**, Number of tCREs detected by CFC-seq, CAGE, or both at varying read count thresholds. tCREs were identified by joint SCAFE analysis of Neuron series CFC-seq and CAGE datasets. **j**, Expression correlation per cell type between CFC-seq and CAGE, shown as RLE-normalized CPM values for each tCRE (each dot represents one tCRE). tCREs were identified by joint SCAFE analysis. **k**, TSS cluster identification coverage by CFC-seq, CAGE paired-end (PE), and CAGE single-end (SE) across increasing MAPQ thresholds. Coverage is shown relative to union TSS clusters at MAPQ = 0. **l**, Repetitive element divergence (as a proxy for evolutionary age) for repeats overlapping CFC-specific TSS clusters at MAPQ 20 versus others. **m**, Proportion of major strand tCREs containing repetitive elements, colored by repeat class. “Multiple” indicates overlap with more than one repeat class. Unless specified, statistical tests were two-sided Wilcoxon; p < 0.05 (*), p < 0.01 (**), p < 0.001 (***), n.s. = not significant.

**Extended Data Figure 4 | Transcript and gene models identified by SALA.**

**a**, Number of transcript models in the finalized transcriptome with 5′ end support from either SCAFE or GENCODE annotations. **b**, Number of transcript models in the finalized transcriptome with 3′ end support. Poly(A)-positive (P(A)) transcripts were defined based on a positive poly(A) classifier score or the presence of a polyadenylation signal (PAS) with motif score >3. **c**, Summary of filtering criteria applied to generate different transcriptome versions for benchmarking. **d**, Table comparing the number of transcript and gene models identified by SALA and other transcriptome assembly tools under different filtering conditions. **e**, Venn diagram showing the overlap of detected GENCODE transcript (ENST) across different software tools. Right panel shows the category of transcript model associated to the ENST observed from other tools but not in SALA. Only Full Splice Match (FSM) and Incomplete Splice Match (ISM) were shown. The sum of the distance difference from the two ends between the FSM transcript models and associated ENST was shown in the distribution plot. **f**, Transcript categories assigned to novel transcript models using SQANTI3 across different software outputs. FSM: Full Splice Match, ISM: Incomplete Splice Match, NIC: Novel in Catalog, NNC: Novel Not in Catalog. **g**, Number of transcript models classified into three categories: “GENCODE transcript,” “Novel isoform from ENSG,” and “Transcript from novel transcriptional unit (TU).” **h**, Proportion of “Transcript from novel TU” models whose 5′ ends overlap with open chromatin (ATAC-seq), ENCODE SCREEN candidate *cis*-regulatory elements (cCREs), or SCAFE genuine TSS clusters. **i**, Gene and transcript class assignments in the SALA Final transcriptome. **j**, Number of ncRNA transcript models included in three groups (GENCODE genes, lncRNA from novel TU, and short_ncRNA from novel TU) according to criteria used to define non-coding RNA classes in this study.

**Extended Data Figure 5 | Transcript and gene models identified by SALA.**

**a**, UpSet plot showing novel transcript models identified in this study and their overlap with FANTOM CAT, LncBook, RefSeq, and GENCODE v47. Transcript model identity was determined by SALA; schematic shown on the left. **b**, UpSet plot of novel gene models and their overlap with the same four databases, as identified by SALA. **c**, Counts of transcript models annotated as ncRNA or other classes across the four databases. **d**, Cell-type specificity (Gini index) of gene models annotated in any of the public databases or completely novel, across the five studied cell types. Right, maximum expression level across the five cell types for each group. **e**, Proportion of gene and transcript models in the final CFC-seq transcriptome also detected by short-read RNA-seq in iPSC, NSC, and Neuron. **f**, Classification of novel gene and transcript models (detected or undetected by short-read RNA-seq) into ncRNA subclasses. **g**, Correlation of expression levels from long- and short-read data using the finalized transcriptome, limited to detectable GENCODE models. **h**, Expression levels of transcript models from short- and long-read libraries, grouped by transcript length. **i**, Classification of novel isoforms of protein-coding genes. "Full-splice match" indicates isoforms with all splice junctions matching a GENCODE transcript; "incomplete-splice match" includes partial matches. Alternative TSS and TES was assessed by ex5_clusters and ex3_clusters relative to best-matched GENCODE transcripts; skipped for unmatched cases (novel in catalog or novel not in catalog). Right, predicted coding potential by CPAT. **j**, For novel isoforms with alternative TES, presence of poly(A) signals in the isoform and the matched GENCODE transcript was assessed via PAS and poly(A) predictions. Isoforms were further stratified by whether their 3′ end overlaps a splice donor site. **k**, Venn diagram showing genes with both significantly up-and down-regulated isoforms across three comparisons. **l**, Venn diagram of genes with isoform switching involving alternative TSS, TES, and exon usage.

**Extended Data Figure 6 | Factors influencing RNA transcription.**

**a**, Transcript lengths of RNA groups defined by ex5_clusters. **b**, Transcript length of eRNAs defined using various enhancer annotation methods. **c**, Proportion of RNA groups identified in this study among enhancers defined by different methods. **d**, Median exon number of eRNAs by enhancer definition method; percentages indicate the proportion with a median exon count > 1. **e**, Venn diagrams showing overlap of ex5_clusters across TATA, CGIap, and CGInap, and the subset with downstream CGI (dCGI) for e_ncRNA. The p_ncRNA was divided into TATA and CGI with dCGI subset. **f**, Transcript length of p_ncRNAs compared across promoter features (as defined in Figure 4c). **g**, Exon count of p_ncRNAs across promoter features; percentages indicate groups with more than one exon. **h**, Transcript length of four mutually exclusive p_ncRNA groups, based on individual promoter features. **i**, Median exon number for the four mutually exclusive p_ncRNA groups; percentages indicate groups with more than one exon. **j**, Length variation within ex5_clusters represented by relative MAD (MAD/median); plotted alongside poly(A) RNA rate (defined in Figure 4k). **k**, SALA analysis of chromatin-bound CFC-seq from iPSCs using the finalized transcriptome plus GENCODE v39. Shown are ncRNA genes originating from GENCODE, the novel ncRNA identified from the finalized transcriptome (ONTG), or the chromatin-bound library alone. **l**, Transcript length differences for shared transcript models between the chromatin-bound and main datasets, grouped by poly(A) status in the main dataset. **m**, Exosome sensitivity of transcript models (1,935 e_ncRNAs and 3,841 p_ncRNAs) with CPM > 0.25, based on RNA-seq of EXOSC3 knockdown in iPSC. Sensitivity (log₂ fold change) was estimated using edgeR on chromatin-bound vs. total RNA CFC-seq (quantified by bambu with the SALA finalized transcriptome updated with chromatin-bound models). **n**, Enrichment of individual RBP interactions from exons of coding vs. non-coding, and enhancer- vs. promoter-derived transcripts. **o**, Enrichment of exosome sensitivity in p_ncRNAs across different promoter features (defined in Figure 4n). Unless specified, statistical tests were two-sided Wilcoxon; p < 0.05 (*), p < 0.01 (**), p < 0.001 (***), n.s. = not significant.

**Extended Data Figure 7 | Splicing efficiency and eRNA transcription termination.**

**a**, Number and percentage of splice junction (SJ) classes of all the SJs from external reference databases. “Short read” refers to confident SJs derived from short-read RNA-seq in the Neuron series. **b**, Novel SJs absent from the databases in (a) were classified as canonical or non-canonical. Plots show the relationship between the mean maximum base-calling score across the 6 nt surrounding each SJ and the SJ’s occurrence. SJs were considered technically confident if all 6 nt scored ≥10 and occurred ≥3 times. **c**, SJs from the final transcriptome were assigned to the four RNA classes, allowing for overlaps. For each class, the percentage of transcripts with and without fully supported SJs is shown. Positive SJ support was defined as either present in the databases in (**a**) or technically confident per (**b**). **d**, Total number of transcript models containing ≥1 SJ and percentage of each SJ class based on motif composition. **e**, Gene-level splicing efficiency compared to SpliceAI score across different occurrence thresholds. **f**, SpliceAI scores were summarized at the gene level and normalized by intron count. Gene models without SJs were excluded. SJs were filtered using three occurrence thresholds (sum of span and splice counts at both donor and acceptor sites). **g**, Evidence of RNA structural depletion (fold change >15%) at non-polyadenylated TESs, including both recursive and non-recursive TESs, based on RNAfold-derived hairpin scores. **h**, Nucleotide composition surrounding recursive transcription end sites (TESs; ≥3 read support) of non-poly(A) eRNAs, stratified by regulatory elements. **i**, Position of CGI enhancer-derived RNA TESs relative to the downstream promoter (SCREEN promoter cCRE). Not Reaching: TES is located > 500nt upstream the promoter region, Terminate: TES is located within 500nt upstream or inside the promoter, Read-through: TES is located downstream of the promoter. **j**, Distribution of CpG islands proximal to TESs of non-polyadenylated RNA types, restricted to recursive TESs. **k**, Distribution of MYC binding sites (from ChIP-seq dataset GSM1505809) near TESs of non-polyadenylated RNA types, restricted to recursive TESs. **l**, Exosome sensitivity of transcript models was weighted and grouped by TES and compared between TES with and without MYC binding, stratified as different classes. Unless specified, statistical tests were two-sided Wilcoxon; p < 0.05 (*), p < 0.01 (**), p < 0.001 (***), n.s. = not significant.

**Extended Data Figure 8 | Genetic variants and enhancer connectivity.**

**a**, Number of SNPs associated with e_ncRNAs grouped by mutually exclusive regulatory elements of ex5_clusters. SNP counts were normalized by the length of the ex5_cluster or transcript model (per kb). The “all_SNP” catalog refers to common SNPs from the NIH dataset. **b**, Hi-C interactions (5 kb resolution, FDR < 0.01, ≥5 reads) were used to infer ncRNA–mRNA connectivity via ex5_clusters. Only e_ncRNA ex5_clusters with ≥1 connection were included. Bins containing ex5_clusters with multiple regulatory elements (CGI and TATA box) were excluded. Median connection counts per ex5_cluster are shown, analyzed separately for each of the three cell types. **c**, Chromatin accessibility from matching ATAC-seq samples was mapped to 5 kb bins containing a single regulatory element (multi-element bins excluded). Median read counts and Spearman correlation between chromatin accessibility and Hi-C connectivity (as in panel **b**) are shown. **d**, GC content was calculated for 5 kb bins as in (**c**), and Spearman correlation with Hi-C connectivity (as in Figure 6d) is shown. **e**, Hi-C connectivity across all three cell types was compared against tCRE directionality and super-enhancer (SE) clustering. **f**, Two-sided Fisher’s exact test was used to assess enrichment across regulatory element combinations. ncRNA source was grouped as in panel **b**; mRNA promoters were classified by presence or absence of CGI or TATA box. Odds ratios are shown for each combination. **g**, Gene connectivity of enhancers estimated by ABC-model. Analysis was performed on merged tCREs and linked to the features of ex5_clusters. Merged tCREs containing ex5_clusters with multiple regulatory elements (CGI and TATA box) were excluded. Only the enhancers with at least one connection with ABC score > 0.02 were included. **h**, Same analysis as (**g**) showing the mean ABC score for each enhancer. Unless specified, statistical tests were two-sided Wilcoxon; p < 0.05 (*), p < 0.01 (**), p < 0.001 (***), n.s. = not significant.

**Extended Data Figure 9 | TATA box enhancers enriched with LTR element are activated by NF-Y.**

**a**, Proportion of ex5_clusters and exons (grouped by cluster) intersecting with repetitive elements, applying the same criteria as in Figure 6e. **b**, Family composition of LTR elements intersecting enhancers and eRNAs. **c**, Enrichment with LTR across different groups in p_ncRNA and other_ncRNA was tested using Fisher’s exact test. **d**, LTR coverage percentage over exons of transcript models that intersected with LTR, grouped by promoter type and regulatory element. Percentages on the top indicate the proportion of transcript models with LTR overlap across ≥2 exons. **e**, Expression profile of e_ncRNA ex5_clusters across five cell types, grouped into CGI, Null, and TATA categories. **f**, Enrichment ratio of transcription factor motifs was shown for those with FDR < 0.05 and enrichment ratio >12 and expression levels >1 CPM. Motif analysis was performed on non-redundant 501 nt CRE sequences (summit +100 nt & −400 nt). **g**, Distribution of NF-Y binding across ex5_cluster summits, grouped into mRNA, e_ncRNA, p_ncRNA, and other_ncRNA and sub-grouped by CGI, Null, and TATA. The total number of ex5_clusters in each group is shown. **h**, Enrichment between LTR and binding of NF-Y and TEAD considering only ex5_cluster transcribed in iPSC. Fisher’s exact derived odds ratio and p-values of the 12 subgroups were shown. **i**, Enrichment between chromatin states and binding of NF-Y and TEAD. Fisher’s exact derived odds ratio and p-values were shown from TATA and Null subgroup of e_ncRNA ex5_clusters transcribed from iPSC, including only LTR^+^. Co-marked (H3K27ac^+^ & H3K27me3^+^), Active (H3K27ac^+^), Repressed (H3K27me3^+^), Un-marked (H3K27ac^-^ & H3K27me3^-^). Unless specified, statistical tests were two-sided Wilcoxon; p < 0.05 (*), p < 0.01 (**), p < 0.001 (***), n.s. = not significant.

**Supplementary Figure 1 | Features of extended 5’ end clusters (ex5_clusters)**

**a**, Comparison between TSS clusters, ex5_clusters, and tCREs, showing average coverage of each feature across the others. **b**, Number of transcript models originating from each 5′ end feature (TSS cluster, ex5_cluster, and tCRE). **c**, Distribution of DNA regulatory elements associated with ex5_clusters (linked to transcript models in the finalized transcriptome, *n* = 75,658) in reference to summit position, categorized as promoter-like, enhancer-like, or unclassed. Both major and minor strand clusters were included. See Methods and **Extended Data Figure 2e** legend for classification criteria. **d**, Presence of CGI, TATA box, or both (mix) in 2D and 1D ex5_clusters, assessed using non-redundant 1001 nt windows. **e**, Conservation scores (phastCons 17-way) of ex5_clusters grouped as 2D or 1D promoters or enhancers, and further stratified by presence or absence of CGI or TATA box. Mean phastCons values were calculated within ±500 nt of cluster summits. **f**, Distance of different classes of enhancer from promoter cCRE. **g**, Coordinations of enhancer-like ex5_clusters (major strand only) were intersected with iPSC H3K27Ac and H3K27Me3 CUT&Tag data and grouped into C-marked (both 27ac and 27me3), Active (27ac alone), Repressed (27me3 alone), and Un-marked (Neither). TSS counts ≥1 were considered positive in expression. Top panel: percentage of CGI-positive (TATA-negative) ex5_cluster per enhancer state; second to bottom panels: CGIap-positive (CGI-positive ex5_clusters that are adjacent to promoter), dCGInap-positive (dCGI-positive ex5_clusters that are not adjacent to promoter) and TATA-positive (CGI-negative). **h**, Enrichment of enhancer states with different classes of ex5_clusters as shown in (**g**). **i**, Same analysis as (**h**), restricted to expressing ex5_clusters per cell-type and incorporating cell-type-specific chromatin states. Unless otherwise stated, statistical comparisons were performed using a two-sided Wilcoxon test. Fisher’s exact test was used in (**h**) and (**i**) to calculate odds ratios (OR). p < 0.05 (*), p < 0.01 (**), p < 0.001 (***), n.s. = not significant.

**Supplementary Figure 2 | Example genes with significant isoform switching due to switched splice junction usage.**

**a**, Genome browser view of the *AP1S2* locus. Tracks include the finalized SALA transcriptome alongside GENCODE v39 gene and transcript annotations. Switched isoform track displays only transcript models with significant changes in expression fraction between any pair of the three cell types (determined by isoformSwitchAnalyzeR). Color scale represents CPM values normalized within each cell type (from bambu). CFC-seq tracks display the summed signal of 5′ ends, 3′ ends, and exons across all samples. Tracks for ATAC-seq, cCRE promoter/enhancer regions, and SCAFE tCREs show genomic annotation at this locus. **b**, Isoform expression fractions of *AP1S2* in the three cell types, as estimated by isoformSwitchAnalyzeR. **c**, Predicted ORF alignment of the switched isoforms of *AP1S2*. **d**, Genome browser view of the *PORCN* locus with track structure as in (**a**). **e**, Isoform expression fractions of *PORCN* in the three cell types. **f**, Predicted ORF alignment of the switched isoforms of *PORCN*.

**Supplementary Figure 3 | Example genes with significant isoform switching due to switched promoter usage.**

**a**, Genome browser view of the *PRKCZ* locus. Tracks include the finalized SALA transcriptome alongside GENCODE v39 gene and transcript annotations. Switched isoform track displays only transcript models with significant changes in expression fraction between any two of the three cell types (as identified by isoformSwitchAnalyzeR). Color scale indicates CPM values derived from bambu, normalized within each cell type. CFC-seq tracks show the aggregated signal for 5′ ends, 3′ ends, and exons across all samples. Additional tracks show annotations from ATAC-seq, cCRE promoter and enhancer regions, and SCAFE-defined tCREs. **b**, Isoform expression fractions of *PRKCZ* in the three cell types, estimated by isoformSwitchAnalyzeR. **c**, Chromatin activity of corresponding ex5_clusters derived from ATAC-seq signal normalized by library size. **d**, Predicted ORF alignment of the switched isoforms of *PRKCZ*. **e**, Genome browser view of the *CA14* locus, track layout as in (a). **f**, Isoform expression fractions of *CA14* in the three cell types. **g**, Chromatin activity of ex5_clusters derived from ATAC-seq signal. **h**, Predicted ORF alignment of the switched isoforms of *CA14*.

**Supplementary Figure 4 | Length distribution revealed by bioanalyser before and after PCR amplification.**

Length distribution before and after PCR and size selection for **a**, iPSC, **b**, NSC and **c**, Neuron.

