## Supplementary Figure 1-4 for "Genomic codes governing enhancer RNA fate"

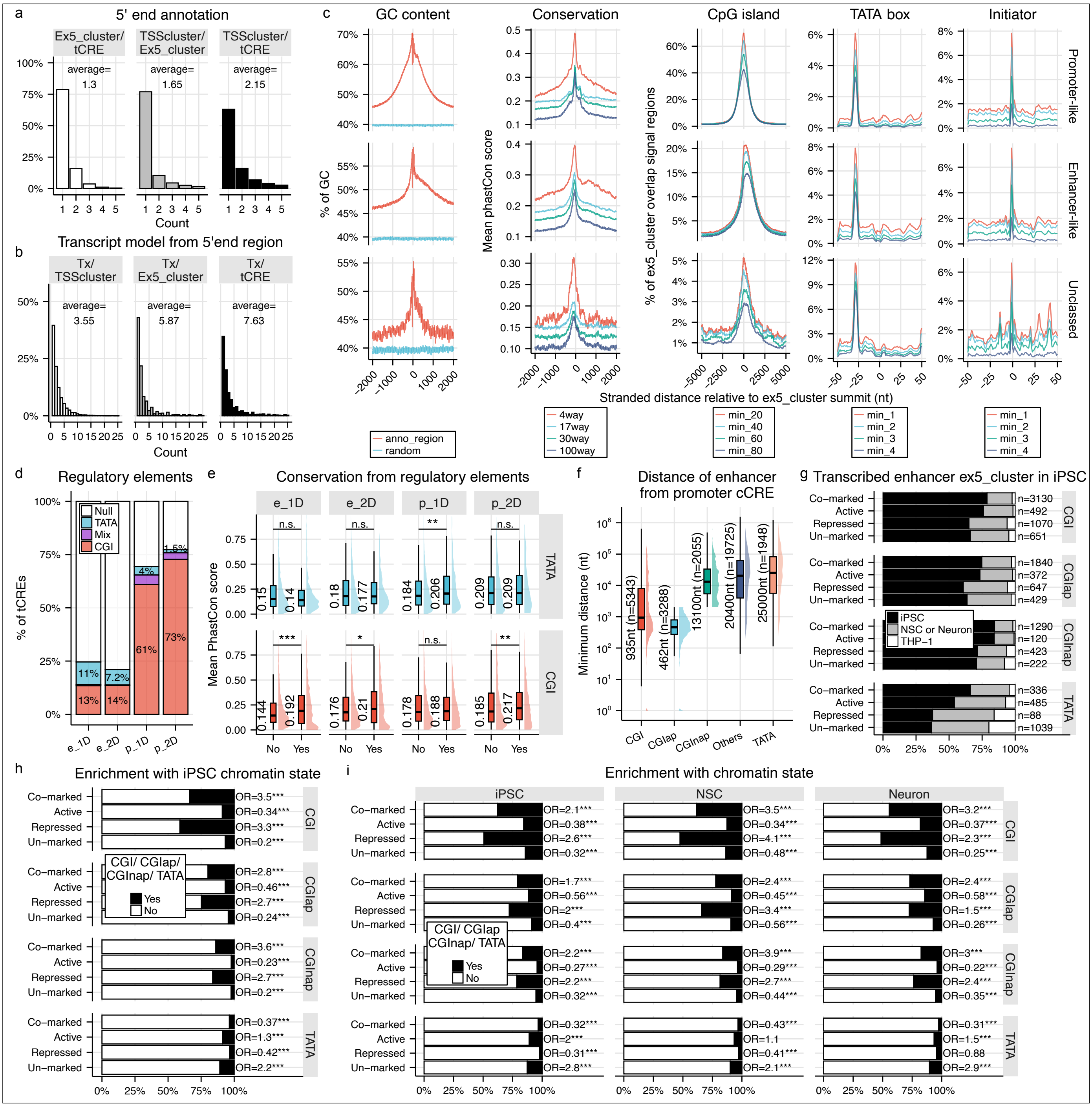


**Supplementary Figure 1 | Features of extended 5' end clusters (ex5_clusters)**

**a**, Comparison between TSS clusters, ex5_clusters, and tCREs, showing average coverage of each feature across the others. **b**, Number of transcript models originating from each 5′ end feature (TSS cluster, ex5_cluster, and tCRE). **c**, Distribution of DNA regulatory elements associated with ex5_clusters (linked to transcript models in the finalized transcriptome, *n* = 75,658) in reference to summit position, categorized as promoter-like, enhancer-like, or unclassed. Both major and minor strand clusters were included. See Methods and **Extended Data Figure 2e** legend for classification criteria. **d**, Presence of CGI, TATA box, or both (mix) in 2D and 1D ex5_clusters, assessed using non-redundant 1001 nt windows. **e**, Conservation scores (phastCons 17-way) of ex5_clusters grouped as 2D or 1D promoters or enhancers, and further stratified by presence or absence of CGI or TATA box. Mean phastCons values were calculated within ±500 nt of cluster summits. **f**, Distance of different classes of enhancer from promoter cCRE. **g**, Coordinations of enhancer-like ex5_clusters (major strand only) were intersected with iPSC H3K27ac and H3K27me3 CUT&Tag data and grouped into Co-marked (both 27ac and 27me3), Active (27ac alone), Repressed (27me3 alone), and Un-marked (Neither). TSS counts ≥1 were considered positive in expression. Top panel: percentage of CGI-positive (TATA-negative) ex5_cluster per enhancer state; second to bottom panels: CGIap-positive (CGI-positive ex5_clusters that are adjacent to promoter), dCGInap-positive (dCGI-positive ex5_clusters that are not adjacent to promoter) and TATA-positive (CGI-negative). **h**, Enrichment of enhancer states with different classes of ex5_clusters as shown in (**g**). **i**, Same analysis as (**h**), restricted to expressing ex5_clusters per cell-type and incorporating cell-type-specific chromatin states. Unless otherwise stated, statistical comparisons were performed using a two-sided Wilcoxon test. Fisher's exact test was used in (**h**) and (**i**) to calculate odds ratios (OR). $p < 0.05$ (*), $p < 0.01$ (**), $p < 0.001$ (***), n.s. = not significant.

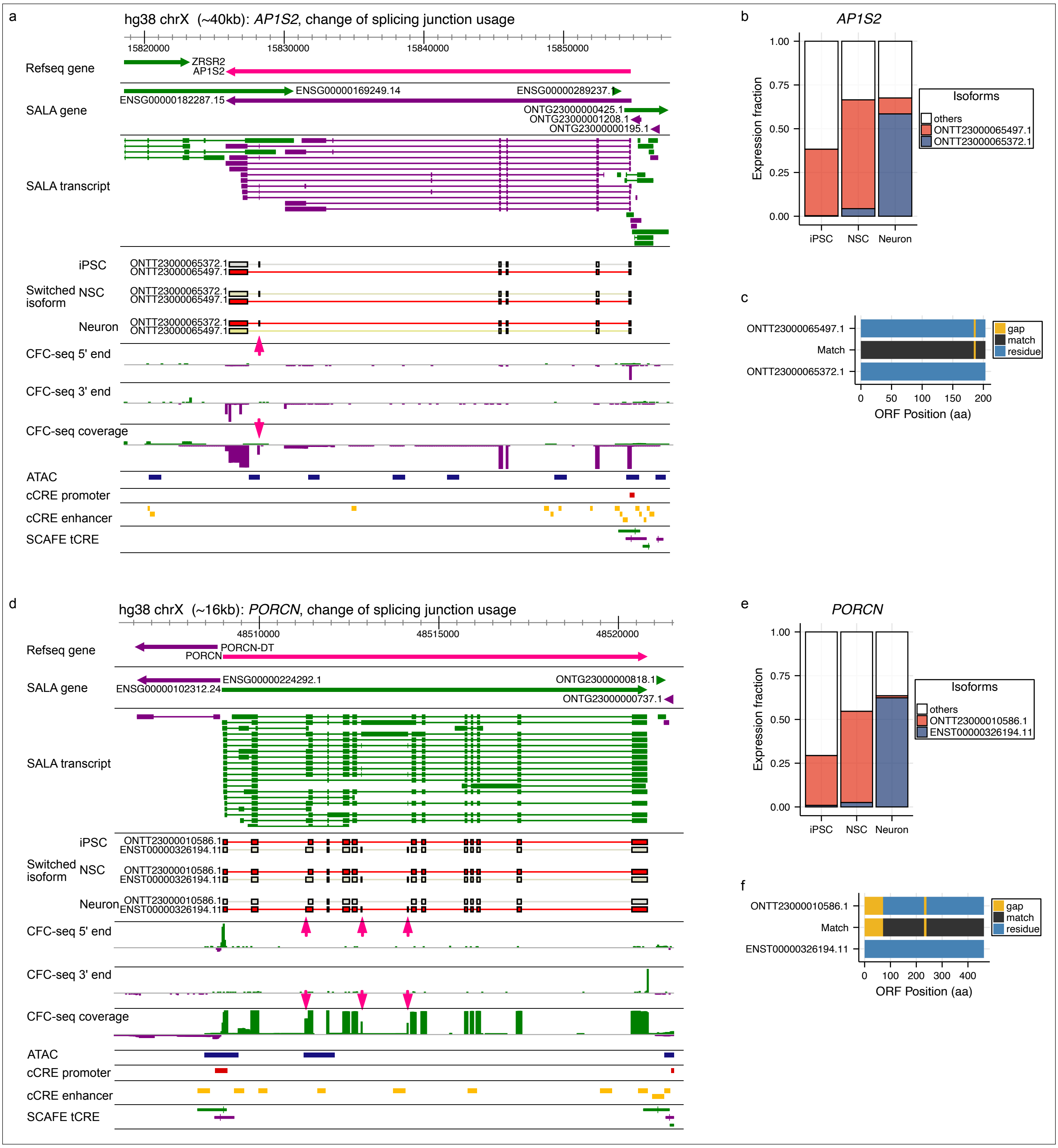


**Supplementary Figure 2 | Example genes with significant isoform switching due to switched splice junction usage.**

**a**, Genome browser view of the *AP1S2* locus. Tracks include the finalized SALA transcriptome alongside GENCODE v39 gene and transcript annotations. Switched isoform track displays only transcript models with significant changes in expression fraction between any pair of the three cell types (determined by isoformSwitchAnalyzeR). Color scale represents CPM values normalized within each cell type (from bambu). CFC-seq tracks display the summed signal of 5′ ends, 3′ ends, and exons across all samples. Tracks for ATAC-seq, cCRE promoter/enhancer regions, and SCAFE tCREs show genomic annotation at this locus. **b**, Isoform expression fractions of *AP1S2* in the three cell types, as estimated by isoformSwitchAnalyzeR. **c**, Predicted ORF alignment of the switched isoforms of *AP1S2*. **d**, Genome browser view of the *PORCN* locus with track structure as in (**a**). **e**, Isoform expression fractions of *PORCN* in the three cell types. **f**, Predicted ORF alignment of the switched isoforms of *PORCN*.

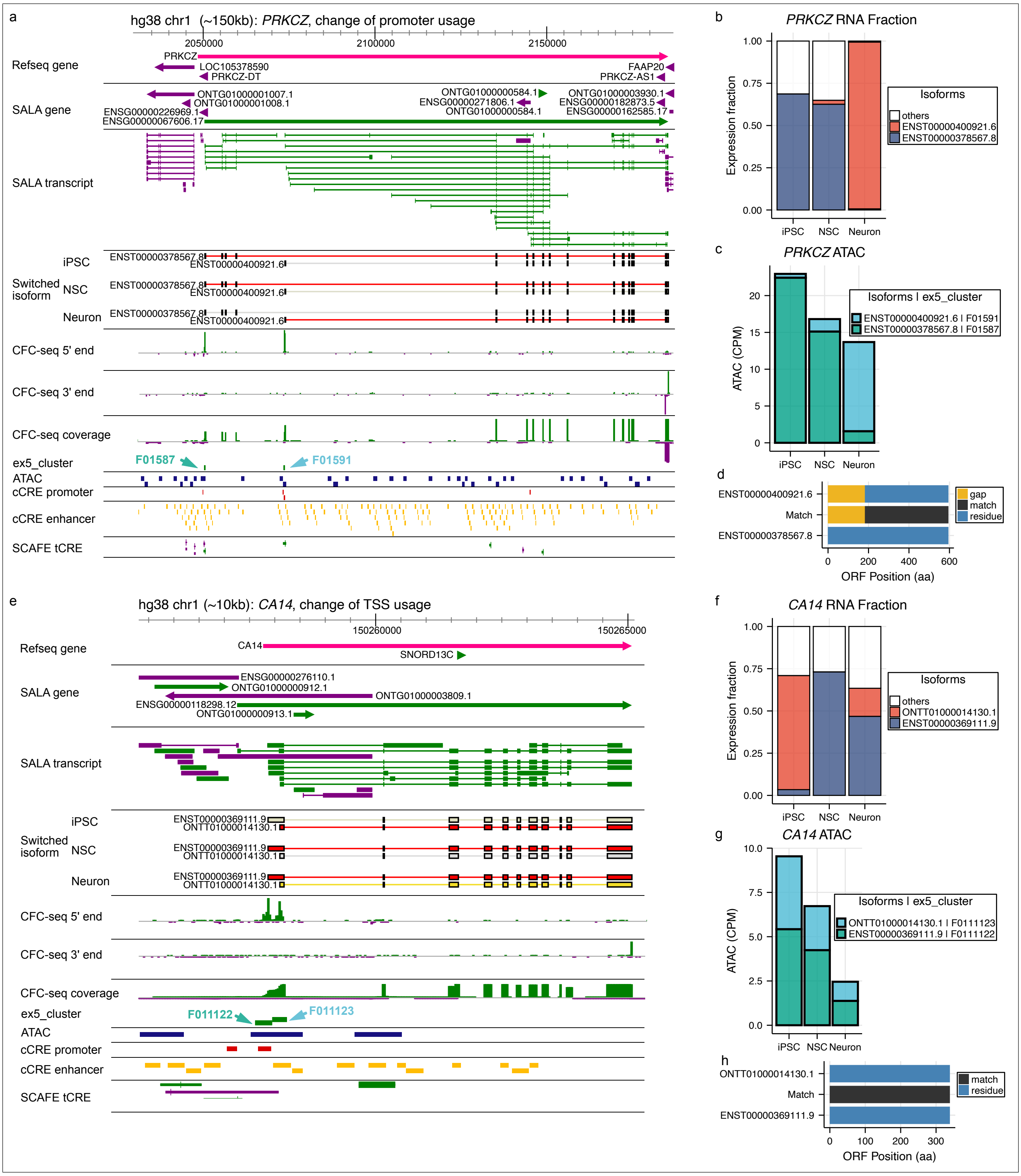


**Supplementary Figure 3 | Example genes with significant isoform switching due to switched promoter and TSS usage.**

**a**, Genome browser view of the *PRKCZ* locus. Tracks include the finalized SALA transcriptome alongside GENCODE v39 gene and transcript annotations. Switched isoform track displays only transcript models with significant changes in expression fraction between any two of the three cell types (as identified by isoformSwitchAnalyzeR). Color scale indicates CPM values derived from *bambu*, normalized within each cell type. CFC-seq tracks show the aggregated signal for 5′ ends, 3′ ends, and exons across all samples. Additional tracks show annotations from ATAC-seq, cCRE promoter and enhancer regions, and SCAFE-defined tCREs. **b**, Isoform expression fractions of *PRKCZ* in the three cell types, estimated by isoformSwitchAnalyzeR. **c**, Chromatin activity of corresponding ex5_clusters derived from ATAC-seq signal normalized by library size. **d**, Predicted ORF alignment of the switched isoforms of *PRKCZ*. **e**, Genome browser view of the *CA14* locus, track layout as in (a). **f**, Isoform expression fractions of *CA14* in the three cell types. **g**, Chromatin activity of ex5_clusters derived from ATAC-seq signal. **h**, Predicted ORF alignment of the switched isoforms of *CA14*.

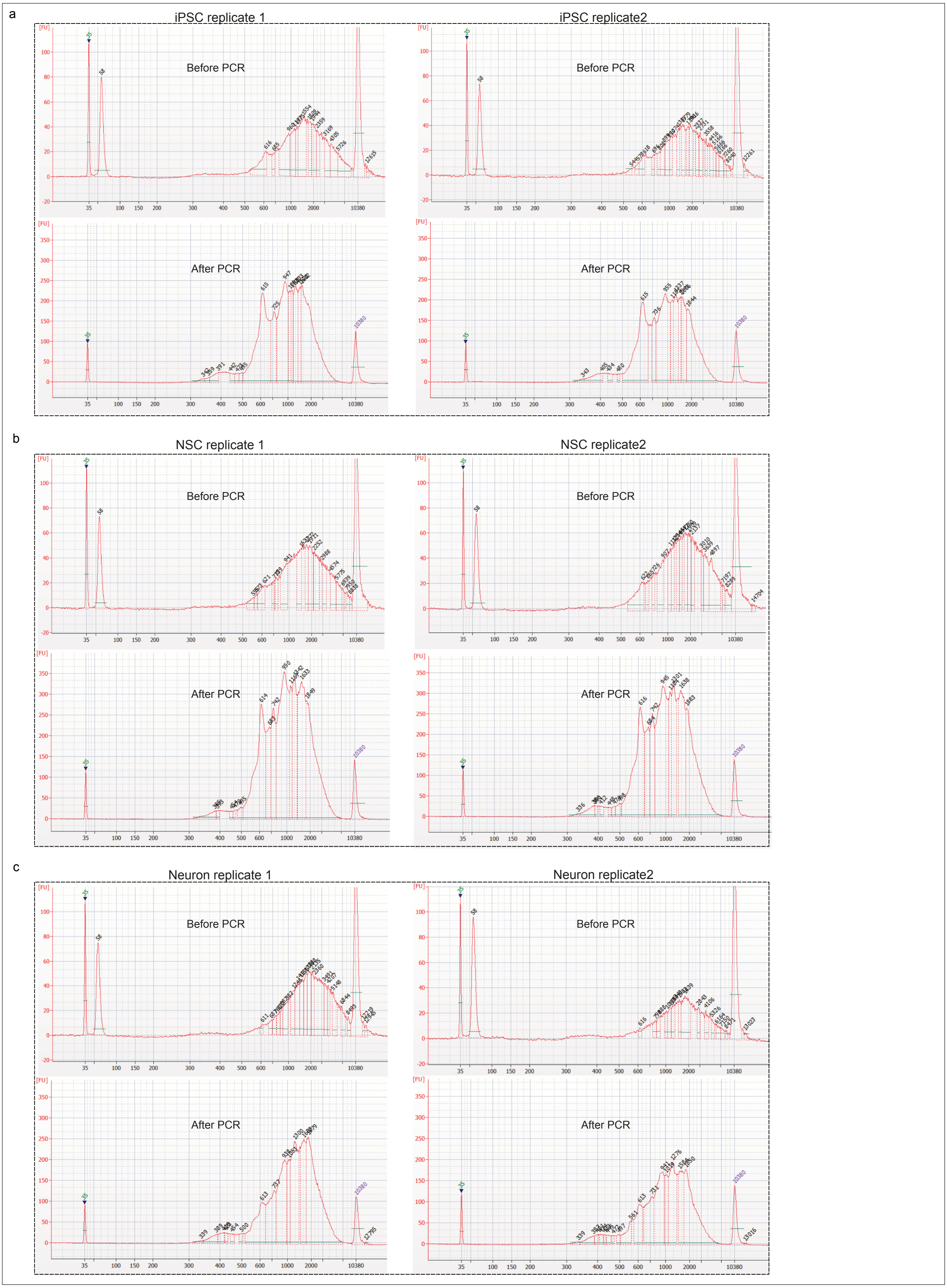


**Supplementary Figure 4 | Length distribution revealed by bioanalyser before and after PCR amplification.**

Length distribution before and after PCR and size selection for **a**, iPSC, **b**, NSC and **c**, Neuron.
