## Extended Figure 1-9 for "Genomic codes governing enhancer RNA fate"

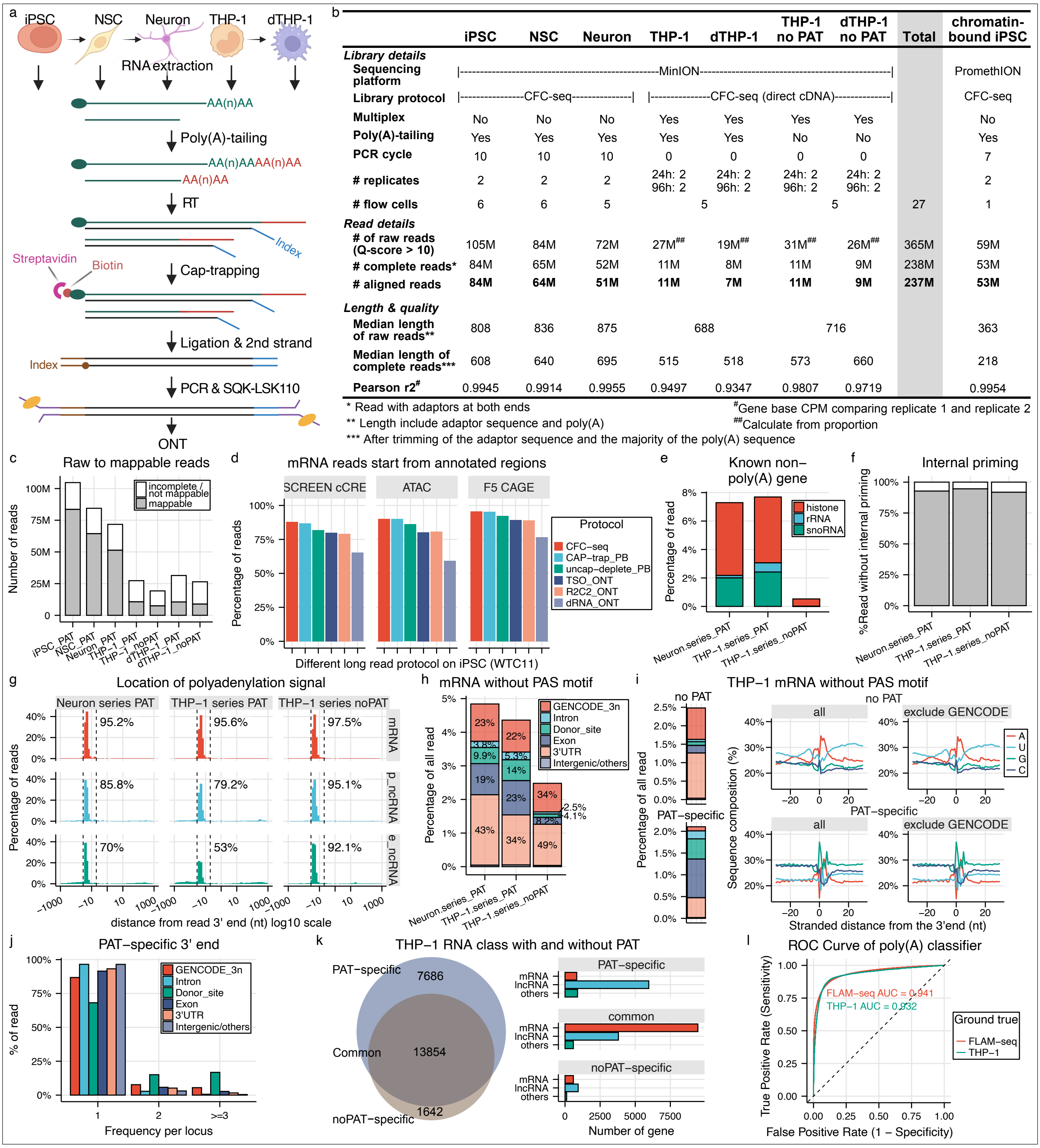


b

| | iPSC | NSC | Neuron | THP-1 | dTHP-1 | THP-1 no PAT | dTHP-1 no PAT | Total | chromatin-bound iPSC |
|---|---|---|---|---|---|---|---|---|---|
| ***Library details*** | | | | | | | | | |
| **Sequencing platform** | \|----------MinION---------- | | | | | | ----------\| | | PromethION |
| **Library protocol** | \|-----CFC-seq----- | | -----\| | \|-----CFC-seq (direct cDNA)----- | | | -----\| | | CFC-seq |
| **Multiplex** | No | No | No | Yes | Yes | Yes | Yes | | No |
| **Poly(A)-tailing** | Yes | Yes | Yes | Yes | Yes | No | No | | Yes |
| **PCR cycle** | 10 | 10 | 10 | 0 | 0 | 0 | 0 | | 7 |
| **# replicates** | 2 | 2 | 2 | 24h: 2<br>96h: 2 | 24h: 2<br>96h: 2 | 24h: 2<br>96h: 2 | 24h: 2<br>96h: 2 | | 2 |
| **# flow cells** | 6 | 6 | 5 | 5 | | 5 | | 27 | 1 |
| ***Read details*** | | | | | | | | | |
| **# of raw reads (Q-score > 10)** | 105M | 84M | 72M | 27M## | 19M## | 31M## | 26M## | 365M | 59M |
| **# complete reads*** | 84M | 65M | 52M | 11M | 8M | 11M | 9M | 238M | 53M |
| **# aligned reads** | **84M** | **64M** | **51M** | **11M** | **7M** | **11M** | **9M** | **237M** | **53M** |
| ***Length & quality*** | | | | | | | | | |
| **Median length of raw reads**** | 808 | 836 | 875 | 688 | | 716 | | | 363 |
| **Median length of complete reads***** | 608 | 640 | 695 | 515 | 518 | 573 | 660 | | 218 |
| **Pearson r2#** | 0.9945 | 0.9914 | 0.9955 | 0.9497 | 0.9347 | 0.9807 | 0.9719 | | 0.9954 |

* Read with adaptors at both ends
** Length include adaptor sequence and poly(A)
*** After trimming of the adaptor sequence and the majority of the poly(A) sequence
#Gene base CPM comparing replicate 1 and replicate 2
##Calculate from proportion

**Extended Data Figure 1 | Technical validation and read statistics of the CFC-seq platform.**

a, CFC-seq experimental workflow. b, Sequencing statistics and library metrics for all generated datasets. c, Quality control breakdown of long reads (base-calling score ≥ 10) into incomplete (missing adaptors), non-mappable, and mappable fractions. d, Intersection of mRNA 5′ ends with SCREEN cCREs, ATAC-seq peaks, and FANTOM5 CAGE clusters. Long-read datasets from iPSC WTC11 were obtained from ENCODE LRGASP (Pardo-Palacios et al. 2024). e, Enrichment of non-polyadenylated (non-poly(A)) transcripts in libraries following in vitro poly(A)-tailing (PAT). f, Evaluation of internal priming based on genomic sequence composition downstream of read 3′ ends. g, Polyadenylation signal (PAS) motif frequency (motif score >3, -35 to -5 bp upstream of the 3′ end) in reads across mRNA, promoter-ncRNA, and eRNA classes, comparing ±PAT libraries. h, Genomic distribution of non-PAS-associated mRNA 3′ ends to explore fraction of potential nascent RNA. i, Characterization of emergent mRNA 3′ ends lacking PAS motifs in THP-1 ±PAT libraries. Left: genomic localization relative to coding exons and introns. Right: nucleotide composition frequency surrounding 3′ end sites (±30 nt). j, Frequency of PAT-specific mRNA 3′ ends across genomic features, showing accumulation at splice donor sites. k, Overlap of genes detected by CFC-seq ±PAT and the distribution of their RNA classes. l, ROC curve for the sequence-based poly(A) classifier, validated against FLAM-seq 3′ ends (as positives) and PAT-specific 3′ ends (as negatives).

a

| | iPSC | NSC | Neuron | THP-1 | dTHP-1 | Total | iPSC | NSC | Neuron | Total |
|---|---|---|---|---|---|---|---|---|---|---|
| **Sequencing type** | CFC-seq | CFC-seq | CFC-seq | CFC-seq | CFC-seq | | CAGE | CAGE | CAGE | |
| **# of aligned reads** | 83,567,511 | 64,395,952 | 51,339,392 | 21,160,166 | 16,325,453 | 236,788,474 | 70,424,817 | 56,984,290 | 60,295,551 | 187,704,658 |
| **# SCAFE qualified reads** | 57,956,149 | 43,713,279 | 35,040,796 | 15,642,435 | 12,163,141 | 164,515,800 | 69,090,345 | 55,815,035 | 59,096,669 | 184,002,049 |
| **# CTSS with unencoded G** | 46,467,822 | 36,449,396 | 27,046,208 | 12,226,166 | 9,346,143 | 131,535,735 | 36,510,177 | 29,765,046 | 30,116,233 | 96,391,456 |
| **SCAFE per sequencing type** | | | | | | | | | | |
| **Included** | ✓ | ✓ | ✓ | ✓ | ✓ | | ✓ | ✓ | ✓ | |
| **# TSS cluster / # tCRE** | 159,116 / 73,328 | | | | | | 151,870 / 59,863 | | | |
| **# raw reads in cluster** | 73,479,071 | 55,618,351 | 44,704,247 | 17,952,524 | 13,984,123 | | 67,273,753 | 54,005,792 | 56,507,903 | |
| **# qualified reads in cluster** | 54,072,147 | 40,357,588 | 32,120,590 | 14,144,897 | 11,053,367 | | 66,682,910 | 53,470,499 | 56,028,124 | |
| **SCAFE for comparison** | | | | | | | | | | |
| **Included** | ✓ | ✓ | ✓ | | | | ✓ | ✓ | ✓ | |
| **# TSS cluster / # tCRE** | 239,141 / 84,320 | | | | | | | | | |
| **# raw reads in cluster** | 75,602,791 | 57,484,441 | 45,698,667 | | | | 67,195,194 | 53,940,922 | 56,596,952 | |
| **# qualified reads in cluster** | 55,305,641 | 41,460,241 | 32,766,720 | | | | 66,320,841 | 53,185,685 | 55,852,445 | |

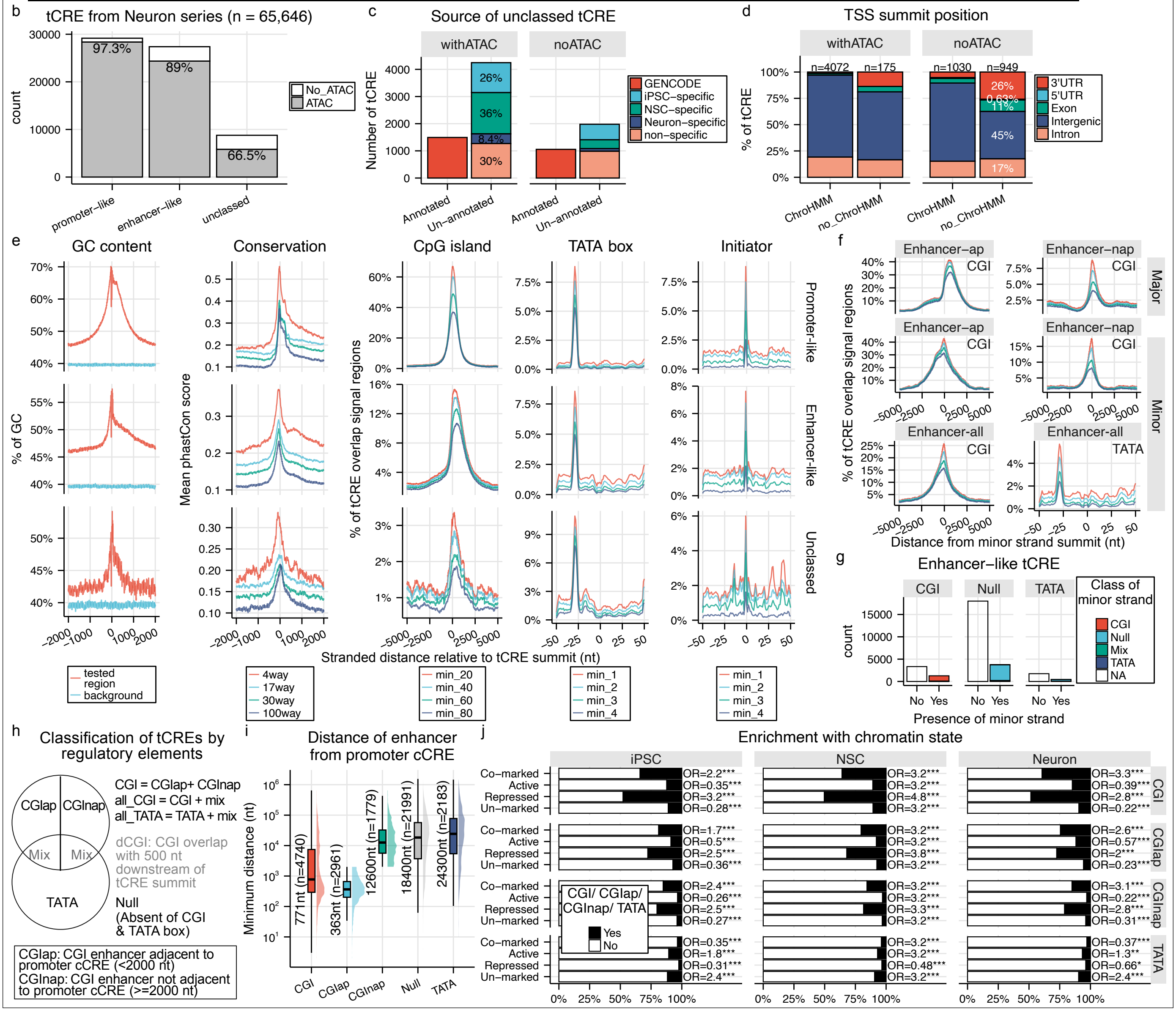


**Extended Data Figure 2 | Identification of genuine TSSs, clusters, and tCREs from 5′ ends of CFC-seq reads.**

**a**, Summary of reads processed with SCAFE for identification of genuine TSS clusters and tCREs. **b**, Neuron-series tCREs overlap with ATAC-seq-defined open chromatin from the same samples. **c**, Unclassed tCREs were stratified by ACTA-seq support, GENOCODE annotation status, and the originating cell type. **d**, Overlap of tCREs with chromatin states defined by ChromHMM using CUT&Tag data (H3K27ac, H3K27me3, H3K4me1, H3K4me3, CTCF) from three Neuron-series cell types. Colors indicate summit positions relative to GENCODE models. **e**, Genomic features of major-strand tCREs (n = 57,597) categorized as promoter-like, enhancer-like, or unclassed. Analyses include PhastCons conservation (4way–100way), overlap with GC content and CGI (UCSC), and motif enrichment (TATA box: MA0108.3, INR: POL002.1, from JASPAR). Summits were extended ±2 kb for conservation and GC content, and ±50 bp for motif analysis; backgrounds were exon- and tCRE-masked. **f**, Distribution of CGI and TATA box around summits of major-strand and minor-strand tCREs of all the enhancers and enhancers divided into adjacent to promoter (enhancer-AP) and not adjacent (enhancer-NAP). **g**, Proportion of enhancer-like major-strand tCREs with antisense minor-strand tCREs and their regulatory features. **h**, Classification and naming schema for regulatory elements. **i**, Distance of different classes of enhancer with their closest promoter cCRE. **j**, Enrichment of chromatin states with CGI, CGIap, CGInap, and TATA elements in enhancer-like tCREs across the 3 cell types. Enrichment test is performed by using Fisher's exact; $p < 0.05$ (*), $p < 0.01$ (**), $p < 0.001$ (***), n.s. = not significant.

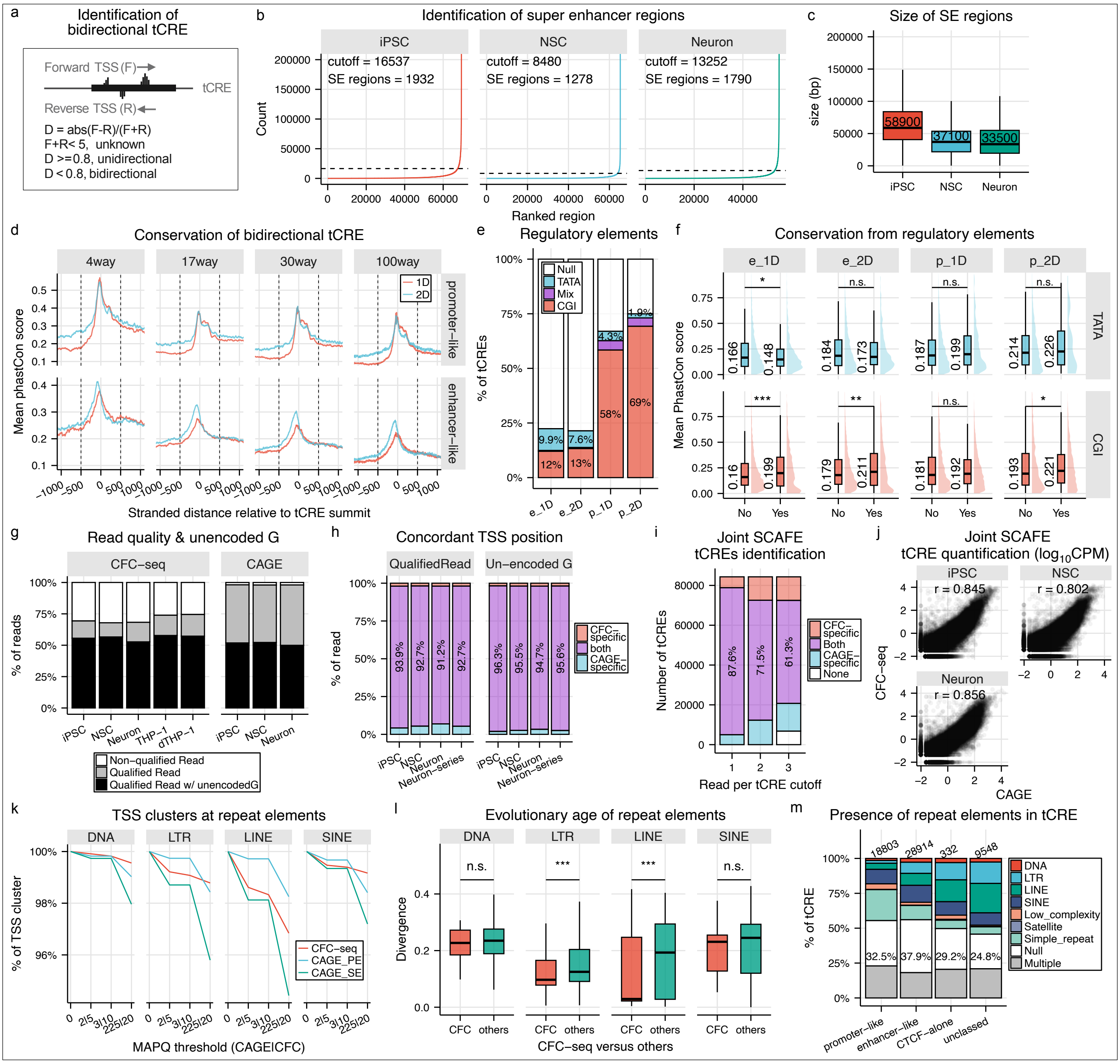


**Extended Data Figure 3 | Features of tCREs related to directionality, super enhancers and repetitive elements.**

**a**, Schematic illustrating the classification of tCREs as bidirectional (2D) or unidirectional (1D) based on transcription activity from both strands. **b**, Read count distribution of stitched H3K27ac peaks generated by ROSE. Regions exceeding the dotted line threshold were classified as super enhancers. **c**, Size distribution of super enhancers identified in the three analyzed cell types. **d**, Sequence conservation around tCRE summits for 2D and 1D promoters and enhancers, based on phastCons scores as described in **Extended Data Figure 2e**. **e**, Proportion of 2D and 1D tCREs (major strand only) containing CpG islands (CGI), TATA boxes, or both (Mix). **f**, Sequence conservation (phastCons 17-way scores) around the summit (±500 nt) of 2D and 1D, promoter-like and enhancer-like tCREs, stratified by presence of CGI or TATA box. Only major strand tCREs were included. **g**, Classification of mapped reads analyzed by SCAFE from CFC-seq and CAGE datasets into: not qualified (5′ soft-clipping >3 nt), qualified without unencoded G, and qualified with unencoded G. **h**, Distribution of single-nucleotide TSSs found uniquely in CFC-seq, in both CFC-seq and CAGE, or uniquely in CAGE. **i**, Number of tCREs detected by CFC-seq, CAGE, or both at varying read count thresholds. tCREs were identified by joint SCAFE analysis of Neuron series CFC-seq and CAGE datasets. **j**, Expression correlation per cell type between CFC-seq and CAGE, shown as RLE-normalized CPM values for each tCRE (each dot represents one tCRE). tCREs were identified by joint SCAFE analysis. **k**, TSS cluster identification coverage by CFC-seq, CAGE paired-end (PE), and CAGE single-end (SE) across increasing MAPQ thresholds. Coverage is shown relative to union TSS clusters at MAPQ = 0. **l**, Repetitive element divergence (as a proxy for evolutionary age) for repeats overlapping CFC-specific TSS clusters at MAPQ 20 versus others. **m**, Proportion of major strand tCREs containing repetitive elements, colored by repeat class. “Multiple” indicates overlap with more than one repeat class. Unless specified, statistical tests were two-sided Wilcoxon; $p < 0.05$ (*), $p < 0.01$ (**), $p < 0.001$ (***), n.s. = not significant.

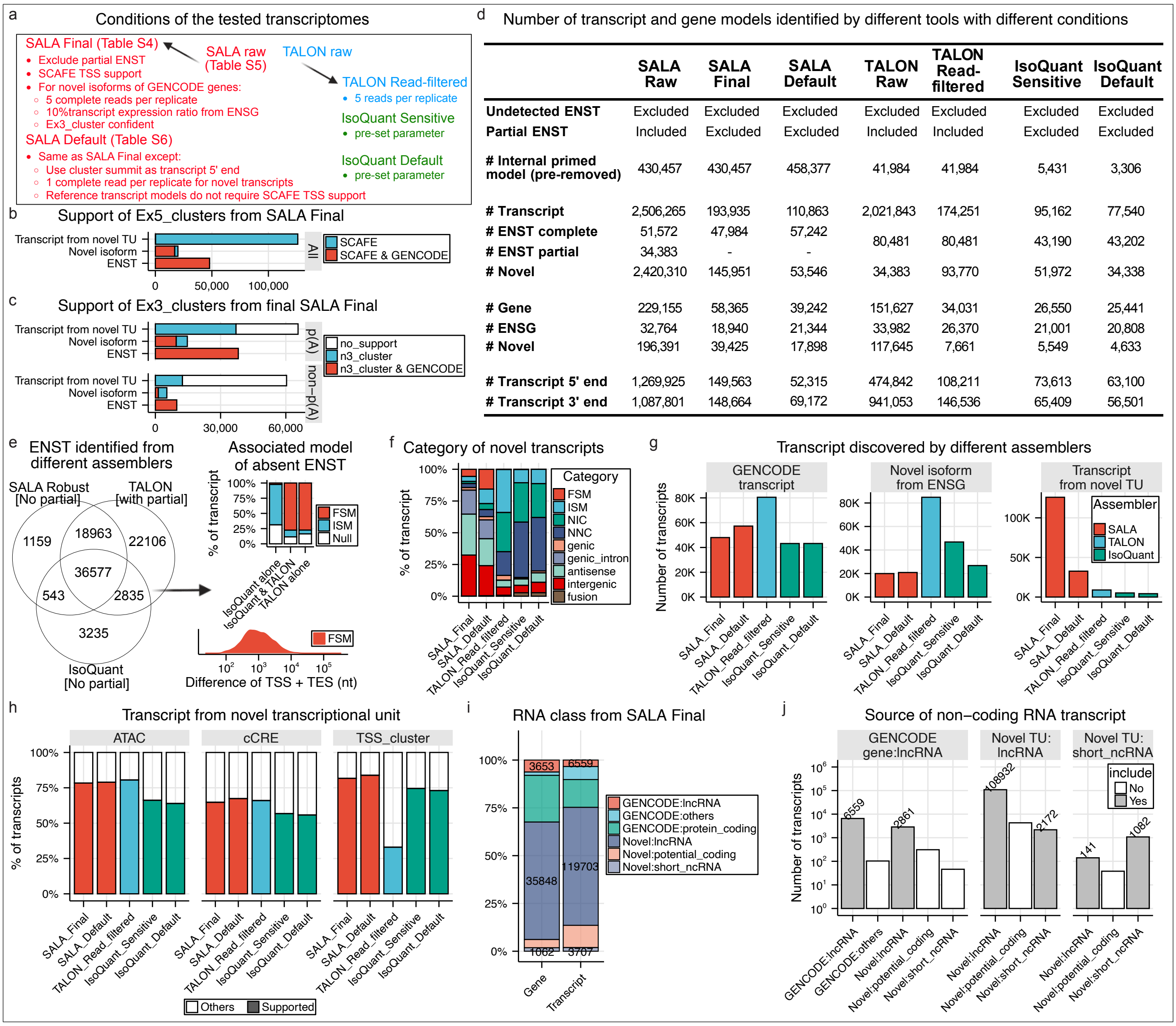


| | SALA Raw | SALA Final | SALA Default | TALON Raw | TALON Read-filtered | IsoQuant Sensitive | IsoQuant Default |
|---|---|---|---|---|---|---|---|
| **Undetected ENST** | Excluded | Excluded | Excluded | Excluded | Excluded | Excluded | Excluded |
| **Partial ENST** | Included | Excluded | Excluded | Included | Included | Excluded | Excluded |
| **# Internal primed model (pre-removed)** | 430,457 | 430,457 | 458,377 | 41,984 | 41,984 | 5,431 | 3,306 |
| **# Transcript** | 2,506,265 | 193,935 | 110,863 | 2,021,843 | 174,251 | 95,162 | 77,540 |
| **# ENST complete** | 51,572 | 47,984 | 57,242 | 80,481 | 80,481 | 43,190 | 43,202 |
| **# ENST partial** | 34,383 | - | - | | | | |
| **# Novel** | 2,420,310 | 145,951 | 53,546 | 34,383 | 93,770 | 51,972 | 34,338 |
| **# Gene** | 229,155 | 58,365 | 39,242 | 151,627 | 34,031 | 26,550 | 25,441 |
| **# ENSG** | 32,764 | 18,940 | 21,344 | 33,982 | 26,370 | 21,001 | 20,808 |
| **# Novel** | 196,391 | 39,425 | 17,898 | 117,645 | 7,661 | 5,549 | 4,633 |
| **# Transcript 5' end** | 1,269,925 | 149,563 | 52,315 | 474,842 | 108,211 | 73,613 | 63,100 |
| **# Transcript 3' end** | 1,087,801 | 148,664 | 69,172 | 941,053 | 146,536 | 65,409 | 56,501 |

**Extended Data Figure 4 | Transcript and gene models identified by SALA.**

**a**, Number of transcript models in the finalized transcriptome with 5′ end support from either SCAFE or GENCODE annotations. **b**, Number of transcript models in the finalized transcriptome with 3′ end support. Poly(A)-positive (P(A)) transcripts were defined based on a positive poly(A) classifier score or the presence of a polyadenylation signal (PAS) with motif score >3. **c**, Summary of filtering criteria applied to generate different transcriptome versions for benchmarking. **d**, Table comparing the number of transcript and gene models identified by SALA and other transcriptome assembly tools under different filtering conditions. **e**, Venn diagram showing the overlap of detected GENCODE transcript (ENST) across different software tools. Right panel shows the category of transcript model associated to the ENST observed from other tools but not in SALA. Only Full Splice Match (FSM) and Incomplete Splice Match (ISM) were shown. The sum of the distance difference from the two ends between the FSM transcript models and associated ENST was shown in the distribution plot. **f**, Transcript categories assigned to novel transcript models using SQANTI3 across different software outputs. FSM: Full Splice Match, ISM: Incomplete Splice Match, NIC: Novel in Catalog, NNC: Novel Not in Catalog. **g**, Number of transcript models classified into three categories: “GENCODE transcript,” “Novel isoform from ENSG,” and “Transcript from novel transcriptional unit (TU).” **h**, Proportion of “Transcript from novel TU” models whose 5′ ends overlap with open chromatin (ATAC-seq), ENCODE SCREEN candidate *cis*-regulatory elements (cCREs), or SCAFE genuine TSS clusters. **i**, Gene and transcript class assignments in the SALA Final transcriptome. **j**, Number of ncRNA transcript models included in three groups (GENCODE genes, lncRNA from novel TU, and short_ncRNA from novel TU) according to criteria used to define non-coding RNA classes in this study.

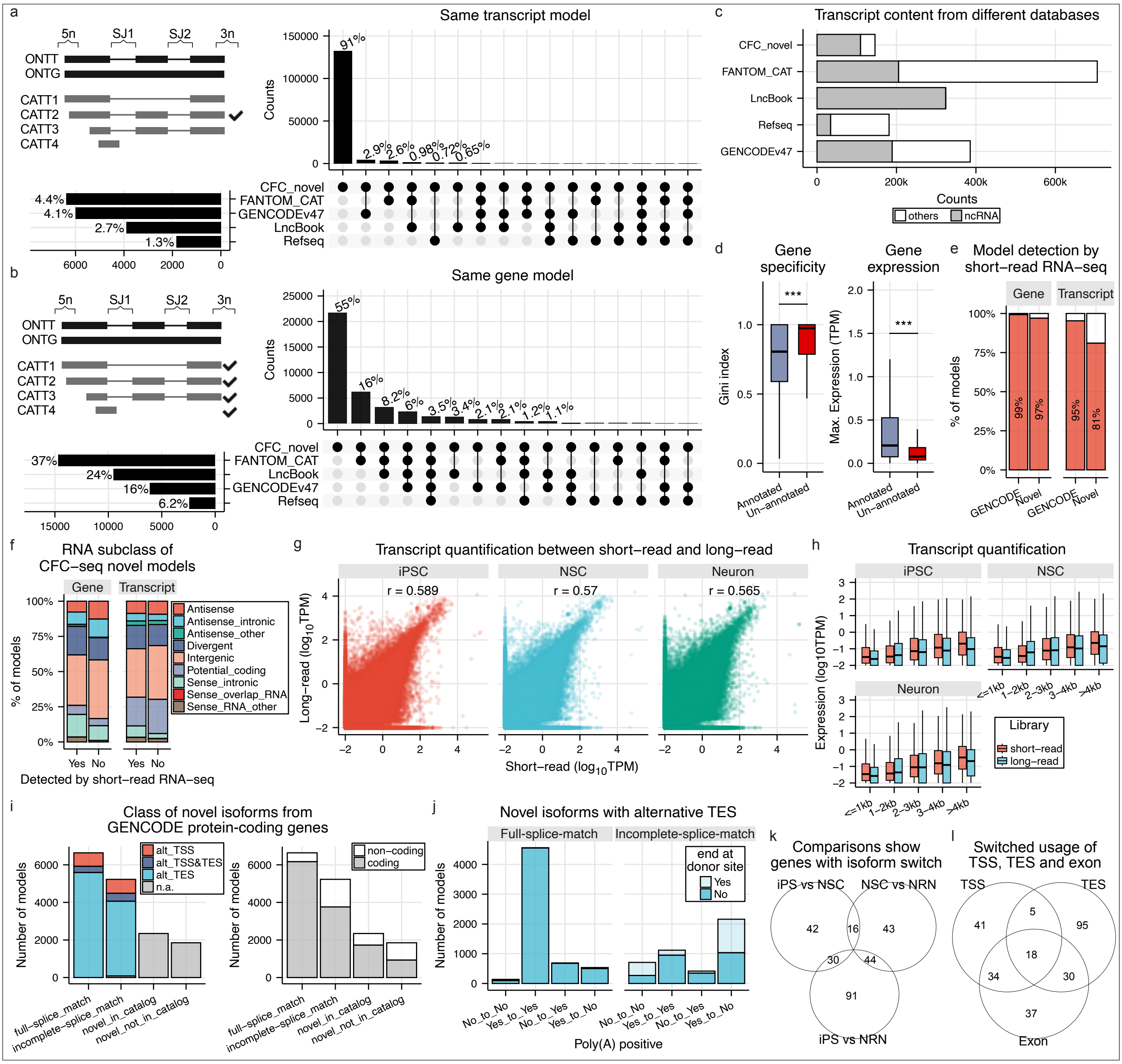


**Extended Data Figure 5 | Transcript and gene models identified by SALA.**

**a**, UpSet plot showing novel transcript models identified in this study and their overlap with FANTOM CAT, LncBook, RefSeq, and GENCODE v47. Transcript model identity was determined by SALA; schematic shown on the left. **b**, UpSet plot of novel gene models and their overlap with the same four databases, as identified by SALA. **c**, Counts of transcript models annotated as ncRNA or other classes across the four databases. **d**, Cell-type specificity (Gini index) of gene models annotated in any of the public databases or completely novel, across the five studied cell types. Right, maximum expression level across the five cell types for each group. **e**, Proportion of gene and transcript models in the final CFC-seq transcriptome also detected by short-read RNA-seq in iPSC, NSC, and Neuron. **f**, Classification of novel gene and transcript models (detected or undetected by short-read RNA-seq) into ncRNA subclasses. **g**, Correlation of expression levels from long- and short-read data using the finalised transcriptome, limited to detectable GENCODE models. **h**, Expression levels of transcript models from short- and long-read libraries, grouped by transcript length. **i**, Classification of novel isoforms of protein-coding genes. "Full-splice match" indicates isoforms with all splice junctions matching a GENCODE transcript; "incomplete-splice match" includes partial matches. Alternative TSS and TES was assessed by ex5_clusters and ex3_clusters relative to best-matched GENCODE transcripts; skipped for unmatched cases (novel in catalog or novel not in catalog). Right, predicted coding potential by CPAT. **j**, For novel isoforms with alternative TES, presence of poly(A) signals in the isoform and the matched GENCODE transcript was assessed via PAS motif and poly(A) predictions. Isoforms were further stratified by whether their 3′ end overlaps a splice donor site. **k**, Venn diagram showing genes with both significantly up- and down-regulated isoforms across three comparisons. **l**, Venn diagram of genes with isoform switching involving alternative TSS, TES, and exon usage.

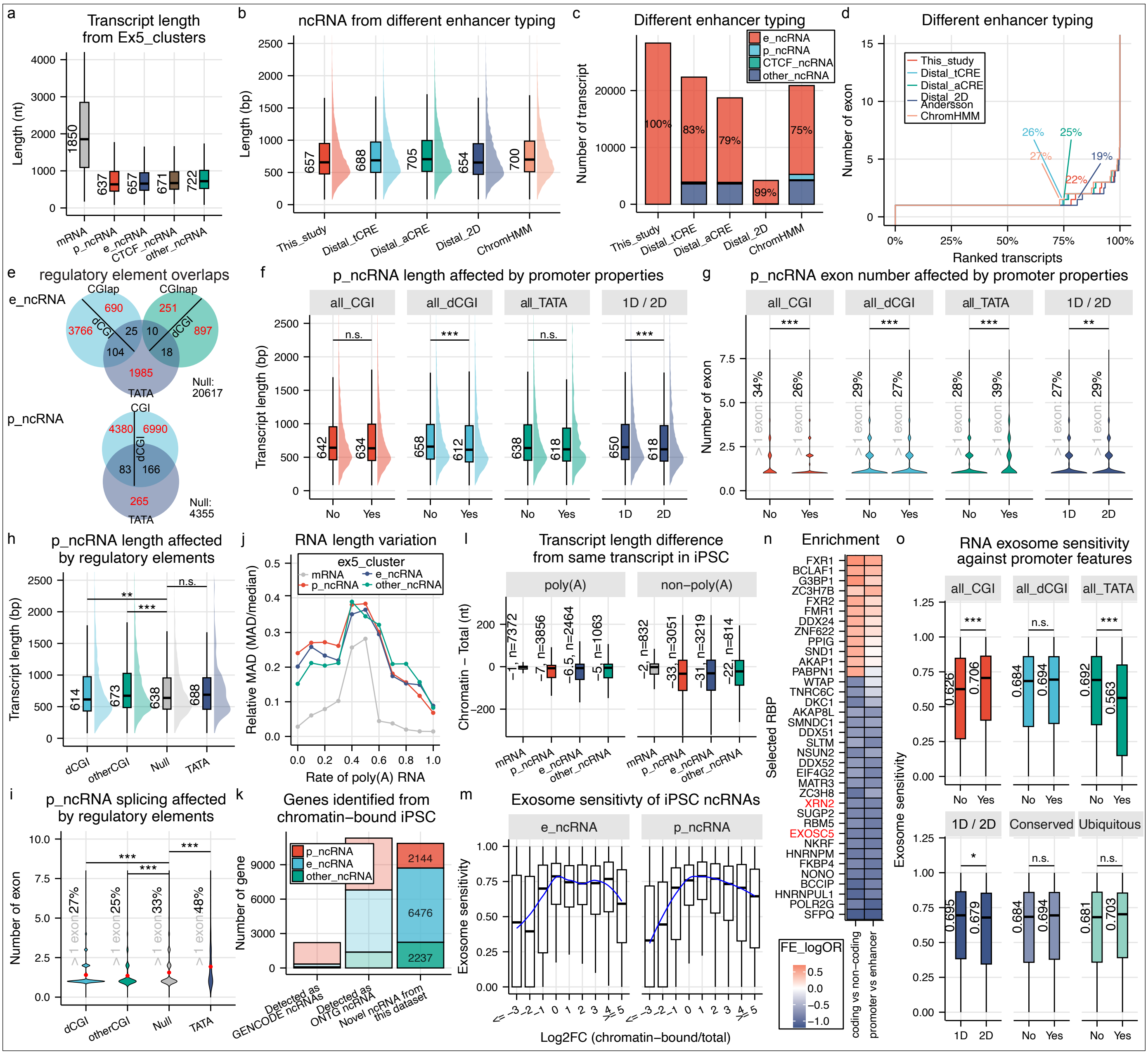


**Extended Data Figure 6 | Factors influencing RNA transcription.**

**a**, Transcript lengths of RNA groups defined by ex5_clusters. **b**, Transcript length of eRNAs defined using various enhancer annotation methods. **c**, Proportion of RNA groups identified in this study among enhancers defined by different methods. **d**, Median exon number of eRNAs by enhancer definition method; percentages indicate the proportion with a median exon count > 1. **e**, Venn diagrams showing overlap of ex5_clusters across TATA, CGIap, and CGInap, and the subset with downstream CGI (dCGI) for e_ ncRNA. The p_ncRNA was divided into TATA and CGI with dCGI subset. **f**, Transcript length of p_ncRNAs compared across promoter features (as defined in **Figure 4c**). **g**, Exon count of p_ncRNAs across promoter features; percentages indicate groups with more than one exon. **h**, Transcript length of four mutually exclusive p_ncRNA groups, based on individual promoter features. **i**, Median exon number for the four mutually exclusive p_ncRNA groups; percentages indicate groups with more than one exon. **j**, Length variation within ex5_clusters represented by relative MAD (MAD/median); plotted alongside poly(A) RNA rate (defined in **Figure 4k**). **k**, SALA analysis of chromatin-bound CFC-seq from iPSCs using the finalized transcriptome plus GENCODE v39. Shown are ncRNA genes originating from GENCODE, the novel ncRNA identified from the finalized transcriptome (ONTG), or the chromatin-bound library alone. **l**, Transcript length differences for shared transcript models between the chromatin-bound and main datasets, grouped by poly(A) status in the main dataset. **m**, Exosome sensitivity of transcript models (1,935 e_ncRNAs and 3,841 p_ncRNAs) with CPM > 0.25, based on RNA-seq of EXOSC3 knockdown in iPSC. Sensitivity ($log_2$ fold change) was estimated using edgeR on chromatin-bound vs. total RNA CFC-seq (quantified by bambu with the SALA finalized transcriptome updated with chromatin-bound models). **n**, Enrichment of individual RBP interactions from exons of coding vs. non-coding, and enhancer- vs. promoter-derived transcripts. **o**, Enrichment of exosome sensitivity in p_ncRNAs across different promoter features (defined in **Figure 4n**). Unless specified, statistical tests were two-sided Wilcoxon; $p < 0.05$ (*), $p < 0.01$ (**), $p < 0.001$ (***), n.s. = not significant.

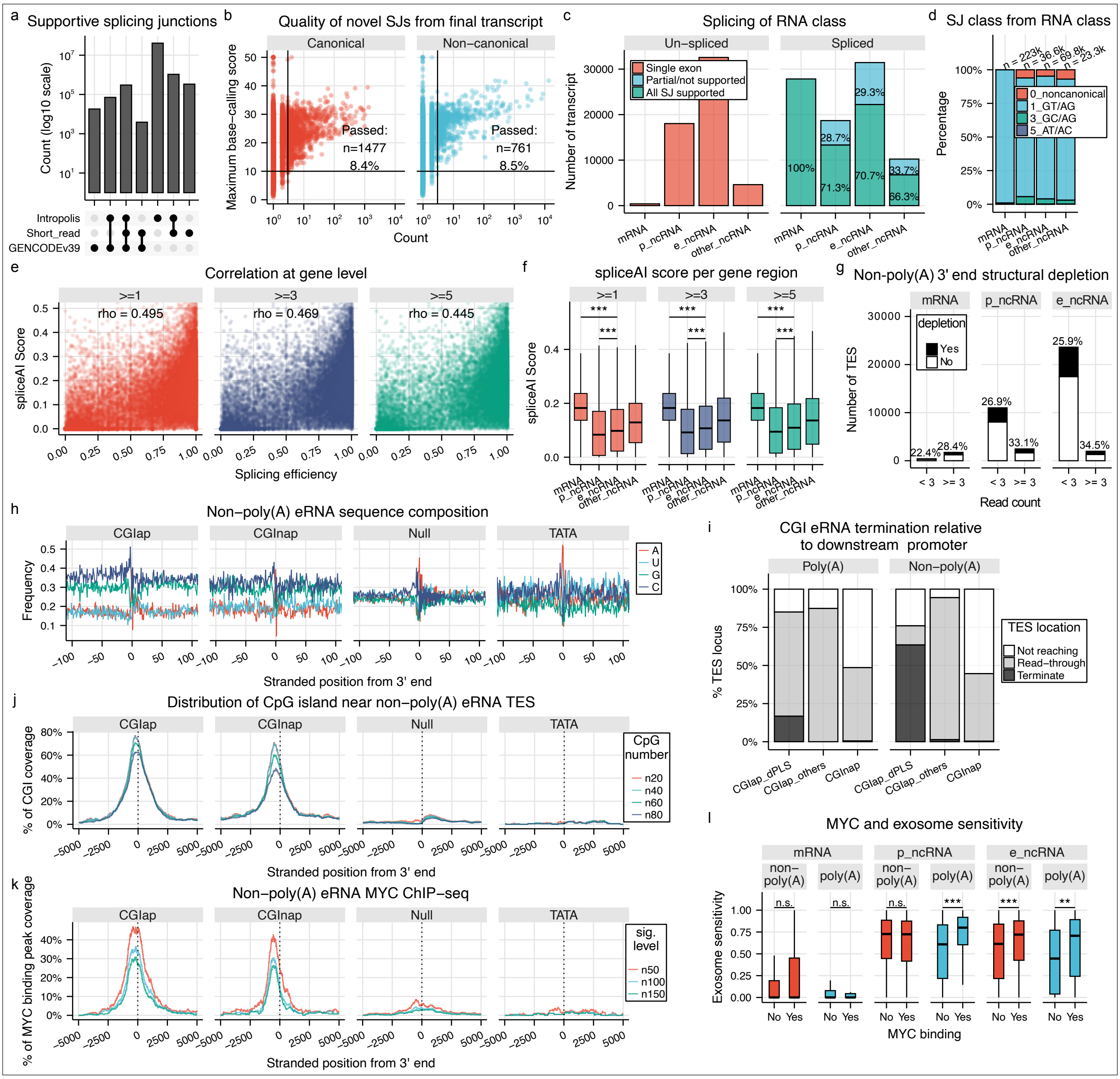


**Extended Data Figure 7 | Splicing efficiency and eRNA transcription termination.**

**a**, Number and percentage of splice junction (SJ) classes of all the SJs from external reference databases. “Short read” refers to confident SJs derived from short-read RNA-seq in the Neuron series. **b**, Novel SJs absent from the databases in (a) were classified as canonical or non-canonical. Plots show the relationship between the mean maximum base-calling score across the 6 nt surrounding each SJ and the SJ’s occurrence. SJs were considered technically confident if all 6 nt scored ≥10 and occurred ≥3 times. **c**, SJs from the final transcriptome were assigned to the four RNA classes, allowing for overlaps. For each class, the percentage of transcripts with and without fully supported SJs is shown. Positive SJ support was defined as either present in the databases in (**a**) or technically confident per (**b**). **d**, Total number of transcript models containing ≥1 SJ and percentage of each SJ class based on motif composition. **e**, Gene-level splicing efficiency compared to SpliceAI score across different occurrence thresholds. **f**, SpliceAI scores were summarized at the gene level and normalized by intron count. Gene models without SJs were excluded. SJs were filtered using three occurrence thresholds (sum of span and splice counts at both donor and acceptor sites). **g**, Evidence of RNA structural depletion (fold change >15%) at non-polyadenylated TESs, including both recursive and non-recursive TESs, based on RNAfold-derived hairpin scores. **h**, Nucleotide composition surrounding recursive transcription end sites (TESs; ≥3 read support) of non-poly(A) eRNAs, stratified by regulatory elements. **i**, Position of CGI enhancer-derived RNA TESs relative to the downstream promoter (SCREEN promoter cCRE). Not Reaching: TES is located > 500nt upstream the promoter region, Terminate: TES is located within 500nt upstream or inside the promoter, Read-through: TES is located downstream of the promoter. **j**, Distribution of CpG islands proximal to TESs of non-polyadenylated RNA types, restricted to recursive TESs. **k**, Distribution of MYC binding sites (from ChIP-seq dataset GSM1505809) near TESs of non-polyadenylated RNA types, restricted to recursive TESs. **l**, Exosome sensitivity of transcript models was weighted and grouped by TES and compared between TES with and without MYC binding, stratified as different classes. Unless specified, statistical tests were two-sided Wilcoxon; $p < 0.05$ (*), $p < 0.01$ (**), $p < 0.001$ (***), n.s. = not significant.

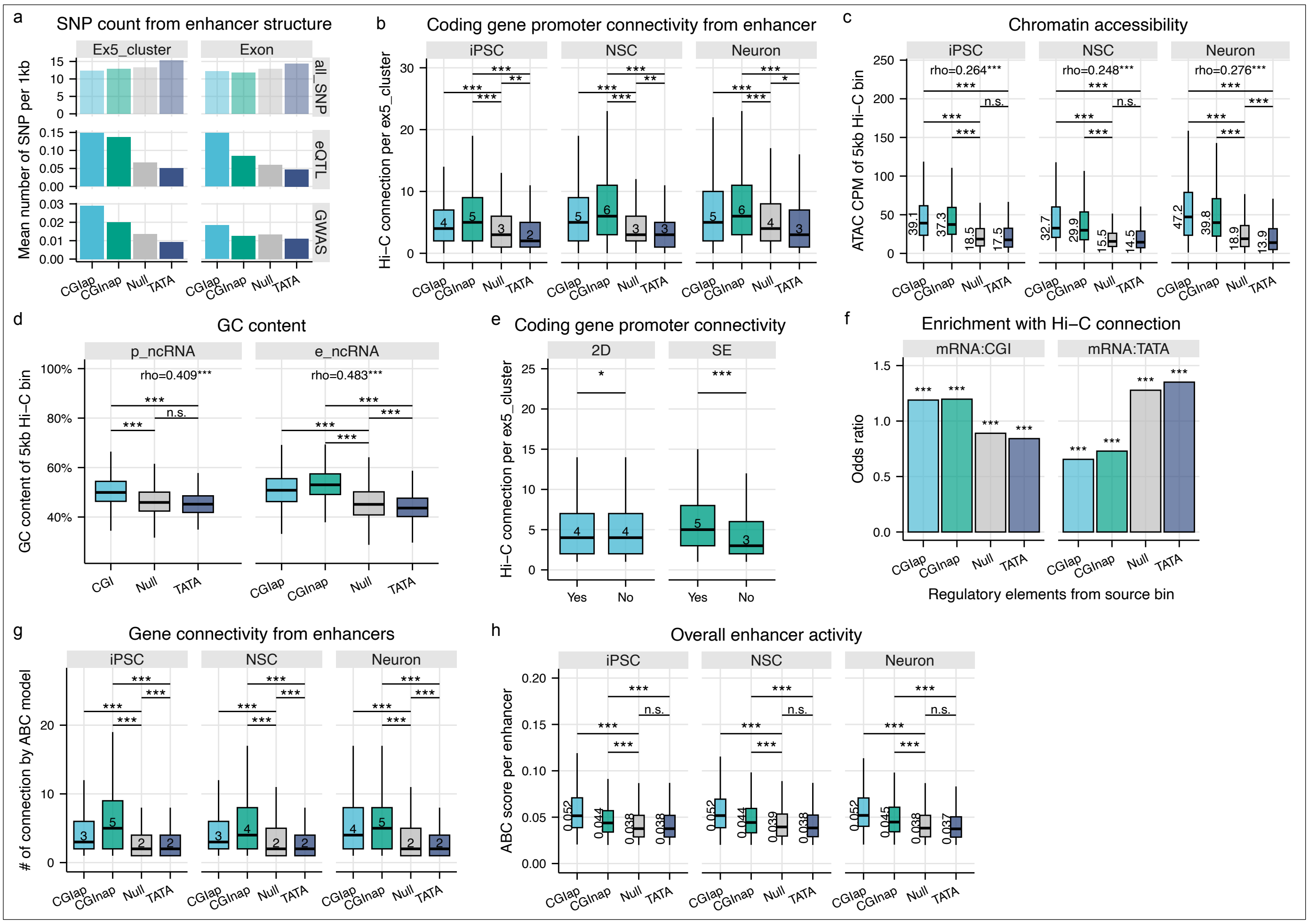


**Extended Data Figure 8 | Genetic variants and enhancer connectivity.**

**a**, Number of SNPs associated with e_ncRNAs grouped by mutually exclusive regulatory elements of ex5_clusters. SNP counts were normalized by the length of the ex5_cluster or transcript model (per kb). The "all_SNP" catalog refers to common SNPs from the NIH dataset. **b**, Hi-C interactions (5 kb resolution, FDR < 0.01, ≥5 reads) were used to infer ncRNA–mRNA connectivity via ex5_clusters. Only e_ncRNA ex5_clusters with ≥1 connection were included. Bins containing ex5_clusters with multiple regulatory elements (CGI and TATA box) were excluded. Median connection counts per ex5_cluster are shown, analyzed separately for each of the three cell types. **c**, Chromatin accessibility from matching ATAC-seq samples was mapped to 5 kb bins containing a single regulatory element (multi-element bins excluded). Median read counts and Spearman correlation between chromatin accessibility and Hi-C connectivity (as in panel **b**) are shown. **d**, GC content was calculated for 5 kb bins as in (**c**), and Spearman correlation with Hi-C connectivity (as in **Figure 6d**) is shown. **e**, Hi-C connectivity across all three cell types was compared against tCRE directionality and super-enhancer (SE) clustering. **f**, Two-sided Fisher's exact test was used to assess enrichment across regulatory element combinations. ncRNA source was grouped as in panel **b**; mRNA promoters were classified by presence or absence of CGI or TATA box. Odds ratios are shown for each combination. **g**, Gene connectivity of enhancers estimated by ABC-model. Analysis was performed on merged tCREs and linked to the features of ex5_clusters. Merged tCREs containing ex5_clusters with multiple regulatory elements (CGI and TATA box) were excluded. Only the enhancers with at least one connection with ABC score > 0.02 were included. **h**, Same analysis as (**g**) showing the mean ABC score for each enhancer. Unless specified, statistical tests were two-sided Wilcoxon; $p < 0.05$ (*), $p < 0.01$ (**), $p < 0.001$ (***), n.s. = not significant.

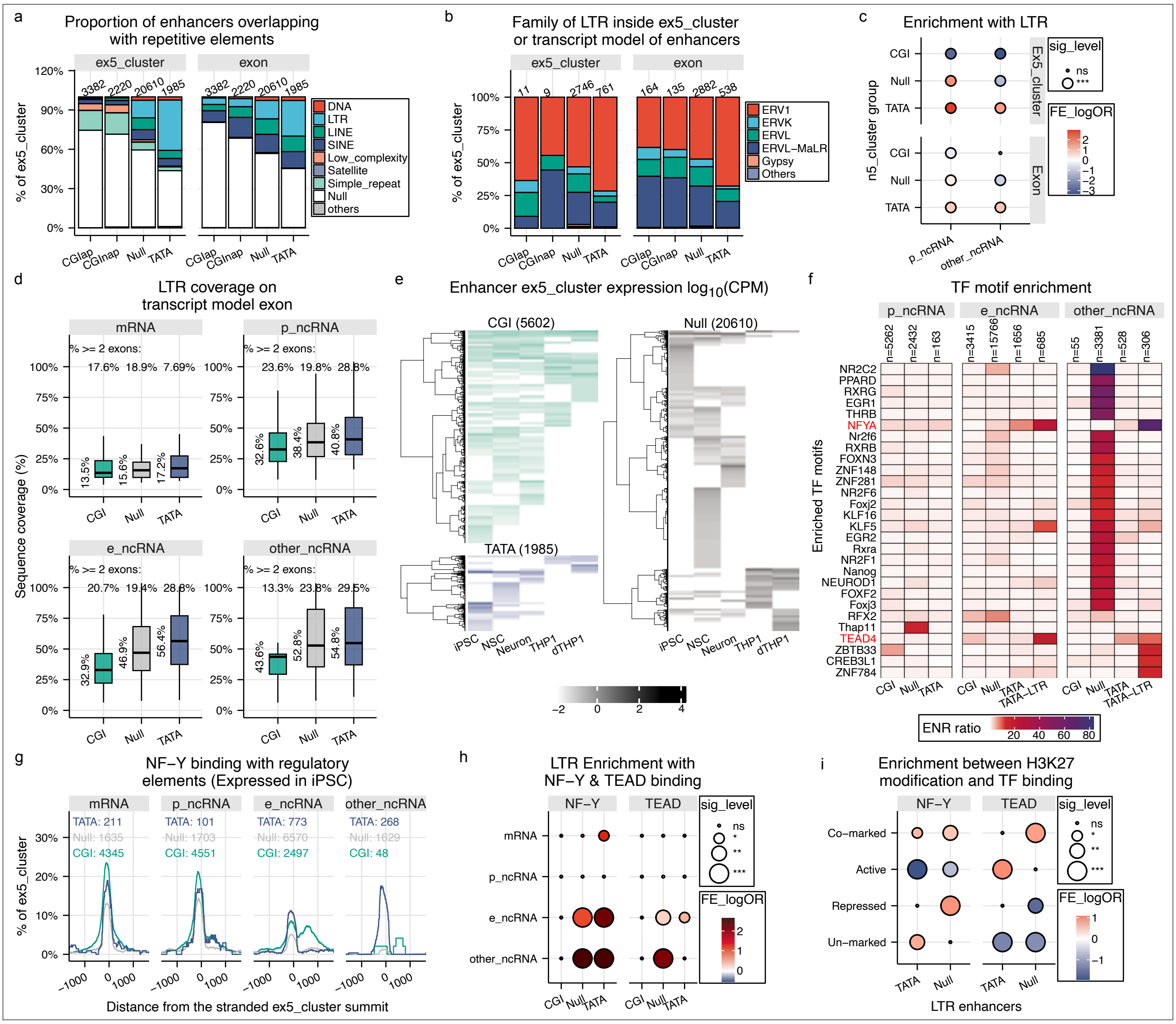


**Extended Data Figure 9 | TATA box enhancers enriched with LTR element are activated by NF-Y.**

**a**, Proportion of ex5_clusters and exons (grouped by cluster) intersecting with repetitive elements, applying the same criteria as in **Figure 6e**. **b**, Family composition of LTR elements intersecting enhancers and eRNAs. **c**, Enrichment with LTR across different groups in p_ncRNA and other_ncRNA was tested using Fisher's exact test. **d**, LTR coverage percentage over exons of transcript models that intersected with LTR, grouped by promoter type and regulatory element. Percentages on the top indicate the proportion of transcript models with LTR overlap across ≥2 exons. **e**, Expression profile of e_ncRNA ex5_clusters across five cell types, grouped into CGI, Null, and TATA categories. **f**, Enrichment ratio of transcription factor motifs was shown for those with FDR < 0.05 and enrichment ratio >12 and expression levels >1 CPM. Motif analysis was performed on non-redundant 501 nt CRE sequences (summit +100 nt & -400 nt). **g**, Distribution of NF-Y binding across ex5_cluster summits, grouped into mRNA, e_ncRNA, p_ncRNA, and other_ncRNA and sub-grouped by CGI, Null, and TATA. The total number of ex5_clusters in each group is shown. **h**, Enrichment between LTR and binding of NF-Y and TEAD considering only ex5_cluster transcribed in iPSC. Fisher's exact derived odds ratio and p-values of the 12 subgroups were shown. **i**, Enrichment between chromatin states and binding of NF-Y and TEAD. Fisher's exact derived odds ratio and p-values were shown from TATA and Null subgroup of e_ncRNA ex5_clusters transcribed from iPSC, including only LTR$^{+}$. Co-marked (H3K27ac$^{+}$ & H3K27me3$^{+}$), Active (H3K27ac$^{+}$), Repressed (H3K27me3$^{+}$), Un-marked (H3K27ac$^{-}$ & H3K27me3$^{-}$). Unless specified, statistical tests were two-sided Wilcoxon; $p < 0.05$ (*), $p < 0.01$ (**), $p < 0.001$ (***), n.s. = not significant.
