## Supplementary material for "Genomic codes governing enhancer RNA fate": Online Methods

### Section 1 | Experimental workflows

#### Cell models

Human iPS line i$^3$N and THP-1 cells were used in this study. iPS line i$^3$N is a gift from Dr. Michael Ward from NIH and derived from the WTC11 iPSC by introducing a doxycycline-inducible mNGN2 transgene at the AAVS1 site. THP-1 cells were subcloned by limited dilution and one clone (1-E9) was selected for relatively high sensitivity to PMA. The ability to differentiate relatively homogeneously in response to PMA was evidenced by expression of CD14 and CSF-1R quantified by ssCAGE.

#### Cell culture

The iPS cells were cultured in StemFit medium (Ajinomoto; Cat. No. RCAK02N) under feeder-free conditions at 37 °C in a 5% $CO_2$ incubator. The cells were passaged when the culture was ~80% confluent. The cells were washed with DPBS and detached by incubating with Accutase (Sigma; Cat.No. A6964) at 37 °C for 10 min. After centrifuge, cells were resuspended in StemFit medium supplemented with 10 nM Rho-associated kinase (ROCK) inhibitor (Y-27632, Wako; Cat.No. 036-24023), and re-plated onto iMatrix-511 (Nippi; Cat.No. 892012)-coated plates. The following day, ROCK inhibitor was withdrawn by refreshing the StemFit medium. The medium was refreshed every other day. To prepare the cells for transcriptomic profiling, iPSC was plated at a density of 0.5 million cells per 10 cm-dish in 10mL of medium and collected by Accutase after four days. The THP-1_1-E9 cells were cultured in RPMI1640-GlutaMAX-I (Gibco) supplemented with 10% heat inactivated fetal bovine serum (Gibco), 100 U/mL penicillin, and 100 µg/mL streptomycin (Gibco). The cells were maintained in a humidified 37℃ incubator with 5% $CO_2$. Both cell lines are free from mycoplasma contamination.

#### Neural differentiation from iPSC to Neural stem cell (NSC)

NSCs were generated by using the PSC Neural Induction Medium (NIM) (Gibco; Cat.No. A1647801). iPSCs were seeded onto an iMatrix511-coated 6-well plate at a density of 0.25 million per well in 2 mL of StemFit medium with 10 nM ROCK inhibitor. The following day, medium was replaced with 2.5 mL of NIM. NIM was refreshed every other day. On day 6 of neural induction, the NSCs (P0) were harvested by incubating with Accutase at 37 °C for 10 min and scraping. The cells were centrifuged at 300 x g for 5 min. The supernatant was aspirated and the cell pellet was resuspended with PSC Neural Expansion Medium (NEM) containing 10 nM ROCK inhibitor and seeded onto a iMatrix511-coated 10-cm dish at a density of one million cells per dish. The following day, the medium was refreshed to remove the ROCK inhibitor. Hereafter, NEM was replaced every other day. After 5 days, the expanded P1-NSCs were harvested and stocked with STEM-CELLBANKER (Nippon Zenyaku, Cat.No. CB045). To prepare the cells for transcriptomic profiling, the cryopreserved P1-NSCs were quickly thawed and cultured in NEM with 10 nM ROCK inhibitor on an iMatrix511-coated 10 cm-dish at a density of one million. After four days of culture, P2-NSCs were harvested by incubating with Accutase and scraping.

**Neural differentiation from NSC to cortical neuron**

The differentiation of cortical neurons from NSC using doxycycline-inducible mNGN2 was performed following the protocol developed in Dr. Peter Heutink's Lab (DZNE), with modifications. Briefly, the cryopreserved P1-NSCs were passaged once. P2-NSCs were then replated onto 0.1mg/mL Poly-L-ornithine (Sigma, Cat.No. P3655)-coated 10cm-dish at a density of $5 \times 10^6$ cells per dish in the differentiation medium I (50% DMEM/F12 (Gibco, Cat.No. 10565018) and 50% Neurobasal Medium (Gibco; Cat.No. 21103049) as the base, supplemented with 0.5 × Non-Essential Amino Acids (NEAA, Gibco; Cat.No. 11140050), 0.5 × GlutaMAX (Gibco; Cat. No.35050-061), 0.5 × N-2 Supplement (Gibco; Cat. No. 17502048), 0.5X B-27 Supplement (Gibco; Cat. No. 17504-044), 2.5 µg/mL human Insulin (Sigma; Cat.No. 19278), 2 µM DAPT (Wako; Cat.No. 043-33581), 50 µM 2-Mercaptoethanol (Gibco; Cat. No. 21985023), 2 µg/mL doxycycline hydrochloride (Sigma; Cat.No. D-9891), 5 µg/mL Mouse Laminin, and 10 nM ROCK inhibitor). The following day, the same medium without ROCK inhibitor was used to replace medium. On day 3, the medium was changed to differentiation medium II (50% DMEM/F12 and 50% Neurobasal Medium as the base, supplemented with 0.5 × Non-Essential Amino Acids, 0.5 × GlutaMAX, 0.5 × N-2 Supplement, 0.5 × B-27 Supplement, 2.5 µg/mL human Insulin, 10 µM DAPT, 50 µM 2-Mercaptoethanol, 2 µg/mL doxycycline hydrochloride, 0.5 µg/mL Mouse Laminin, 10 ng/ml brain-derived neurotrophic factor (BDNF, PeproTech; Cat.No. 450-02), 10 ng/ml glial cell-derived neurotrophic factor (GDNF, PeproTech; Cat.No. 450-10), 10 ng/ml neurotrophin-3 (NT-3, PeproTech; Cat.No. 450-03), and 0.5 µg/ml laminin). On Day 6 onwards, the half of the medium was exchanged every 3 days without Doxycycline. On day 10, the differentiated neurons were harvested for the transcriptomic profiling. Briefly, the cells were incubated with dissociation buffer (50% Accutase, 50% DPBS, and ≥50 units papain (Worthington; Cat.No. LK003178)) at 37 °C for 20 min. Then, wash buffer (DMEM/F12 supplemented with GlutaMax, 10 nM ROCK inhibitor, and ≥ 500 Kunitz units DNaseI (Worthington, Cat.No. LK003172)) was added to the cells. The cells were collected, washed 3 times with 0.1% BSA in DPBS and centrifuged at 150 × g for 10 min to remove the dead cell debris.

**THP-1 differentiation**

$2 \times 10^6$ cells of THP-1_1-E9 were seeded in 10 cm dishes and treated with 30 ng/mL of PMA (Sigma) for 24 h and 96 h. As a negative control, THP-1_1-E9 cells were treated with 1/1,000 volume DMSO (Wako) in the culturing medium for 24 h and 96 h.

**RNA purification**

The harvested cells were subjected to RNA extraction and purification using the RNeasy Kit (Qiagen) with on-column DNaseI treatment according to the manufacturer's instructions. The eluted RNA was quantified by the NanoDrop™ spectrophotometry platform (Thermo Fisher Scientific). The purified RNA samples were stored at -80 °C until they were used for the CFC-seq, ssCAGE and RNA-seq libraries construction.

**Cell fractionation**

Subcellular fractionation of $i^3N$ cells was performed similarly as previously described[1] with minor optimization. Briefly, approximately 10 million cells were used per fractionation experiment to obtain cytoplasmic (CYTO), soluble nucleoplasmic (SNUC) and chromatin (CHR) fractions. Cells detached by Accutase were washed with cold DPBS and lysed for 5 min using a cold lysis buffer containing 0.15% Igepal CA-630, 10 mM Tris pH 7.5, 150 mM NaCl. The lysate was overlaid on a sucrose cushion containing 10 mM Tris pH 7.5, 150mM NaCl, 24% sucrose and centrifuged at 1000 × g for 10 min at 4 °C. The supernatant was removed and centrifuged at 14,000 × g for 1 min, after which the supernatant was taken as the CYTO fraction. The nuclear pellet was washed once in PBS containing 0.5 mM EDTA pH 8.0 and resuspended in a buffer containing 20 mM Tris pH 7.5, 50% glycerol, 75 mM NaCl, 0.5 mM EDTA. An equal volume of nuclear lysis buffer containing 10 mM Tris pH 7.5, 300 mM NaCl, 1M Urea, 1% Igepal CA-630, 7.5 mM $MgCl_2$, 0.2 mM EDTA was added and incubated on ice for 5 min to lyse the nuclei. After centrifugation at 13,000 x g for 2 min at 4 °C, the supernatant was taken as the SNUC fraction and the pellet as the CHR fraction. The chromatin pellet was washed once in PBS containing 0.5 mM EDTA pH 8.0 and resuspended in the same buffer. RNA from each fraction was isolated using Trizol LS (Invitrogen), according to the manufacturer's instructions. Prior to RNA extraction, chromatin which formed a tight precipitate was solubilized in Trizol LS by first passing through a 27-gauge needle, then treating with a dounce homogenizer. Extracted RNA was subjected to DNase I treatment followed by phenol-chloroform extraction. Average RNA yields were 260 µg, 53 µg and 7 µg from the cytoplasmic, nucleoplasmic and chromatin fractions, respectively. The RNA from the chromatin fraction was subjected to CFC-seq cDNA library construction and sequenced by PromethION.

**CFC-seq experimental workflow**

For the total RNAs from the iPS, NSC and cortical neuron, 10 µg of RNAs were polyadenylated by incubating with E-coli poly(A) Polymerase (PAP) (NEB M0276) at 37°C for 15 min and purified with AMPure RNA Clean XP beads. For the total RNAs from the THP-1 series (DMSO 24h, DMSO 96h, PMA 24h and PMA 96h), 80 - 100 µg of RNAs were used both for PAP-treated and PAP-nontreated RNAs. For the iPS chromatin fraction RNAs, 5 µg of RNAs were polyadenylated. The RNA was reverse transcribed by Prime Script II Reverse Transcriptase (Takara Bio) at 42°C for 60 min with different poly-dT primer sets dependent on samples. For the PAP treated RNAs from iPS, NSC and cortical neuron, reverse transcription was done by poly-dT primer containing UMI: oligodT_16VN_UMI15 (GAGATGTCTCGTGGGCTCGG$N_{15}$CTACG$T_{16}$VN). The PAP-treated and PAP-nontreated RNAs from THP-1 series and PAP-treated chromatin RNAs were reverse transcribed with poly-dT primer containing ONT barcode: FL-ONT_dT16VN_NB01-NB08 (GAGATGTCTCGTGGGCTCGG[NB01-NB08]CTACG$T_{16}$VN). The product was purified with RNAClean XP beads. Cap-trapping from the RNA/cDNA hybrids was performed as previously described.[2] Then, RNA was digested with RNase H (Takara Bio) at 37°C for 30 min and the cDNA was purified with AMPureXP beads. Double stranded 5' linker of N6 and GN5 (**Table S17**) were mixed at a ratio of 1:4 and ligated to the cDNA with Mighty Mix (Takara Bio) for overnight and the ligated cDNA was purified with AMPure XP beads. Shrimp Alkaline Phosphatase (Takara Bio) was used to remove phosphates at the ligated linker and purified with AMPureXP beads. The 5' linker ligated cDNA was then subjected to second strand synthesis with KAPA HiFi mix (Roche) using a second strand primer with an UMI of 25 nucleotides (CTACACTCGTCGGCAGCGTC$N_{25}$GTGGTATCAACGCAGAGTAC) at 95°C for 5 min, 55°C for 5 min and 72°C for 30 min. Exonuclease I (Takara Bio) was added to digest the excessed primers at 37°C for 30 min. After AMPureXP purification, the cDNA/DNA hybrids derived from the neuron series were amplified with PrimerSTAR GXL DNA polymerase (Takara Bio) for 10 cycles. The cDNA/DNA hybrids from the iPS chromatin-bound fraction were amplified with LongAmpR Taq DNA Polymerase (NEB, M0323) for 7 cycles. The amplified products were purified by AMPureXP. The amplified products from the Neuron series and the cDNA/DNA hybrids from the THP-1 series were subjected to library construction using the kit SQK-LSK110 (Oxford Nanopore Technologies) with the manufacturer's instructions. The libraries were sequenced with R9.4.1 flow cell (FLO-MIN106) in MinION sequencer, by adding 50 fmol DNA per flow cell and sequenced for 72 hrs. For the chromatin-bound iPSC, purified product was subjected to library construction using kit V14 SQK-LSK114 (Oxford Nanopore Technologies). The libraries were sequenced in the R10.4.1 flow cell in the PromethION sequencer. For each sample replicate, the technical runs were pooled for further downstream analyses. All the oligo sequences are listed in **Table S17**.

**RNA-seq**

The purified RNA from the 3 cell types with duplicates was subjected to library construction with the TruSeq Stranded Total RNA Sample Prep Kit with Ribo-Zero Human/Mouse/Rat (Illumina, 20020596) & TruSeq RNA Single Indexes Set A (Illumina, 20020492). After quantification, the 6 libraries were sequenced in one lane of S4 flow cell in the NovaSeq 6000 Sequencing System (Illumina).

**Single-strand CAGE (ssCAGE)**

Following RNA extraction and purification from iPSC, NSc and Neuron, 15 μg of input RNA per sample was processed for single-strand CAGE library construction.[3] Sequencing was performed using the Illumina NextSeq 1000/2000 P2 Reagents (200 cycles, v3) with a 100-bp paired-end configuration.

### Section 2 | Pre-processing and quality control

#### Pre-processing of the long-read ONT data

Basecalling was performed using Dorado (v0.2.4, Oxford Nanopore Technologies) to generate FASTQ files from raw FAST5 data files using the high-accuracy model "dna_r9.4.1_e8_sup@v3.3". Reads were filtered for quality, retaining only those with a mean Q-score > 10 for the downstream analyses. To ensure high-quality transcript models, adapter sequences were trimmed and strands were oriented using primer chop (https://gitlab.com/mcfrith/primer-chop). Only reads containing confirmed adaptors at both ends were retained, while the relative position of the head and tail linkers was used to assign read orientation. Subsequent to adaptor removal, poly(A) tails including those basecalled as mixed adenine and guanine dinucleotides were identified, recorded and trimmed. We developed tail trimmer (v1.4), which utilizes a 20 bp sliding window to identify and trim poly(A) sequences, allowing a maximum of 5 terminal adenines to remain.

```
# Basecalling for Neuron & THP1 series:
dorado basecaller -x cuda:all --min-qscore 10 --emit-fastq dna_r9.4.1_e8_sup@v3.3 $POD5 | gzip >
$POD5/$POD5.fastq.gz
# Basecalling for iPS chromatin bound:
dorado basecaller -x cuda:all --min-qscore 10 --emit-fastq dna_r10.4.1_e8.2_400bps_sup@v4.1.0
$POD5 | gzip > $POD5/$POD5.fastq.gz

# Pimer chop
primer-chop \
-q \
-P 12 \
riken-yonsei-primers_minus_poly-A.fa \
"dorado_basecaller/${LINE}.fastq.gz" \
dorado_primer-chop_minus_poly-A

# Re-orentiation
sed -i '1~4 s/$/ strand=-/g' dorado_primer-chop_minus_poly-A/good-rev.fq &
sed -i '1~4 s/$/ strand=+/g' dorado_primer-chop_minus_poly-A/good-fwd.fq
cat dorado_primer-chop_minus_poly-A/good-fwd.fq dorado_primer-chop_minus_poly-A/good-rev.fq >
dorado_primer-chop_minus_poly-A/good.fq

# tail trimmer
tail_trimmer_v1.4.pl \
--rescue_tail_nt=5 \
--revise_read_prefix="$LINE" \
--fastq_path="$LINE/dorado_primer-chop_minus_poly-A/good.fq.gz" \
```

```
--out_dir="$LINE/dorado_primer-chop_minus_poly-A_tail-trimmer"
```

### Mapping to genome and TranscriptClean

The trimmed and orientated reads were then mapped to the human reference genome (GRCh38, GCA_000001405.15), using minimap2 (v2.17-r974-dirty),[4] guided by GENCODE (v39) annotations. Raw alignments were error-corrected using TranscriptClean (v2.0.3).[5] Splice junction (SJ) correction was guided by a reference set comprising GENCODE v39 junctions and high confidence SJs identified short-read RNA seq of the Neuron series. At the locus-level, TranscriptClean removed 40,024 SJs and introduced 8,107 SJs while 1,654,272 (97.17%) remained unchanged. As a requirement for running TALON, an additional tag was added to the corrected alignments using the TALON (v5)[6] talon_label_reads function which calculates the fraction of adenines in a 16-bp window within the genome immediately downstream of the last 3'-bp alignment of the read. However, internal priming was defined universally across different assemblers as described in the next section.

```
[1] minimap2 -I 1000G -k 15 -d reference.fa.mmi reference.fa
[2] minimap2 -t [threads] -2 -ax splice -uf --MD --junc-bed gencode.v39.annotation.bed --
secondary=no GRCh38_no_alt_analysis_set_GCA_000001405.15.fasta.mmi [fastq_file]> [sam_file]
[3] python TranscriptClean.py -t [threads] --sam [sam_file] --genome
GRCh38_no_alt_analysis_set_GCA_000001405.15.fasta -j gencode.v39.annotation.SJs.txt --
outprefix sample --deleteTmp --tmpDir sample/tmp
[4] talon_label_reads --f pass_trim_clean_corrected.sam --ar 16 --fracA 0.5 --g
GRCh38_no_alt_analysis_set_GCA_000001405.15.fasta --t 12 --tmpDir sample/tmp --deleteTmp --o
sample
```

### Comparative analysis of long-read 5' end precision

To benchmark the 5' end accuracy of CFC-seq against established methodologies, raw FASTQ datasets from five distinct long-read sequencing protocols targeting the same iPSC line (WTC11) were obtained from ENCODE LRGASP consortium.[7] These included CAP_trap_PacBio (ENCSR309IKK), uncap_deplete_PacBio (ENCSR507JOF), TSO_ONT (ENCSR539ZXJ), R2C2_ONT (ENCSR925UQZ) and dRNA_ONT (ENCSR392BGY). All reads were aligned to hg38 by minimap2 as described above. The alignments of these libraries and our iPSC libraries were intersected with GENCODE v39 protein-coding exons using bedtools intersect. Reads overlapping with $\geq$ 90% of an annotated protein-coding exon were classified as mRNA and retained for downstream analysis, excluding mitochondrial transcripts.

To quantify transcriptional initiation precision, the empirical 5' end coordinates of these alignments were then intersected with three independent genomic reference datasets defining promoter and active regulatory boundaries: 1) candidate *cis*-regulatory elements (cCREs) from the SCREEN registry annotated as promoter-like or enhancer-like signatures, 2) open chromatin peaks derived from single-nucleus ATAC-seq data generated from the identical WTC11 iPSC line, and 3) robust CAGE-defined promoter clusters obtained from the FANTOM5 consortium (median width 300bp). Comprehensive genomic coordinates for these reference annotations are archived in **Supplementary table S18**.

**Assessment of 3' ends capture and filtration of potential internal priming**

To evaluate the efficacy of in vitro poly(A)-tailing (PAT) for non-poly(A) transcript capture, we performed comparative library testing using THP-1 cells processed with or without the PAT enzyme cascade. Natural non-poly(A) control genes, including small nucleolar RNAs (snoRNAs) and replication-dependent histone gene blocks , were utilized to evaluate non-poly(A) RNA target rescue efficiency.

The single nucleotide 3' end coordinates were extracted from all alignments, and the flanking genomic sequences (± 50 nt) were retrieved from the human reference genome GRCh38 (GCA_000001405.15). To mitigate internal priming artifacts caused by oligo-dT hybridization to homopolymeric adenine stretches within transcript bodies, these flanking regions were profiled for downstream adenine composition. A transcript was flagged if adenine composition exceeded 50% within 16 nucleotides immediately downstream of the 3' end, or exceeding 75% within the 8 nucleotides immediately downstream. Termini directly corresponding to annotated GENCODE v39 transcript models were exempt from this filtering cascade.

**Computational identification and hierarchical scoring of PAS motifs**

For polyadenylation signal (PAS) motif identification, we performed a genome-wide search using scanMotifGenomeWide.pl from Homer (v4.11)[8] with a position weight matrix derived from GENCODE v39 annotation. Only the loci with score > 3 were used for the analysis. The genomic locations of these PAS motifs were overlapped with the single-nucleotide locations 25 nt upstream of the 3' ends by running bedtools closest. Presence of PAS motif with position of its last nucleotide located -5 nt to -35 nt from the 3' ends was annotated as PAS-positive. We applied a hierarchical search strategy based on motif strength, starting from PAS motifs with score >6 ("AATAAA" "ATTAAA"), followed by PAS motifs with score > 4 ("TATAAA", "AGTAAA", "AATATA"), and by motifs with score >3 ("CATAAA", "GATAAA", "AAAAAA", "TTTAAA", "ACTAAA", "AATACA", "AATAGA", "AAGAAA", "AATAAG", "AATAAT", "AATGAA", "AATTAA", "ATTATA").

```
[1] scanMotifGenomeWide.pl PAS.motif hg38 -bed -keepAll -p 10 > output.bed
[2] bedtools closest -a all3n_up25.sort.bed.gz -b
GENCODE_polyA_signal.motif_rng.main.FASTA.6.bed.gz -s -D a | cut -f-6,10- | gzip >
all3n_up25.PAS6.bed.gz
[3] bedtools closest -a all3n_up25.sort.bed.gz -b
GENCODE_polyA_signal.motif_rng.main.FASTA.4.bed.gz -s -D a | cut -f-6,10- | gzip >
all3n_up25.PAS6.bed.gz
[4] bedtools closest -a all3n_up25.sort.bed.gz -b
GENCODE_polyA_signal.motif_rng.main.FASTA.3.bed.gz -s -D a | cut -f-6,10- | gzip >
all3n_up25.PAS6.bed.gz
```

#### Capture of non-poly(A) ncRNA and incomplete transcripts derived from PAT protocol

Following the exclusion of known non-poly(A) reference transcripts and loci affected by internal priming, the baseline transcript capture layout of non-PAT standard poly-dT protocol was compared with PAT protocol (**Ext_Fig. 1g**). To investigate whether the PAT protocol increases the prevalence of incomplete transcript models, protein-coding mRNAs lacking a PAS motif were classified based on genomic coordinations of their 3' ends. Finally, PAT-specific 3'ends were isolated by subtracting the 3' end coordinates detected in the non-PAT control. These 3' ends were intersected with different genomic regions derived from annotations of GENCODE protein-coding transcripts.

#### Poly(A)-tail prediction

In order to call bona fide 3' ends for the transcripts detected by CFC-seq, we trained a random forest classifier on a curated database of 3' ends identified by FLAM-seq and 3p-seq.[9,10] Briefly, we collected all 3' ends detected by CFC-seq and flagged those overlapping with the curated (hereon "reference") dataset within a window of 10 nt. We then trained a random forest classifier using the ranger package (v. 0.16.0) in R 4.3.1, considering the 3' ends found in the reference as true positives and defining a set of features to train the model on, including the presence, type and position of polyadenylation signals and the nucleotide composition of a 50 nt window upstream and downstream of each 3' end. We then used the trained model to assign a probability score ("poly(A) score") to each 3' end in the CFC-seq dataset. We defined an upper threshold at which 95% of 3' ends found in the reference and containing a polyadenylation signal were classified as true to define "poly(A)" 3' ends (threshold > 0.34), and a lower threshold at which 95% of all 3' ends were classified as true to flag "non-poly(A)" 3' ends (threshold < 7.6e-5). For downstream analyses, PAS-positive 3' ends and 3' ends classified as true poly(A) were considered as poly(A) 3' ends while the others were considered as non-poly(A). The read-based analysis result was transferred to the transcript-level and maintained in **Table S4**.

#### Confident splice junctions from short-read RNA-seq

To obtain a confident splice junction file (SJ.out.tab), all the fastq files of the short-read RNA-seq were combined into read1 and read2 fastq files and subjected to STAR for genome mapping using hg38 GENCODE v39 as reference.

```
[1] zcat *_R1_*.fastq.gz | gzip > all_reads_R1.fastq.gz
[2] zcat *_R2_*.fastq.gz | gzip > all_reads_R2.fastq.gz
[3] STAR \
  --runThreadN 8 \
  --genomeDir /path/to/STAR_index_hg38_GENCODEv39 \
  --readFilesIn all_reads_R1.fastq.gz all_reads_R2.fastq.gz \
  --readFilesCommand zcat \
  --outFileNamePrefix star_output/ \
  --outSAMtype BAM Unsorted \
  --outSJfilterReads Unique \
  --outFilterMultimapNmax 1 \
  --alignSJoverhangMin 8 \
  --alignSJDBoverhangMin 1 \
  --outSAMstrandField intronMotif
```

#### TSS clusters and tCREs identification by SCAFE

The BAM files containing all alignments mapped to the hg38 genome were used as the input for SCAFE v1.01.[11] The functions acquired from SCAFE were incorporated into SALA, where detailed workflow and explanation can be found in the GitHub (https://github.com/fantom-prj/SALA). Briefly, scafe.workflow.bk.bam_to_ctss extracted alignments with ≤ 3 nt softclip at the 5' end and ≥ 30 matched nt, where ~70% reads passed. Approximately 30% of reads from the CFC-seq had a relatively long 5' end unaligned region (> 3 nt), these reads were excluded from TSS cluster identification while their reads were used for quantification. The qualified alignments (~165 million) were converted into CTSS, which were grouped into TSS clusters. According to the multiple properties of each TSS cluster (percentage of CTSS with unencoded G, corrected expression, read count, summit count and flanking count), a multiple logistic regression model was trained to distinguish TSS clusters that are likely genuine. The identification of these TSS clusters was performed per replicate where cutoffs of at least 3 reads and at least 1 read with unencoded G were applied. Next, scafe.tool.cm.aggregate defines tCREs by extending and merging overlapped TSS clusters defined from the replicates. This aggregation step only merges the clusters without pooling signals to generate new TSS clusters. The TSS clusters were then filtered by scafe.tool.cm.filter according to the multiple logistic regression model. Additional commands including scafe.tool.cm.directionality and scafe.tool.cm.annotate were used to calculate bi-directional reads in each tCRE and pre-annotate the tCREs with nearby GENCODE transcript model. The codes for running SCAFE for CFC-seq in this study are listed:

```
#Convert bam into CTSS with and without unencoded G:
[1] scafe.tool.bk.bam_to_ctss
--TSS_mode=softclip \
--bamPath=[bamPath] \
--unencoded_G_upstrm_nt=3 \
--max_thread=5 \
--genome=hg38.gencode_v39 \
--max_softclip_length=3 \
--outputPrefix=[outputPrefix] \
--outDir=[outDir]

#Aggregate individual library into cell types [2-4]:
[2] scafe.tool.cm.aggregate \
--lib_list_path=[path to CTSS per library ] \
--max_thread=5 \
--genome=hg38.gencode_v39 \
--outputPrefix=[outputPrefix] \
--outDir=$baseDir/out/aggregate

[3] scafe.tool.cm.ctss_to_bigwig \
--genome=hg38.gencode_v39 \
--ctss_bed_path=aggregate.collapse.ctss.bed.gz \
--outputPrefix=[outputPrefix.all] \
--outDir=$baseDir/out/ctss_to_bigwig

[4] scafe.tool.cm.ctss_to_bigwig \
--genome=hg38.gencode_v39 \
--ctss_bed_path=aggregate.unencoded_G.collapse.ctss.bed.gz \
--outputPrefix=[outputPrefix.ung] \
--outDir=$baseDir/out/ctss_to_bigwig

#Identify TSS clusters, tCREs and annotate the tCREs [5-9]:
[5] scafe.tool.cm.aggregate \
--lib_list_path=[path to CTSS per library ] \
```

```
--max_thread=5 \
--genome=hg38.gencode_v39 \
--outputPrefix=[outputPrefix] \
--outDir=$baseDir/out/aggregate

[6] scafe.tool.cm.cluster \
--overwrite=yes \
--cluster_ctss_bed_path=aggregate.collapse.ctss.bed.gz \
--count_ctss_bed_path=aggregate.unencoded_G.collapse.ctss.bed.gz \
--min_summit_count=0 \
--min_cluster_count=1 \
--outputPrefix=[outputPrefix] \
--outDir=$baseDir/out/cluster

[7] scafe.tool.cm.filter \
--overwrite=yes \
--ctss_bed_path=aggregate.collapse.ctss.bed.gz \
--ung_ctss_bed_path=aggregate.unencoded_G.collapse.ctss.bed.gz \
--tssCluster_bed_path=tssCluster.bed.gz \
--genome=hg38.gencode_v39 \
--outputPrefix=[outputPrefix] \
--outDir=$baseDir/out/filter

[8] scafe.tool.cm.annotate \
--overwrite=yes \
--tssCluster_bed_path=tssCluster.default.filtered.bed.gz \
--tssCluster_info_path=tssCluster.log.tsv \
--min_CRE_count=3 \
--genome=hg38.gencode_v39 \
--outputPrefix=[outputPrefix] \
--outDir=$baseDir/out/annotate

[9] scafe.tool.cm.directionality \
--overwrite=yes \
--CRE_bed_path=CRE.coord.bed.gz \
--CRE_info_path=CRE.info.tsv.gz \
--ctss_bed_path=aggregate.collapse.ctss.bed.gz \
--outputPrefix=[outputPrefix] \
--outDir=$baseDir/out/directionality
```

### Promoter typing

The tCREs were then classified for the promoter types by intersecting with SCREEN cCREs, in an order of promoter-like, enhancer-like and CTCF-alone. If a tCRE was intersected with both promoter-like and enhancer-like SCREEN cCREs, the tCRE was defined as promoter-like. If no cCRE was found to intersect with the tCREs, these tCREs were defined as "unclassed". These promoter types were directly transferred to the TSS clusters and transcript models. At the gene model level, promoter types of the linked transcript models were collected. If more than one promoter type were found in one gene model, the promoter type was decided at the priority of promoter-like > enhancer-like > CTCF-alone > unclassed. Notably, the transcript models transcribed from unclassed tCREs without ATAC support were included in the finalized transcriptome, annotated with the linked tCRE.

Expression level of tCREs was measured by CTSS of qualified alignments located inside the genuine TSS clusters, where the per-cluster signal was collapsed into per-tCRE level. The signal was normalized across all the samples with individual replicates by using the RLE method in edgeR. In the preparation of running SALA, strand-specific TSS clusters were merged if they are within 75 nt distance. If the signal from the merged clusters revealed bi-modal distribution (>20% from each cluster), the merged clusters were split at the midpoint of the two summits. This extension was designed for concluding reads and reference transcript models into the same read class if their 5' ends are in a proximity.

### Comparison of CFC-seq and CAGE in identification of TSSs and tCREs

Sequenced reads from ssCAGE were de-multiplexed and aligned to the hg38 human genome assembly using the STAR aligner.[12] This yielded a median depth of 60 million mapped paired-end reads across the generated libraries. Additionally, single-end alignment was also performed using the Read1 alone. Along the analysis with different MAPQ thresholds, alignments were filtered by MAPQ to obtain CTSS in the SCAFE pipeline. To assess the content of these tCREs *de novo* identified from CFC-seq, ssCAGE from the corresponding samples were analysed. The reads from ssCAGE were independently subjected to the SCAFE pipeline or aggregated with the CFC-seq CTSS by SCAFE (**Ext_Fig. 2a**). The expression level shown in **Ext_Fig. 3j** was derived from the aggregated tCREs derived from both CFC-seq and ssCAGE signals. Read counts per tCRE were included only if the CTSSs were positioned inside genuine TSS clusters. The counts were RLE normalized using the edgeR package.

### Mappability of repetitive elements using CFC-seq and CAGE

To determine whether the extended read length from CFC-seq improves the resolution of TSSs within or near repetitive elements, we compared the MAPQ across technologies. We utilized a set of 202,862 TSS clusters identified via joint SCAFE analysis that were detected across all three platforms: CFC-seq, single-end CAGE, and paired-end CAGE. Summits of these TSS clusters were extended 150 nt downstream and intersected with the genomic coordination of RepeatMasker obtained from UCSC. Each TSS cluster was assigned the maximum MAPQ score of its constituent reads per platform. As the MAPQ is not directly comparable between long-read and short-read aligners, the MAPQ score from short-read datasets were revised for visualization: 0 to 0; 2 to 5; 3 to 10; and 225 to 20.

To compare the evolutionary age of the repetitive elements associated to CFC-seq specific TSS clusters, we summarized the milliDiv (divergence from the consensus sequence) as provided by RepeatMasker. From the 32,365 TSS clusters overlapped with repetitive elements, we extracted the repetitive elements successfully resolved by CFC-seq but not CAGE at the TSS clusters (MAPQ ≥ 20 from CFC-seq and MAPQ < 225 from single-end CAGE). This allowed us to determine if CFC-seq's increased read length is particularly critical for resolving transcription from younger, more sequence-homologous transposable elements.

Genomic coordination of human short tandem repeats (STR) were obtained from the HipSTR[13] reference dataset (hg38.hipstr_reference.bed.gz). These STR intervals were intersected with the summits of major-strand CRE. STR groups with fewer than 30 independent intersections across our tCREs dataset were excluded from downstream analysis. To identify motif overrepresentation, two tailed Fisher's exact tests were performed for each distinct promoter class (promoter-like, enhancer-like, CTCF-alone and unclassified tCREs), using all remaining classes as the background control set.

### Section 3 | Transcriptome construction

#### Transcript model construction using Transcript Start-site Aware Long-read Assembler (SALA)

SALA reconstructs transcriptomes through four sequential modules: feature collection, transcript model assembly, transcript model filtering and gene annotation. The steps were documented in the wiki of SALA GitHub. The parameters used for SALA Final are the same as the "sensitive mode", while SALA Default is from the "default mode" in GitHub. Notably, these two runs are independent and generate transcript and gene models independently. SALA Final is the primary dataset for downstream analyses and was used to build the SACAGE transcriptome of FANTOM6 Interactome[14] by adding all GENCODE (v39) transcript models and FANTOM CAT permissive transcript models. Therefore, transcript and gene IDs from SALA Final are transferrable with SACAGE. Detailed description of SALA can be found in the GitHub wiki page (https://github.com/fantom-prj/SALA/wiki).

##### Feature collection

SALA integrates multiple evidence to define high-confidence transcript boundaries and internal structures. 5' end clusters and summits identified via SCAFE were utilized as primary transcription start sites (TSS). For 3' ends, tags were collapsed into clusters using paraclu (minimum 3 tags per cluster) to define termination signal clusters and summits. Orthogonal evidence was incorporated by including SJs from sample-matched short-read RNA-seq (SJ.out.tab from STAR) and reference annotations (GENCODE v39). Splice junctions (SJs) were extracted from alignments before TranscriptClean, alongside maximum base-calling quality scores for the flanking sequences (± 3nt). For SJs without external support, we applied a threshold of a summarized base-calling score ≥ 10 and a read count ≥ 3, for technical confidence (**Ext_Fig. 7b**). To ensure robust boundary definition, SCAFE-defined TSS clusters were merged within a 75-nt window and resulted as extended 5' end clusters (ex5_clusters). This effectively included GENCODE transcript model 5' ends and linked annotated transcript models to the reads in the same ex5_clusters. Bimodal clusters (exhibiting > 10% signal across the distribution) were split at their midpoint to distinguish proximal ex5_clusters. 3' end clusters were merged within a 150-nt window. Clusters supported by either SCAFE, paraclu cluster, or GENCODE 5' and 3' ends annotation were designated as confident features. In feature collection, SALA Final used the same parameter as SALA Default.

```
# Convert read bam file into bed file
```

```
[1] sh ./SALA/code/others/SALA.input.bamtobed.sh \
[bamTC_path.txt (path of bam files after transcriptclean)] \
[./SALA/input/bam_to_bed (output_directory)] \
./SALA/resources

# Prepare 3' end cluster
[2] perl ./SALA/code/SALA/3n_cluster/transcript_bed_to_end_bed_bigwig.pl \
./SALA/input/bam_to_bed/combined.bed.bgz \
./SALA/resources/chrom.sizes.tsv \
[outputPrefix] \
[./SALA/input/CTES_clusters/end3_bed_bigwig (output_directory)] \
./SALA/resources/bin/bedGraphToBigWig/bedGraphToBigWig

[3] perl ./SALA/code/SCAFEv1.0.1/scripts/scafe.tool.cm.cluster \
--overwrite=yes \
--cluster_ctss_bed_path=./SALA/input/CTES_clusters/end3_bed_bigwig/outputPrefix.end3.bed \
--count_ctss_bed_path=./SALA/input/CTES_clusters/end3_bed_bigwig/outputPrefix.end3.bed \
--min_summit_count=3 \
--min_nt_count=3 \
--min_cluster_count=5 \
--outputPrefix=[outputPrefix.CTES.s3_n3_c5] \
--outDir=./SALA/input/CTES_clusters/scafe/cluster

# Prepare splice junction
[4] perl ./SALA /code/SALA/junction_extractor/junction_extractor.pl \
--in_bam=[iPSC_rep1_run1_subset.sorted.bam(bam files before transcriptclean)] \
--chrom_size_path=./resources/chrom.sizes.tsv \
--chrom_fasta_path=./resources/GRCh38_no_alt_analysis_set_GCA_000001405.15.fasta \
--out_prefix=[iPSC_rep1_run1] \
--out_dir=./SALA/input/junction_extractor/output \
--max_thread=1 \
--min_nt_qual=10 \
--min_MAPQ=20 \
--samtools_bin=./resources/bin/samtools/samtools \
--bedtools_bin=./resources/bin/bedtools/bedtools \
--tabix_bin=./resources/bin/tabix/tabix \
--bgzip_bin=./resources/bin/bgzip/bgzip

[5] perl ./SALA/code/SALA/junction_extractor/junction_pool.pl \
--outDir=./SALA/input/junction_extractor/pool \
--outTag=[outputPrefix] \
--findStr=./SALA/input/junction_extractor/output/*/log/*.junct.info.tsv.gz
```

### Transcript model assembly and read assignment

Each long-read was assigned a specific 5' end cluster, a 3' end cluster, and a set of SJ IDs. Using the default parameters, confident ex5_clusters were derived from SCAFE requiring at least 3 reads and at least one un-encoded G while confident ex3_clusters require ≥ 5 counts and ≥ 3 summit counts. If a GENCODE annotated TSS and TES is located inside the ex5_clusters and ex3_clusters, these clusters become confident independent of read count. Reads possessing both confident 5' and 3' boundaries, as well as all reference-matching models, were classified as complete transcript models. Reads sharing an identical triplet of features (TSS cluster, 3' cluster, and SJ chain) were collapsed into a unique isoform. To prevent the erroneous identification of nascent or fragmented RNAs as novel isoforms, SALA utilized a hierarchical assignment strategy. Incomplete long-reads (lacking one or both confident boundaries) were first mapped to existing complete models. If an incomplete read's structure was a subset of a complete model, it was assigned as partial support. Only remaining incomplete reads without a matching full-length scaffold were permitted to form independent models. The final coordinates for a model's 5' and 3' ends were defined by the most frequent terminal positions among its constituent reads. In this step, the parameter `--trnscpt_set_end_priority=commonest:summit:longest` was used for SALA Final while for SALA Default, `--trnscpt_set_end_priority=summit:commonest:longest` was used. This setup in SALA Final enabled the observation of individual transcript 5' ends when multiple transcript models derived from the same ex5_clusters. Notably, transcript model assignment is not affected by this parameter. The parameters used for SALA Final is shown as below:

```
[1] end5_guided_assembler_v0.1.20231102.pl
--qry_bed_bgz=[Neuron_THP1.bed.bgz]
--ref_bed_bgz=[GENCODEv39.transcript.bed.bgz]
--chrom_size_path=[hg38.gencode_v39_chrom.sizes.tsv]
--out_dir=[output]
--max_thread=3
--out_prefix=Neuron_THP1
--min_transcript_length=15
--doubtful_end_avoid_summit=yes
--min_exon_length=1
--print_trnscrptID=no
--chrom_fasta_path=[hg38.gencode_v39_genome.fa]
--min_output_qry_count=1
--trnscpt_set_end_priority=commonest:summit:longest #for SALA Final
--doubtful_end_merge_dist=150
--novel_model_prefix=ONTT
--conf_end3_merge_flank=150
```

```
--conf_end5_merge_flank=75
--conf_end5_bed_bgz=[ontCAGE.Neuron_THP1.end5.cluster.bed.bgz]
--conf_end3_bed_bgz=[ontCAGE.Neuron_THP1.end3.cluster.bed.bgz]
--min_summit_dist_split=50
--retain_no_qry_ref_bound_set=no
--doubtful_end_avoid_summit=yes
--min_size_split=100
--min_frac_split=0.2
--signal_end5_bed_bgz=[ontCAGE.Neuron_THP1.end5.signal.bed.bgz]
--signal_end3_bed_bgz=[ontCAGE.Neuron_THP1.end3.signal.bed.bgz]
--conf_end3_add_ref=yes
--conf_end5_add_ref=yes
--conf_junction_bed=[Gencode_v39.junct.bed],
[long_read.hi_qual.junct.bed] ,[short_read.hi_qual.junct.bed]
--min_qry_score=0
```

### Initial gene annotation

The transcript model assembly generates the SALA Raw transcript models, which include all the reference transcripts (even they are not detectable) and novel transcripts. These models were then subjected to initial gene annotation using the parameter `--disable_ref_chain_bound_gene_anno=yes`. This constraint is critical to prevent the erroneous fusion of distinct reference gene models based on the detection of low-level, nascent read-through transcripts. Initial gene annotation is only used for setting thresholds on transcript read count ratio per gene. Another round of gene annotation was performed on the filtered transcript models.

### Filtering for confident transcript models

We include different levels of filtering for benchmarking. Filtering criteria and the resulting number of transcript models are shown in **Extended Figure 4a,d**. In SALA Raw and SALA Final were obtained from a run using SALA sensitive. While the Raw dataset retained all the models excluding internal priming and undetected GENCODE models, the Final dataset was further subjected to different filterings according to the transcript classes. Additionally, SALA Default was also obtained using the "default" mode.

Following assembly, all the transcript models generated by SALA were categorized relative to the GENCODE v39 reference into three classes: known transcripts (assigned with an ENST transcript ID), novel transcripts derived from GENCODE genes (Novel isoforms attached to an ENSG gene ID), and novel transcripts derived from novel transcriptional units (including intergenic, intronic and antisense). To minimize technical artifacts, we first excluded models associated with putative internal priming and removed GENCODE reference transcripts that showed no evidence in our dataset to generate SALA Raw dataset. For SALA Final, only the novel transcript models with their starting sites linked to the confident ex5_cluster supported by SCAFE were kept. Specifically for the novel transcripts derived from GENCODE genes, 5 supporting reads from each of the replicates and 10% transcript ratio according to complete read count from one of the cell types were required. A confident ex3_cluster supported by clustering or GENCODE was also required for this transcript group. For the transcripts from novel transcriptional units (majority are novel lncRNAs), no further filter was applied. Finally, an additional filter was applied to exclude reference transcript models without SCAFE TSS supported for consistent downstream analyses, while this filter is not part of the SALA sensitive mode.

In SALA default, an additional filter was applied to the novel transcripts derived from novel transcriptional units, where at least one complete read from each replicate was required. The requirement of SCAFE TSS support from reference transcript models was voided. For each transcriptome set, transcript models were subjected to the gene annotator with the parameter "--disable_ref_chain_bound_gene_anno=no".

```
[1] perl assemble_gene_annotator_v0.1.pl \
--chrom_size_path=[hg38.gencode_v39_chrom.sizes.tsv] \
--model_bed_bgz=[table4wENST.bed12.bed.bgz] \
--model_info_gz=[table4wENST.info.gz] \
--revert_ref_model_bed_bgz=[GENCODEv39.transcript.bed.bgz] \
--ref_model_gene_link=[GENCODEv39.transcript_to_gene.tsv] \
--novel_gene_prefix=ONTG \
--disable_ref_chain_bound_gene_anno=no \
--min_ref_exon_overlap_pct=10 \
--exon_overlap_dist=-1 \
--locus_merge_dist=100000 \
--exclude_t_type=retained_intron \
--out_prefix=Neuron_THP1_T4_10percent \
--out_dir=[output] \
--max_thread=1
```

### Coding potential of transcript and gene models

The coding potential of all assembled transcripts was evaluated using the Coding Potential Assessment Tool (CPAT, v3.0.4)[15] with default parameters. For novel transcript models, CPAT score

< 0.364 were defined as non-coding RNA (ncRNA). Transcripts meeting this non-coding criterion with a length of 200 bp were further classified as long non-coding RNAs (lncRNAs). For the GENCODE annotated transcript models, we adopted the lncRNAs annotation as in GENCODE. To assign functional categories at the gene level, we utilized a majority-rule logic based on transcript abundance. A novel transcriptional unit was classified as non-coding if > 50% of its total transcriptional output (determined by complete read counts) was derived from ncRNA transcripts. Within this non-coding pool, genes containing at least one ncRNA transcript ≥ 200 bp were designated as lncRNA transcriptional units, while those exclusively producing shorter transcripts were classified as short_ncRNAs.

For downstream analyses, we applied strict filtering to ensure the purity of the non-coding dataset. For novel loci, only ncRNA transcripts associated with ncRNA genes were retained. For GENCODE loci annotated as lncRNA, we included both known and novel lncRNA isoforms. Thus, novel lncRNAs discovered from GENCODE protein-coding genes were not included for downstream analyses.

**Sub-classification of lncRNA transcripts and genes**

From the CFC-seq with 5 cell-types, "SALA Final" detected 6,559 known ncRNA transcripts and identified 115,188 novel ncRNA transcripts. To characterize their genomic context relative to transcript models annotated as protein-coding and pseudogenes (GENCODE v39), we implemented a hierarchical classification system. Transcripts were assigned to one of five mutually exclusive categories using an ascending order of priority: divergent ncRNAs, sense overlap ncRNAs, sense intronic ncRNAs, other sense ncRNAs, antisense ncRNAs, anisense intronic ncRNAs, other antisense ncRNAs and intergenic ncRNAs.

Divergent ncRNAs: transcripts with their TSSs located within 2 kb of a protein-coding or pseudogene transcript TSS on the opposite strand. Sense intronic ncRNAs: transcripts that intersect with protein coding or pseudogenes, with their TSS and the whole transcript lengths lying inside the intron. For those transcripts with <= 10% and > 10% exon sequence overlap with exons are classed as other sense ncRNAs and sense overlap ncRNAs respectively. Antisense ncRNAs: transcripts with >10% of their exon region overlapping with the protein-coding or pseudogenes exon on the opposite strand. Anisense intronic ncRNAs with their entire transcript region located inside the protein-coding or pseudogenes intron on the opposite strand while the remaining antisense ncRNAs are defined as other antisense ncRNAs. Intergenic ncRNAs: All remaining ncRNA transcripts that did not meet the criteria for the previous categories, residing in regions devoid of annotated gene features.

For ncRNA loci containing multiple transcript isoforms, the same hierarchical priority was applied at the loci level. For example, a gene containing both sense-intronic and antisense isoforms was classified as sense-intronic due to the higher hierarchical priority of that category. This systematic approach ensured that each novel transcriptional unit was uniquely categorized based on its most structurally distinct regulatory relationship with the annotated genome.

**Transcript models construction by TALON**

For comparison with SALA, we analysed the CFC-seq libraries (Neuron series and THP1-series) by TALON.[6] Same as SALA, the bam files after TranscriptClean and TALON label containing only primary alignments were used as input for TALON with the same reference transcriptome (GENCODE v39). We used --allowGenomic to include all the permissive models and filtered afterwards. The run was performed on each chromosome and combined manually. After the processing by TALON, transcript IDs were extracted from the TALON database to include all those novel transcripts classed as 'Genomic'. To keep the comparison with SALA and other assemblers consistent, we remove the potential internal primed transcripts in the same way: the fraction of adenines downstream the 3' end of the alignment in the reference genome. All the undetected GENCODE transcript models were also excluded to make it comparable with the SALA analysis. This dataset was termed "TALON_Raw". In addition, "TALON_Read_filtered" was generated by applying a filter for the transcript models as having 5 supporting reads from both replicates of the sample, as in the default filtering suggested by TALON.

```
[1] talon_initialize_database --f gencode.v39.annotation.gtf --g hg38 --a gencode.v39 --o F6_interactome
[2] talon --f config.csv --db F6_interactome.db --build hg38 --threads [threads} --o F6_interactome_run1 && cp F6_interactome.db /scratch/
[3] talon_fetch_reads --db F6_interactome.db --build hg38 --o talon_read_annot/F6_interactome_all
[4] talon_filter_transcripts --db F6_interactome.db --annot gencode.v39 --minCount 1 --minDatasets 1 --allowGenomic --o F6_interactome_permissive_genomic.csv
[5] talon_create_GTF --db F6_interactome.db --build hg38 --annot gencode.v39 --whitelist F6_interactome_permissive_genomic.csv --o F6_interactome_permissive_genomic_talon.gtf
```

#### Transcript models construction by Isoquant

IsoQuant (v3.4.1)[16] assembling was performed using two distinct modes (Sensitive and Standard) with the following parameters:

```
#IsoQuant Sensitive:
[1] isoquant.py -d nanopore
--bam_list BAM_list.txt
--reference GRCh38_no_alt_analysis_set_GCA_000001405.15.fasta
-p Neuron_Series_THP1
--model_construction_strategy sensitive_ont
--genedb gencode.v39.primary_assembly.annotation.gtf
--threads 40
-o IsoQuant_NS_THP1_sensitive

#IsoQuant Default:
[1] isoquant.py -d nanopore
--bam_list BAM_list.txt
--reference GRCh38_no_alt_analysis_set_GCA_000001405.15.fasta
-p Neuron_Series_THP1
--genedb gencode.v39.primary_assembly.annotation.gtf
--threads 40
-o IsoQuant_NS_THP1_standard
```

#### Construction of different GTF files

The transcriptome datasets (SALA Raw, SALA Final and SALA Default) were used to construct corresponding GTF files. These GTF files were restricted to gene, transcript and exon features. The attribute field for these features was standardized to include gene_id, transcript_id, gene_type, gene_name, transcript_type, transcript_name, gene_novelty, transcript_novelty and exon_number. Reference gene and transcript annotations were inherited from GENCODE v39. When alternative TSSs and TESs were detected from the reference genes or transcripts, their coordinate boundaries were updated in the GTF file, and the gene_novelty and transcript_novelty fields were flagged as "GENCODE_updated". While the GTF file of SALA Raw contains all novel transcripts (without internal priming affected ones) and all reference transcripts, SALA Final and Default contains only full-length detected reference transcript models.

To facilitate annotation and quantification, reference models were integrated back to the detectable transcriptomes. For instance, SALA Final with partially detectable reference transcript models were used to generate GTF files for Bambu quantification and reference of short-read RNA-seq quantification by kallisto. SALA Final with all reference transcript models were used as a reference for iPSC chromatin-bound transcriptome construction in SALA. GTF files derived from other assemblers are also provided.

#### SALA annotation on Chromatin-bound iPSC dataset

From the CFC-seq of iPSC chromatin-bound RNA, sequencing reads were pre-processed as described above and subjected to SALA. This run was applied to SALA sensitive mode using SALA Final and all GENCODE (v39) transcript models as reference. The parameters used and filtering conditions were the same as SALA Final.

### Section 4 | Downstream analyses

#### Analyses on short-read RNA-seq

Short-read RNA-seq FASTQ files of iPSC, NSC and Neuron were subjected to Kallisto,[17] using the final transcriptome with all the completely and partially detected GENCODE models, excluding ribosomal RNA, as the index input. Only the transcript models and gene models detected by the CFC-seq of iPSC, NSC and Neuron with at least one complete read were used for the comparison. Transcript models and gene models with Kallisto quantification >0 from any of the samples were considered as detectable by short-read RNA-seq.

```
[1] kallisto quant -i [kallisto_index] -o [out_dir] --rf-stranded [fq_path1] [fq_path2]
```

#### Quantification of CFC-seq samples

Bambu[18] (https://github.com/GoekeLab/bambu) was modified to account for the increased presence of full-length reads and enhanced detection of transcript start and end sites. Traditionally, Bambu constructs read-classes based on shared splice junctions. Now, we divide these read classes into subgroups by incorporating transcript annotations with refined start and end sites produced by SALA. Briefly, for each first exon of spliced read-classes, the start positions of all exons in the provided annotations that share the 3’ splice site were used as thresholds for grouping reads. A tolerance of 35 bp was applied to each threshold to accommodate small positional variations, and reads were assigned to the largest exon division they matched. Reads longer than the largest annotated first exon were grouped separately. The same process was applied to the last exon, but based on the 5’ splice site. These modifications were implemented in the test_split_read_classes branch. To ensure Bambu recognizes these more granular read classes, we set the opt.discovery = list(min.exonDistance = 0) parameter, reducing the allowance for deviation when assigning read classes to reference transcripts before the Expectation-Maximization (EM) quantification. Additionally, due to the presence of overlapping transcripts from different genes, we preprocessed the SALA gtf file by performing *de novo* gene annotation using the assignGeneIds function from Bambu. This function assigns new gene IDs based on overlapping stranded exon regions, enabling the consideration of overlapping transcripts as a single entity during EM quantification. The quantification was run with the following commands in R:

```
[1] annotations = prepareAnnotations(SALA_table5)
```

```
[2] mcols(annotations)$GENEID = bambu:::assignGeneIds(annotations,
GRangesList())$GENEID
[3] dir.create(paste0(results_dir,"/rcOut"), recursive=TRUE)
[4] se.new <- bambu(reads =  c(Neuron_Series_bam, Monocytes_bam),
      annotations = annotations,
      genome = fa.file,
      discovery = FALSE,
      opt.discovery = list(min.exonDistance = 0),
      rcOutDir = paste0(results_dir,"/rcOut"),returnDistTable=TRUE)
[5] saveRDS(se.new, paste0(results_dir,"/se.rds"))
[6] writeBambuOutput(se.new, results_dir)
```

The transcript CPM and transcript count outputs of the libraries were combined into replicate level. Gene CPM and gene count were obtained by summing the transcript values into gene level.

**Comparing final gene and transcript models with four reference repositories**

To evaluate the novelty and comprehensiveness of our final transcriptome, we utilized the comparison mode of SALA to intersect our models with multiple independent datasets at both the isoform and gene levels. The reference scaffold for this comparison consisted of the SALA Final transcriptome, supplemented by undetected or partially detected models from GENCODE v39. The transcripts from four major annotation resources including Refseq (release from 2024_08), GENCODE (v47), FANTOM CAT (permissive) and LncBook (v2), were subjected to SALA as a query. A transcript model from an external dataset was defined as a match if it shared identical ex5_cluster, ex3_cluster, and splice-junction chains with a CFC-seq model, based on the standard SALA assignment criteria.

For gene level matching, the integrated transcript model output was processed through the SALA gene annotator. We utilized the CFC-seq transcript-to-gene associations as the primary reference to map external transcripts to our identified gene loci. Those gene models annotated with any external transcript models were considered as detection of the gene models. The criteria of matching were the same as the original SALA run: transcript models sharing any features (5' end clusters, 3' end clusters and any splice junction) or having 10% exon overlap with the CFC-seq models were grouped as the same genes. Finally, novel transcript models and gene models identified from this study were labeled with annotated IDs from these four databases if matches were identified.

**Comparison across assemblers**

Transcriptomes generated by SALA (final and read-filtered), IsoQuant (Default and Sensitive) and Talon (read-filtered) were analysed by SQANTI3.[7] Isoform categories were extracted and summarized. With the precision of TSS and TES derived from SALA, we further included alternative TSS (novel transcripts start from different ex5_cluster from transcript annotated by GENCODE) and alternative TES (novel transcripts end at different ex3_cluster).

```
[1] python3 sqanti3_qc.py -t 1 --report both --force_id_ignore --aligner_choice=minimap2
[query_gtf_file, eg SALA_final] [hg38_genome_fasta_file] -o [out_dir]
```

[/code_n_data/Fig3_transcript_model_analyses/assemblies_vs_reference.sh]

**Isoform switching analyses**

The transcript count derived from bambu at the replicate level of iPSC, NSC and Neuron were subjected to the IsoformSwitchAnalyzeR.[20] The object was analysed by isoformSwitchTestDEXSeq to obtain significant changes of isoform fraction. The comparison was performed between iPSC and NSC, between NSC and Neuron and between NSC and Neuron. Only the transcripts with absolute difference of fraction > 0.5 and fdr < 0.05 from any of the three combinations were included. We further filtered the transcripts with CPM > 2 from one of the two cell types. This resulted in 1,192 isoforms of GENCODE annotated genes. Genes that have been linked to diseases related to brain in Open Targets (https://platform.opentargets.org) with an overall association score > 0.2 were linked to this isoform switching results.

[/code_n_data/Fig3_transcript_model_analyses/compare_shortread_n_isoform_switch.R]

**Genomic properties incorporation to tCRE and ex5_cluster**

In order to validate the accuracy of the identified TSS clusters and tCREs, including their capability of initiating transcription and the precise position of the TSSs, we analyzed the spatial distribution of sequence properties relative to the TSS summits. For tCRE-level analysis, we utilized a non-redundant major strand subset as the representative (n = 57,597) when the tCRE overlap with another one in the opposite strand, only the strand with the higher signal intensity was retained to prevent redundant counting of genomic features. These results were shown in **Extended Figure 2e**. However, this approach missed some summits especially when defining TATA box and position specific CGI (upstream and downstream to the summit). Therefore, strand-specific ex5_clusters were also used to test the genomic properties. These ex5_clusters are strand-specific, containing 1.65 TSS clusters (**Fig. S1**) on average and only have one parental tCRE. Similar to the tCREs, ex5_clusters are non-overlapping on the same strand. In contrast to the tCREs, both ex5_clusters were included in the analyses if they overlap with the opposite strand. This will include all the ex5_cluster summits and provide annotation of regulatory elements to all of them.

For conservation and GC content profiling, we extracted sequence conservation scores (PhastCons 4-way, 17-way, 30-way and 100-way) and 5-base GC content from the UCSC genome browser (hg38). Summits of the ex5_clusters (or tCREs) were extended 5 kb up- and down-stream and intersected with the annotations. The mean values per position were extracted for the positional plot. Global conservation for each cluster was defined as the mean 17-way PhastCons score within 500 nt upstream and 500 nt downstream the summit. GC content was benchmarked against a genomic background of identical size, generated by masking all GENCODE v39 exons and identified ex5_clusters (or tCREs).

For CGI profiling, annotations of CGI were obtained from UCSC and intersected with the 10 kb window surrounding the summits. The coverage of CGI for each position was extracted. Each ex5_cluster (or tCRE) was globally defined as CGI-positive if the 1,001-nt central region exhibited 200 nt overlap with an annotated CGI that contained 40 CGI units. We classified CGI enhancers as CGIap (CGI adjacent to promoter) if their distance from promoter cCRE is ≤ 2,000 nt, and CGInap (CGI not adjacent to promoter) if the distance is > 2,000 nt. We further identified downstream CGI (dCGI) by searching 500 nt region downstream the summit with > 200 nt overlap of CGI annotation containing at least 40 CGI units.

For profiling TATA-box and Initiator (INR), their positional motifs (TBP: MA0108.3 and INR: POL002.1) were retrieved from the JASPAR database and mapped genome-wide using scanMotifGenomeWide.pl of Homer v4.11.[8] We searched a 101-nt window centered on the summit. If multiple motifs were detected, the one with the highest motif score was retained. An ex5_cluster (or tCRE) was defined as TATA-positive if a TATA box with motif score at least 3 is found between -34 to -27 nt from the summit according to the first base of the TATA motif.

In comparison of the sequence conservation among features (1D vs 2D & SE vs TE), 1001 nt regions surrounding the summits were used as non-redundant units. Overlapping units were grouped, and any group containing multiple regulatory classes (CGIap, CGInap and TATA) was excluded to maintain class purity. For regions containing multiple tCRE or ex5_clusters with the same classes, the tCRE or ex5_cluster with the highest read count was selected as the representative.

```
[1] scanMotifGenomeWide.pl MA0108.3.motif hg38 -bed -keepAll -p 10 > hg38.TATAbox.bed
[2] scanMotifGenomeWide.pl INR_oldJaspar_10100.motif hg38 -bed -keepAll -p 10 > hg38.INR.bed
```

### Bidirectionality of tCREs

Bidirectionality was acquired from SCAFE output for each tCRE. Qualified reads were used for counting the number of CTSS on both strands (**Ext_Fig. 3a**). The D was calculated as absolute difference of the forward and reverse counts divided by the sum of the forward and reverse counts. The tCRE was defined as bidirectional if $D < 0.8$, unidirectional otherwise. If the sum of the forward and reverse counts is less than 5, the directionality was set as undefined. Directionality of ex5_clusters was inherited from tCREs. [/code_n_data/Fig2_CRE_analysis/CRE_feature.R]

### Super enhancer identification

Super enhancers were predicted based on the magnitude of the H3K27ac ChIP-seq signal from various studies.[21–23] The histone mark H3K27ac was determined from iPSC, NSC and Neuron by CUT&Tag.[24] Briefly, reads were mapped to hg38 human genome using Bowtie2 (v2.2.6). After removing PCR duplicates, the bam files of the two replicates for each cell type were subjected to MACS2 (v2.1.0) for peak calling. The significant peaks ($p < 0.01$) were merged and extended into a minimum window size of 150bp. The peaks were used as the reference loci for counting the read number from the bam files including H3K27ac and IgG control using ROSE (v1.3.1).[25,26] The enhancer-like CREs were then intersected with the super enhancer regions defined from the 3 cell-types and marked as super enhancers. The number, activity and size of the SE region were shown in **Ext_Fig. 3b&c**. Super enhancer definition of ex5_clusters were inherited from tCREs.

```
[1] ROSE_main.py -g hg38 -i [significant H3K27ac peaks] -r [H3K27ac bam file] -c [IgG bam
file] -o [output folder] -s 10000 -t 2500
```

[/code_n_data/Fig2_CRE_analysis/CRE_feature.R]

### Integration of histone modification

Chromatin state results derived from H3K4me1, H3K4me3, H3K27ac, H3K27me3 CUT&Tag data and single-cell ATAC-seq from iPSC, NSC and Neuron were integrated from our another study.[24] Briefly, this model integrated CUT&Tag data (with corresponding IgG controls) and scATAC-seq data, divided as three samples (iPSC, NSC, and neurons). Both CUT&Tag and scATAC-seq data were applied as mapped reads (bam files). A 16-state model was trained using the LearnModel function. The clusters identified from ChromHMM[27] using 200 nt bins from iPSC, NSC and Neuron were manually curated and intersected with all the tCREs (n = 73,328) to identify potential promoter and enhancer. These chromatin states were transferred to the daughter ex5_clusters. This annotation was only used for two analyses: 1) investigate unclassed tCRE without

SCREEN cCRE support, and 2) eRNA length and splicing as an additional enhancer definition. Otherwise, promoter-typing utilizes SCREEN cCRE annotation.

In parallel, the H3K27ac and H3K27me3 were used to define active and repressive state of enhancers. Briefly, the bulk CUT&Tag data was processed by the ENCODE ATAC-seq pipeline (v1.7.0). Following the removal of PCR duplicates, peak calling was performed on merged replicates for each cell type using MACS2 (v2.1.0) for peak calling. The significant peaks ($p < 0.01$) were merged and extended into a minimum window size of 150 bp. These significant peaks were intersected with tCREs and ex5_clusters separately.

**Grouping of ex5_clusters, transcript models and gene models into mRNA, p_ncRNA, e_ncRNA, CTCF_ncRNA and other_ncRNA**

The ex5_clusters supported by SCAFE were divided into 4 groups according to the promoter types, which were inherited from the tCRE as promoter-like, enhancer-like, CTCF-alone with ATAC support and unclassed with ATAC support. Only the ex5_clusters producing ncRNAs (> 50% complete read count) were included while the definition of ncRNAs were described according to the CPAT results or GENCODE annotation. Additionally, ex5_clusters producing GENCODE protein-coding genes alone were grouped as a control. The ex5_clusters linked to both coding (defined by GENCODE) and non-coding (defined by GENCODE or CPAT) transcript models were excluded from the analyses. Finally, ex5_clusters linked to ncRNA transcript models belonging to protein-coding genes were also excluded. This grouping was used in most of the analyses in this study. These resulted in 8,759 ex5_clusters with mRNA outputs, 16,200 promoter-like ex5_clusters with ncRNA outputs (p_ncRNA), 28,354 enhancer-like ex5_clusters with ncRNA (e_ncRNA), 190 ATAC-supported CTCF-alone ex5_clusters with ncRNA (CTCF_ncRNA) and 4,606 ATAC-supported unclassed ex5_clusters with ncRNA (other_ncRNA). For downstream analyses where other features were involved, some extra ex5_clusters were excluded. These were described in the specific analyses. Once the ex5_clusters were included, all the reads derived from the clusters were included, independent of coding potential. These reads are considered as the RNA output of the ex5_clusters.

For grouping of transcript models, the identities of coding and non-coding and promoter types (promoter-like, enhancer-like, CTCF-alone with ATAC support and unclassed with ATAC support) were directly transferred from the finalized transcript information table. As described previously, only non-coding transcripts linked to non-coding genes were included into downstream analyses (**Ext_Fig. 4j**). From these criteria, 28,230 mRNAs, 36,723 p_ncRNAs, 64,010 e_ncRNAs, 437 CTCF_ncRNAs and 14,847 other_ncRNAs were obtained, while 49,688 transcript models were excluded. This grouping was used in summarizing if the presence of splicing and PAS correlate with transcript length, and the qualifying the libraries (**Fig. 1e**)

The grouping of gene models was used in comparing splicing efficiency and exosome sensitivity. The identities of coding and non-coding were derived from coding classes at gene level described in the previous part. The identities of promoter types were derived from the promoter type at gene level described in the previous part. If the genes were annotated as protein-coding by GENCODE, the genes were assigned as "mRNA". From these criteria, 14,274 mRNAs, 12,696 p_ncRNAs, 22,403 e_ncRNAs, 157 CTCF_ncRNAs and 3,787 other_ncRNAs were obtained. Gene level classification was used in splicing efficiency analyses.

**Transcription properties analyses**

All the transcription properties including transcript length, genomic range, number of exons, splice junction and TES, are defined at the read level and grouped into ex5_clusters and transcript models. The transcript length and exon number of transcript models and those derived from ex5_clusters were determined from the median of all the reads linked to them. This information was summarized in **Supplementary tables S10 and S11**. The ex5_clusters were grouped into mRNAs, p_ncRNAs, e_ncRNAs, CTCF_ncRNAs and other_ncRNAs as described earlier. Ex5_clusters were also grouped for directionality (1D or 2D) and being within the super enhancer (SE) cluster or not (typical enhancer, TE). This information was inherited from their linked tCREs. The e_ncRNA ex5_clusters were also grouped mutually exclusive as CGIap, CGInap, Null and TATA while p_ncRNA ex5_clusters were grouped in CGI, Null and TATA, according to as earlier described.

**Exosome sensitivity**

RNA-seq data of another clone of iPS cell[1] treated with knockdown of EXOSC3 and control was annotated by the transcriptome generated in this study using pseudo alignment of Kallisto[17] using the chromatin-bound transcriptome containing all the completely and partially detected GENCODE models, excluding ribosomal RNAs, as reference. Exosome sensitivity score was calculated at transcript and gene levels as previously described: $(TPM_{Exosome\text{-}suppressed} - TPM_{Control}) / TPM_{Exosome\text{-}suppressed}$. Transcripts that were not detectable from these short-read libraries (sum of TPM from 4 libraries < 0.1) or not detectable from both the iPSC and chromatin-bound iPSC CFC-seq libraries were excluded from the analysis. Exosome sensitivity smaller than 0 was considered as 0. To collapse exosome sensitivity at transcript and gene levels into ex5_clusters, exosome sensitivity was weighted by read counts when grouping. We performed the downstream analyses using the transcript level exosome sensitivity that one transcript only has one ex5_cluster. For single end RNA-seq with exosome-KD:

```
[1] kallisto quant -i [kallisto_index] -o [out_dir] --single -l 200 -s 50 --rf-stranded [fq_path]
```

#### RNA binding protein interaction analysis

To characterize the post-transcriptional regulatory landscape of the identified transcripts, we analyzed the occupancy of 220 RNA-binding proteins (RBPs) using the human CLIPdb from POSTAR3.[19] Binding regions were intersected with the exonic sequences of finalized transcript models (SALA Final). While 83% of exons (402544/483024) derived from promoter-like transcript models are bound by RBPs, only 26% of enhancer-derived exons (45692/162523) exhibited RBP binding. To identify specific RBP binding on eRNA exon, an enrichment test was performed for each RBP to compare proportion of peaks of a specific RBP in enhancer/promoter relative to the total RBP binding events as background. Significant enrichment of specific RBPs in enhancer exons was determined using a two-sided Fisher's exact test, with p-values adjusted for multiple testing.

#### Splicing efficiency and spliceAI

The splicing efficiency was calculated for all the observed donor sites and acceptor sites from the aligned reads, number of splice / (number of splice + number of span). The splicing efficiency of all the observed splice junctions (SJs) was obtained by number of splice / (number of splice + number of span), where the "number of span" represents the average spanning event from the corresponding donor site and acceptor site (**Fig. 5b**). The calculated splicing efficiencies were grouped at gene levels, where the SJs belonging to the reads were linked to transcript models and to gene models, and the mean splicing efficiency normalized by intron count was used.

All identified donor and acceptor splice sites were extended by ±40 nt and converted into FASTA format. These sequences containing both intron and exon were then analysed using spliceAI (v1.3.1)[28] to predict splicing potential based on the surrounding sequence composition. The prediction was done using the approach for scoring custom sequences described on the official GitHub page. Briefly, it involves padding a custom input sequence to 10,000 nucleotides. The padded sequence is converted into a one-hot encoded format suitable for model input. Five pre-trained SpliceAI models are applied to the encoded sequence, and the predictions are averaged to obtain acceptor and donor site probabilities.

```
[1] nohup python3 junction_run.py 1 all_ontCAGE.SS3J81.tsv > run1.log 2>&1 &
```

### RNA structural prediction proximal to transcript end site

To assess structural features at transcription termini, the TES with the highest read count was selected from each ex3_cluster. These were categorized as poly(A) or non-poly(A) based on the presence of canonical Polyadenylation Signals (PAS) and the poly(A) classifier. We anchored our classification thresholds using an optimal cutoff derived from a THP-1 ROC curve (**Ext_Fig. 1h**). By linking the ex5_cluster and ex3_cluster through the transcript models, the selected TESs were then grouped by ex5_clusters (mRNA, p_ncRNA, e_ncRNA, CTCF_ncRNA and other_ncRNA) and regulatory elements (CGI and TATA-box). If more than one ex5_cluster groups or regulatory element groups were found, the TESs were excluded from the analyses. The TESs were further grouped into recursive (≥ 3 counts) or non-recursive according to the raw counts.

Secondary structure propensities were predicted for sequences flanking the TES (±200 nt from the TES). We utilized RNAfold (v2.6.4) of the ViennaRNA package[29] using the partition function (-p) to calculate the Boltzmann-weighted equilibrium ensemble. This approach accounts for the structural diversity of eRNAs rather than relying solely on a single Minimum Free Energy conformation. Thermodynamic paired probability (P) was calculated per nucleotide per alignment from the base-pairing probability matrix ($P_{ij}$) provided in the PostScript dot plots. These values were summarized into relative hairpin score for each TES group (eg, non-poly(A) e_ncRNA with 3 reads) aligned by the 3' end by taking the average.

```
[1] RNAfold --noPS -p [fasta_file] > [text_file]
```

$$p(i) = \sum_{i \neq j} P_{ij}$$

To predict for a similar pattern of structural depletion for individual TES, mean paired probability of the RNA body (-100 to -6 nt of TES) and the structural valley (-5 to +14 nt of TES) were calculated. Predicted structural depletion was considered if the mean score for the RNA body from ≥ 0.55 and the depletion fold change (1 - valley / RNA body) ≥ 0.15. [/code_n_data/Fig5_TES_analyses/RNAfold_analysis.R]

### Genomic and Regulatory Characterization of TES

To investigate the epigenetic landscape of transcription termination, we analyzed the enrichment of CGI and MYC ChIP-seq data relative to TESs. CGI annotations for the hg38 genome were retrieved from the UCSC Genome Browser. MYC ChIP-seq peak regions derived from human embryonic stem cells (GSM1505809) were lifted over to hg38 and used. Using the recursive TESs, we extended the 1-nt terminal locus 5 kb up- and down-stream and intersected these windows with CGI coordinates and MYC binding regions. For each transcript category, we calculated the percentage coverage of CGI sequences to identify specific enrichment patterns.

For both MYC binding and GCI coordinates, we intersected the 1-nt recursive TES loci with MYC binding regions across four promoter-types (mRNA, p_ncRNA, e_ncRNA and other_ncRNA) to determine presence of co-localization. Termini were further stratified by their polyadenylation status (poly(A)+ vs. poly(A)-) based on our standardized classification criteria.

**eQTL and GWAS SNP locations in expanded transcriptome and their effect on splicing**

The eQTL and GWAS data were obtained from eQTL Catalogue[30] and CAUSALdb[31] respectively. For the eQTL data, a cutoff of pip >0.3 was applied to include SNPs that are putatively casual. This dataset contains 367,989 non-redundant SNP locations. For the GWAS data, credible set version 2.0 was downloaded and converted into hg38 by crossing over the rsid with the SNP 151 dataset obtained from UCSC (n = 710,172,836). The GWAS dataset contains 917,107 non-redundant SNP locations after converting to hg38. These SNP locations were intersected with the regions of the ex5_clusters and transcript models (exons). Ex5_clusters linked to at least one GENCODE transcript model were grouped as ENST, otherwise were novel. The transcript models were intersected as they were. The SNP locations mapped to the GENCODE transcript models were removed from the novel group to reveal only the additional SNP coverage from the novel entries.

The donor and acceptor sites of the finalized transcript models and all the GENCODE v39 transcript models were also subjected to the SpliceAI together with the GWAS and eQTL SNPs to identify the SNPs that affect splicing. We updated the transcript boundaries used by SpliceAI to include our new transcript model by replacing the grch38.txt file in the SpliceAI package files. After this replacement, we ran SpliceAI from the command line using the parameter -A grch38 to specify the custom gene annotation file.

After collecting the SNPs that affect the predicted splicing potential, a threshold of at least delta 0.5 for either gain or loss from donor sites or acceptor sites was applied. These filtered results were maintained in **Table S15**. For visualization in **Figure 6a-b**, we further required the genomic locations of the SNPs within 2 nt from the splice junctions and collapsed the observation at the transcript level. If a SNP affected both annotated model (ENST) and unannotated models, only the annotated one was counted as positive.

```
#Running on separate GPUs the predictions for the VCFs (thus using CUDA_VISIBLE_DEVICES)
[1] CUDA_VISIBLE_DEVICES=0 nohup python3 __main__.py -I fixed_eqtl.hg38.vcf -O spliceai_out/out_eqtl.hg38.vcf -R genome.fa -A grch38 > run0.log 2>&1 &
[2] CUDA_VISIBLE_DEVICES=1 nohup python3 __main__.py -I fixed_credible_set.hg38.vcf -O spliceai_out/out_fixed_credible_set.hg38.vcf -R genome.fa -A grch38 > run1.log 2>&1 &
```

**Enrichment of GWAS and eQTL SNPs in enhancers**

To divide these results according to the enhancer structures (CGIap, CGInap, Null and TATA), intersection was grouped at the ex5_cluster level. Exons were merged from the transcripts that are linked to the same ex5_clusters using bedtools merge. To include the most relevant SNPs, eQTL dataset was filtered for those derived from iPSC and brain related samples while GWAS dataset was filtered for traits related to brain. In order to obtain a normalized enrichment, "common_all_20180418" dataset downloaded from NIH was used as background (n = 37,302,987, unique SNP locations = 36,363,997). The eQTL and GWAS datasets were limited to this scope of SNPs (unique SNP locations = 81,782 & 35,325 respectively) The regions per ex5_cluster were intersected with the common_all dataset, the eQTL dataset and the GWAS dataset. Exons that were shared by more than one ex5_cluster were recounted. The results were normalized as the rate of hit compared to the background SNPs or as the number of SNPs per 1kb. For the rate of hit compared to background SNPs, ex5_clusters containing no background SNPs were excluded.

**Hi-C experimental and analytical procedures**

The cells (iPSC, NSC and Neuron) were cultured and harvested as described above. Two replicates of each cell type were fixed by 1% formaldehyde and Hi-C library construction was performed with the Arima-HiC+ kit and the Arima Library Prep Module (Arima Genomics) according to the manufacturer's protocols. The libraries were then subjected to Illumina NovaSeq6000 for sequencing with 150 bp paired-end mode. In order to yield a higher proportion of aligned reads, we first trim the 6 bp from the left end of each read as suggested by Arima Hi-C data analysis and then trim the 70 bp from the right end of each read using the command line "seqtk trimfq -b 6 -e 70". The remaining 75 bp reads were used for the nf-core hic pipeline v 2.1.0 (https://nf-co.re/hic/2.1.0/). Since the two replicates of the Hi-C libraries yield consistent results in a pre Hi-C processing based on the subsampled samples. The two replicates from the same cell type were pooled together for the final nf-core hic pipeline with using the following options:

```
[1] nextflow run nf-core/hic
--genome 'hg38'
--restriction_site '[^GATC,G^ANTC]'
--ligation_site '[GATCGATC,GANTGATC,GANTANTC,GATCANTC]'
--digestion 'arima'
--res_compartments '500000,250000,100000'
--tads_caller 'insulation,hicexplorer’
--bin_size '1000000,500000,100000,50000,25000,10000,5000,2500,1000'
```

The processed Hi-C data was transformed into valid genomic interaction pairs at 5kb, 10kb and 25kb resolutions. The statistical significance of these interactions was calculated using the Bioconductor package GOTHiC (version 1.40.0).[32] Intra-chromosomal (*cis*) interactions supported by at least five read pairs and a q-value (FDR) ≤ 0.01 were deemed significant for downstream analysis. The Hi-C interaction at 5-kb resolution was used in this study, where about 10% of the valid bin pairs were found significant.

#### Chromatin connectivity

To link the Hi-C data with ncRNA ex5_clusters, 5 kb bins overlapping with their summits were extracted and grouped based on promoter types (p_ncRNA, e_ncRNA, and other_ncRNA) and regulatory elements (CGIap, CGInap, Null, and TATA box), while bins containing more than one promoter type or regulatory element were excluded from the analysis. To link mRNA to Hi-C bins, only GENCODE (v39) protein-coding transcript models detectable in the final transcriptome were included. A "target" bin was considered if it overlapped with the ex5_cluster summit of these protein-coding transcripts. The number of bin pairs grouped by the above criteria was visualized in **Figure 6d**. To minimize the possibility of ncRNA ex5_clusters being missed by Hi-C due to technical differences between Hi-C and CFC-seq, only ex5_clusters showing at least one significant connection to an mRNA ex5_cluster were included. However, including ncRNA ex5_clusters with zero contacts in the analysis resulted in the same overall observation.

#### Chromatin accessibility and GC content for Hi-C bin

The chromatin accessibility of the source Hi-C bins defined in the previous section was assessed using scATAC-seq data. The scATAC-seq data were recounted into the Hi-C bins using the “CreateFragmentObject” function from the R package Signac. The resulting quantification was aggregated based on cell type. The GC content of the Hi-C bins was determined by calculating the ratio of G and C residues.

### Prediction of enhancer-gene interactions using ABC model

Enhancer-gene interactions were predicted using the Activity-by-Contact (ABC) model.[33] Analyses were performed independently for iPSC, NSC and Neuron using single-cell ATAC-seq (grouped by cell types), bulk H3K27ac CUT&Tag and Hi-C data. The ABC pipeline was executed using the Snakemake workflow under the hg38 genome assembly. Signals were normalized using the reference quantile-normalization distribution supplied with the ABC workflow (EnhancersQNormRef.K562.txt). Gene annotations were based on the hg38 CollapsedGeneBounds reference provided by the ABC package.

Instead of generating candidate regulatory elements from MACS2 peak calling, we supplied a custom catalog of tCREs identified from CFC-seq. These tCREs were merged for overlap from opposite strand. Candidate regions include both distal enhancer-like tCREs and promoter-associated tCREs while the median merged tCRE length was 501 bp. These regions were formatted as BED intervals and substituted directly into the ABC workflow at the candidate-region generation stage.

Predicted enhancer–gene interactions were extracted from the thresholded ABC output using the default score threshold of 0.02. Only interactions exceeding this threshold were considered high-confidence enhancer–gene links for downstream analyses. For comparative analyses across differentiation, ABC scores were calculated independently in iPSC, NSC and neuron samples using the same candidate regulatory element catalog. The final output was shown in **Supplementary Table S16**

```
[1]snakemake -c16 --rerun-incomplete --keep-going \
./results/iPSC/Predictions/EnhancerPredictionsAllPutative.tsv.gz \
./results/NSC/Predictions/EnhancerPredictionsAllPutative.tsv.gz \
./results/Neuron/Predictions/EnhancerPredictionsAllPutative.tsv.gz
```

### Repeat elements in ex5_clusters and transcript models

The coordinates of repeat elements in hg38 were obtained from UCSC RepeatMasker file. These repeat elements were intersected with the regions of ex5_clusters and the exons of transcript models. For ex5_clusters, a minimum overlap of 6 nt was required, and if multiple repeat elements were identified, the one with the greatest overlapping length was selected as representative. For exons, the overlap of a single feature across multiple exons of a transcript model was summed, with a minimum overlap of 200 nt required. The repeat elements intersecting at the transcript level were collapsed into the ex5_cluster level by selecting the one with the greatest overlap length. All the overlapped repeat elements that met the above criteria were retained in **Supplementary Table S17**.

#### Transcription factor motif enrichment

Summits from ex5_clusters were extended 400 nt upstream and 100 nt downstream. These regions were excluded if they overlapped and involved more than one transcript class (mRNA, p_ncRNA, e_ncRNA, other_ncRNA and CTCF_ncRNA) or more than one promoter structure (CGI, and TATA box). The ex5_cluster with stronger signal was kept if their extended regions overlap. Finally, each 501 nt region contains only one ex5_cluster. These regions (n = 39,794) were grouped by transcript class and promoter structure and subjected to SEA (Simple Enrichment Analysis) from the MEME-suite (v5.5.7)[34] to calculate the relative motif enrichment. We obtained the sequences from each region as the input sequence in fasta format and the shuffled sequences as the control sequences. The core vertebrates-non-redundant motif collection obtained from JASPAR2024 was used as the scope of transcription factors. For visualization, only motifs from the TFs with CPM >1 according to CAGE quantification, from at least one of the three cell types were included. We further selected the motifs for q value < 0.01 and enrichment > 12.

```
[1] bedtools getfasta -fi hg38.genome.fa -bed ex5_cluster.bed -fo ex5_cluster.fa
[2] sea
--p ex5_cluster.fa
--m JASPAR2024_CORE_vertebrates_non-redundant_pfms_meme.txt
--thresh 10000000
--text
--noseqs > result.txt
```

#### Integration of NFYB and TEAD4 ChIP-seq datasets

ChIP-seq data of NFYB derived from iPSC WTC11 was obtained from ENCODE (ENCSR146UIC) where the IDR threshold peaks (n = 9759, median = 390 nt) were used. ChIP-seq of TEAD4 was performed in our iPSC (WTC11) where the merged peaks from the 2 replicates were used (n = 6986, median = 437 nt). These peaks were intersected with the ex5_cluster non-overlapped regions (summit +100 / -400 nt) described from the previous section. The ex5_clusters were grouped according to the H3K27ac/me3 signals of iPSC from the CUT&Tag results, into 4 groups: Co-mark ($H3K27ac^{+}$ / $H3K27me3^{+}$), Active ($H3K27ac^{+}$ / $H3K27me3^{-}$), Repressed ($H3K27ac^{-}$ / $H3K27me3^{+}$) and Un-marked ($H3K27ac^{-}$ / $H3K27me3^{-}$). Enrichment tests were performed by Fisher's exact, using the other groups as background.

ChIP-seq data of NFYB was also used to overlay with the ex5_clusters, which were restricted to the list of non-overlapped regions and active in iPSC (TSS count > 1). The cumulative coverage was determined from distinct transcript classes and promoter structures, as in **Extended Data Figure 9g**.

### Section 5 | Code Availability

Code and limited data is available in https://github.com/fantom-prj/CFC_eRNA_analysis

SALA software package is available in https://github.com/fantom-prj/SALA

### Section 6 | Datasets used in this study

| | Accession Numbers / sources |
|---|---|
| **Core data** | |
| Bulk RNA-seq (iPSC, NSC, Neuron) | DRA019571 (DRR614936-DRR614941) |
| CFC-seq (iPSC, NSC, Neuron) | DRA019506 (DRR613094-DRR613110) |
| CFC-seq (THP-1, dTHP-1) | DRA019521 (DRR613160-DRR613175) |
| Chromatin-bound CFC-seq (iPSC) | DRA019506 (DRR613111 ,DRR613112) |
| **Supportive data** | |
| ssCAGE (iPSC, NSC, Neuron) | DRA019567 (DRR614867-DRR614872) |
| Single cell ATAC (iPSC, NSC, Neuron) | DRA019608 (DRR618513-DRR618515) |
| CUT&Tag (iPSC, NSC, Neuron; H3K27ac, H3K27me3, H3K4me1, H3K4me3, CTCF & ctrl) | DRA019568 (DRR614873-DRR614905) |
| Hi-C (iPSC, NSC, Neuron) | DRA019572 (DRR614942-DRR614953) |
| iPSC TEAD4 ChIP-seq | DRA026820 (DRR959946-DRR959949) |
| iPSC ATAC with TEAD4 KD | DRA026820 (DRR959956-DRR959961) |
| **External data** | |
| CAP trap PacBio Long-read (WTC11 iPSC) | ENCODE: ENCSR309IKK |
| uncap_deplete_PacBio Long-read (WTC11 iPSC) | ENCODE: ENCSR507JOF |
| R2C2_ONT Long-read (WTC11 iPSC) | ENCODE: ENCSR925UQZ |
| TSO_ONT Long-read (WTC11 iPSC) | ENCODE: ENCSR539ZXJ |
| dRNA_ONT Long-read (WTC11 iPSC) | ENCODE: ENCSR392BGY |
| MYC ChIP-seq (human ES) | GSM1505809 |
| NFYB ChIP-seq (WTC11 iPSC) | ENCODE: ENCSR146UIC |
| Exosome KD RNA-seq (iPSC) | DRA013847 |
| **Public Resources** | |
| eQTL Catalogue[30] | https://www.ebi.ac.uk/eqtl/ |
| GWAS from CAUSALdb[31] | http://www.mulinlab.org/causaldb/index.html |
| cCRE from SCREEN[35] | https://screen.encodeproject.org/ |
| Splice junction resources from Intropolis[36] | https://github.com/nellore/intropolis |
| Disease gene from Open Targets[37] | https://platform.opentargets.org/ |
| RNA-protein interaction from POSTAR3[19] | http://111.198.139.65/ |
| Short tandem repeat from HipSTR[13] | https://github.com/HipSTR-Tool/HipSTR |
| Functions of lncRNAs from EVLncRNAs 3.0[38] | https://www.sdklab-biophysics-dzu.net/EVLncRNAs3/#/ |

### Section 7 | Softwares used in this study

| Tools | Sources |
|---|---|
| SALA (v1.0) [this study] | https://github.com/fantom-prj/SALA |
| Tail trimmer (v1.4) [this study] | https://github.com/fantom-prj/SALA |
| Dorado (v0.2.4) | https://github.com/nanoporetech/dorado |
| Minimap2 (v2.17-r974-dirty)[4] | https://github.com/lh3/minimap2 |
| TranscriptClean (v2.0.3)[5] | https://github.com/mortazavilab/TranscriptClean |
| TALON (v5)[6] | https://github.com/mortazavilab/TALON |
| SCAFE (v1.01)[11] | https://github.com/chung-lab/SCAFE |
| STAR (v2.7.11b)[39] | https://github.com/alexdobin/STAR |
| Homer (v4.11)[8] | https://github.com/javrodriguez/HOMER |
| MEME-suite (v5.5.7)[34] | https://meme-suite.org/meme/meme-software |
| CPAT (v3.0.4)[15] | https://github.com/liguowang/cpat |
| SQANTI3 (v5.3.0)[40] | https://github.com/ConesaLab/SQANTI3 |
| IsoQuant (v3.4.1)[16] | https://github.com/ablab/IsoQuant |
| Bambu (v3.2.4)[18] | https://github.com/GoekeLab/bambu |
| Bowtie2 (v2.2.6)[41] | https://github.com/BenLangmead/bowtie2 |
| MACS2 (v2.1.0)[42] | https://github.com/jdavisturak/MACS2-2.1.1.20160309 |
| ROSE (v1.3.1)[26] | https://github.com/stjude/ROSE |
| RNAfold (v2.6.4)[43] | https://github.com/ViennaRNA/ViennaRNA |
| ChromHMM (v1.24)[27] | https://ernstlab.github.io/ChromHMM/ |
| IsoformSwitchAnalyzeR (v2.6.1)[20] | https://github.com/kvittingseerup/IsoformSwitchAnalyzeR |
| ABC-model (v.1.1.2)[33] | https://github.com/broadinstitute/ABC-Enhancer-Gene-Prediction |
| Primer-chop | https://gitlab.com/mcfrith/primer-chop |
| Paraclu[44] | https://github.com/davetang/paraclu_prep |
| Samtools (v1.11)[45] | https://www.htslib.org/ |
| Bedtools (v2.30.0)[46] | https://bedtools.readthedocs.io/en/latest/ |
| Tabix, Bgzip (v1.15.1)[47] | https://www.htslib.org/ |
| bedGraphToBigWig (v2.8)[48] | https://github.com/ENCODE-DCC/kentUtils/tree/master/src/utils |
| Bedparse (v0.2.3)[49] | https://github.com/tleonardi/bedparse |
| R (v4.3.1)[50] | https://cran.r-project.org/ |
| Perl (v5.26.2) | https://www.perl.org/get.html |
